# A Saturated Map of Common Genetic Variants Associated with Human Height from 5.4 Million Individuals of Diverse Ancestries

**DOI:** 10.1101/2022.01.07.475305

**Authors:** Loic Yengo, Sailaja Vedantam, Eirini Marouli, Julia Sidorenko, Eric Bartell, Saori Sakaue, Marielisa Graff, Anders U. Eliasen, Yunxuan Jiang, Sridharan Raghavan, Jenkai Miao, Joshua D. Arias, Ronen E. Mukamel, Cassandra N. Spracklen, Xianyong Yin, Shyh-Huei Chen, Teresa Ferreira, JI Yingjie, Tugce Karedera, Kreete Lüll, Kuang Lin, Deborah E. Malden, Carolina Medina-Gomez, Moara Machado, Amy Moore, Sina Rüeger, Tarunveer S. Ahluwalia, Masato Akiyama, Matthew A. Allison, Marcus Alvarez, Mette K. Andersen, Alireza Ani, Vivek Appadurai, Liubov Arbeeva, Seema Bhaskar, Lawrence F. Bielak, Sailalitha Bollepalli, Lori L. Bonnycastle, Jette Bork-Jensen, Jonathan P. Bradfield, Yuki Bradford, Peter S. Braund, Jennifer A. Brody, Kristoffer S. Burgdorf, Brian E. Cade, Hui Cai, Qiuyin Cai, Archie Campbell, Marisa Cañadas-Garre, Eulalia Catamo, Jin-Fang Chai, Xiaoran Chai, Li-Ching Chang, Yi-Cheng Chang, Chien-Hsiun Chen, Alessandra Chesi, Seung Hoan Choi, Ren-Hua Chung, Massimiliano Cocca, Maria Pina Concas, Christian Couture, Gabriel Cuellar-Partida, Rebecca Danning, E. Warwick Daw, Frauke Degenhard, Graciela E. Delgado, Alessandro Delitala, Ayşe Demirkan, Xuan Deng, Poornima Devineni, Alexander Dietl, Maria Dimitriou, Latchezar Dimitrov, Rajkumar Dorajoo, Arif B. Ekici, Jorgen E. Engmann, Zammy Fairhurst-Hunter, Aliki-Eleni Farmaki, Jessica D. Faul, Juan-Carlos Fernandez-Lopez, Lukas Forer, Margherita Francescatto, Sandra Freitag-Wolf, Christian Fuchsberger, Tessel E. Galesloot, Yan Gao, Zishan Gao, Frank Geller, Olga Giannakopoulou, Franco Giulianini, Anette P. Gjesing, Anuj Goel, Scott D. Gordon, Mathias Gorski, Sarah E. Graham, Jakob Grove, Xiuqing Guo, Stefan Gustafsson, Jeffrey Haessler, Thomas F. Hansen, Aki Havulinna, Simon J. Haworth, Jing He, Nancy Heard-Costa, Prashantha Hebbar, George Hindy, Yuk-Lam A. Ho, Edith Hofer, Elizabeth Holliday, Katrin Horn, Whitney E. Hornsby, Jouke-Jan Hottenga, Hongyan Huang, Jie Huang, Alicia Huerta-Chagoya, Jennifer E. Huffman, Yi-Jen Hung, Shaofeng Huo, Mi Yeong Hwang, Hiroyuki Iha, Daisuke D. Ikeda, Masato Isono, Anne U. Jackson, Susanne Jäger, Iris E. Jansen, Ingegerd Johansson, Jost B. Jonas, Anna Jonsson, Torben Jørgensen, Ioanna-Panagiota Kalafati, Masahiro Kanai, Stavroula Kanoni, Line L. Kårhus, Anuradhani Kasturiratne, Tomohiro Katsuya, Takahisa Kawaguchi, Rachel L. Kember, Katherine A. Kentistou, Han-Na Kim, Young Jin Kim, Marcus E. Kleber, Maria J. Knol, Azra Kurbasic, Marie Lauzon, Phuong Le, Rodney Lea, Jong-Young Lee, Hampton L. Leonard, Shengchao A. Li, Xiaohui Li, Xiaoyin Li, Jingjing Liang, Honghuang Lin, Shih-Yi Lin, Jun Liu, Xueping Liu, Ken Sin Lo, Jirong Long, Laura Lores-Motta, Jian’an Luan, Valeriya Lyssenko, Leo-Pekka Lyytikäinen, Anubha Mahajan, Vasiliki Mamakou, Massimo Mangino, Ani Manichaikul, Jonathan Marten, Manuel Mattheisen, Laven Mavarani, Aaron F. McDaid, Karina Meidtner, Tori L. Melendez, Josep M. Mercader, Yuri Milaneschi, Jason E. Miller, Iona Y. Millwood, Pashupati P. Mishra, Ruth E. Mitchell, Line T. Møllehave, Anna Morgan, Soeren Mucha, Matthias Munz, Masahiro Nakatochi, Christopher P. Nelson, Maria Nethander, Chu Won Nho, Aneta A. Nielsen, Ilja M. Nolte, Suraj S. Nongmaithem, Raymond Noordam, Ioanna Ntalla, Teresa Nutile, Anita Pandit, Paraskevi Christofidou, Katri Pärna, Marc Pauper, Eva R. B. Petersen, Liselotte V. Petersen, Niina Pitkänen, Ozren Polašek, Alaitz Poveda, Michael H. Preuss, Saiju Pyarajan, Laura M. Raffield, Hiromi Rakugi, Julia Ramirez, Asif Rasheed, Dennis Raven, Nigel W. Rayner, Carlos Riveros, Rebecca Rohde, Daniela Ruggiero, Sanni E. Ruotsalainen, Kathleen A. Ryan, Maria Sabater-Lleal, Richa Saxena, Markus Scholz, Anoop Sendamarai, Botong Shen, Jingchunzi Shi, Jae Hun Shin, Carlo Sidore, Xueling Sim, Colleen M. Sitlani, Roderick C. Slieker, Roelof A. J. Smit, Albert V. Smith, Jennifer A. Smith, Laura J. Smyth, Lorraine Southam, Valgerdur Steinthorsdottir, Liang Sun, Fumihiko Takeuchi, Divya Sri Priyanka Tallapragada, Kent D. Taylor, Bamidele O. Tayo, Catherine Tcheandjieu, Natalie Terzikhan, Paola Tesolin, Alexander Teumer, Elizabeth Theusch, Deborah J. Thompson, Gudmar Thorleifsson, Paul R. H. J. Timmers, Stella Trompet, Constance Turman, Simona Vaccargiu, Sander W. van der Laan, Peter J. van der Most, Jan B. van Klinken, Jessica van Setten, Shefali S. Verma, Niek Verweij, Yogasudha Veturi, Carol A. Wang, Chaolong Wang, Lihua Wang, Zhe Wang, Helen R. Warren, Wen Bin Wei, Ananda R. Wickremasinghe, Matthias Wielscher, Kerri L. Wiggins, Bendik S. Winsvold, Andrew Wong, Yang Wu, Matthias Wuttke, Rui Xia, Tian Xie, Ken Yamamoto, Jingyun Yang, Jie Yao, Hannah Young, Noha A. Yousri, Lei Yu, Lingyao Zeng, Weihua Zhang, Xinyuan Zhang, Jing-Hua Zhao, Wei Zhao, Wei Zhou, Martina E. Zimmermann, Magdalena Zoledziewska, Linda S. Adair, Hieab H. H. Adams, Carlos A. Aguilar-Salinas, Fahd Al-Mulla, Donna K. Arnett, Folkert W. Asselbergs, Bjørn Olav Åsvold, John Attia, Bernhard Banas, Stefania Bandinelli, David A. Bennett, Tobias Bergler, Dwaipayan Bharadwaj, Ginevra Biino, Hans Bisgaard, Eric Boerwinkle, Carsten A. Böger, Klaus Bønnelykke, Dorret I. Boomsma, Anders D. Børglum, Judith B. Borja, Claude Bouchard, Donald W. Bowden, Ivan Brandslund, Ben Brumpton, Julie E. Buring, Mark J. Caulfield, John C. Chambers, Giriraj R. Chandak, Stephen J. Chanock, Nish Chaturvedi, Yii-Der Ida Chen, Zhengming Chen, Ching-Yu Cheng, Ingrid E. Christophersen, Marina Ciullo, John W. Cole, Francis S. Collins, Richard S. Cooper, Miguel Cruz, Francesco Cucca, L. Adrienne Cupples, Michael J. Cutler, Scott M. Damrauer, Thomas M. Dantoft, Gert J. de Borst, Lisette C. P. G. M. de Groot, Philip L. De Jager, Dominique P. V. de Kleijn, H. Janaka de Silva, George V. Dedoussis, Anneke I. den Hollander, Shufa Du, Douglas F. Easton, Petra J. M. Elders, A. Heather Eliassen, Patrick T. Ellinor, Sölve Elmståhl, Jeanette Erdmann, Michele K. Evans, Diane Fatkin, Bjarke Feenstra, Mary F. Feitosa, Luigi Ferrucci, Ian Ford, Myriam Fornage, Andre Franke, Paul W. Franks, Barry I. Freedman, Paolo Gasparini, Christian Gieger, Giorgia Girotto, Michael E. Goddard, Yvonne M. Golightly, Clicerio Gonzalez-Villalpando, Penny Gordon-Larsen, Harald Grallert, Struan F. A. Grant, Niels Grarup, Lyn Griffiths, Leif Groop, Vilmundur Gudnason, Christopher Haiman, Hakon Hakonarson, Torben Hansen, Catharina A. Hartman, Andrew T. Hattersley, Caroline Hayward, Susan R. Heckbert, Chew-Kiat Heng, Christian Hengstenberg, Alex W. Hewitt, Haretsugu Hishigaki, Carel B. Hoyng, Paul L. Huang, Wei Huang, Steven C. Hunt, Kristian Hveem, Elina Hyppönen, William G. Iacono, Sahoko Ichihara, M. Arfan Ikram, Carmen R. Isasi, Rebecca D. Jackson, Marjo-Riitta Jarvelin, Zi-Bing Jin, Karl-Heinz Jöckel, Peter K. Joshi, Pekka Jousilahti, J. Wouter Jukema, Mika Kähönen, Yoichiro Kamatani, Kui Dong Kang, Jaakko Kaprio, Sharon L. R. Kardia, Fredrik Karpe, Norihiro Kato, Frank Kee, Thorsten Kessler, Amit V. Khera, Chiea Chuen Khor, Lambertus A. L. M. Kiemeney, Bong-Jo Kim, Eung Kwon Kim, Hyung-Lae Kim, Paulus Kirchhof, Mika Kivimaki, Woon-Puay Koh, Heikki A. Koistinen, Genovefa D. Kolovou, Jaspal S. Kooner, Charles Kooperberg, Anna Köttgen, Peter Kovacs, Adriaan Kraaijeveld, Peter Kraft, Ronald M. Krauss, Meena Kumari, Zoltan Kutalik, Markku Laakso, Leslie A. Lange, Claudia Langenberg, Lenore J. Launer, Loic Le Marchand, Hyejin Lee, Nanette R. Lee, Terho Lehtimäki, Huaixing Li, Liming Li, Wolfgang Lieb, Xu Lin, Lars Lind, Allan Linneberg, Ching-Ti Liu, Jianjun Liu, Markus Loeffler, Barry London, Steven A. Lubitz, Stephen J. Lye, David A. Mackey, Reedik Mägi, Patrik K. E. Magnusson, Gregory M. Marcus, Pedro Marques Vidal, Nicholas G. Martin, Winfried März, Fumihiko Matsuda, Robert W. McGarrah, Matt McGue, Amy Jayne McKnight, Sarah E. Medland, Dan Mellström, Andres Metspalu, Braxton D. Mitchell, Paul Mitchell, Dennis O. Mook-Kanamori, Andrew D. Morris, Lorelei A. Mucci, Patricia B. Munroe, Mike A. Nalls, Saman Nazarian, Amanda E. Nelson, Matt J. Neville, Christopher Newton-Cheh, Christopher S. Nielsen, Markus M. Nöthen, Claes Ohlsson, Albertine J. Oldehinkel, Lorena Orozco, Katja Pahkala, Päivi Pajukanta, Colin N. A. Palmer, Esteban J. Parra, Cristian Pattaro, Oluf Pedersen, Craig E. Pennell, Brenda W. J. H. Penninx, Louis Perusse, Annette Peters, Patricia A. Peyser, David J. Porteous, Danielle Posthuma, Chris Power, Peter P. Pramstaller, Michael A. Province, Qibin Qi, Jia Qu, Daniel J. Rader, Olli T. Raitakari, Sarju Ralhan, Loukianos S. Rallidis, Dabeeru C. Rao, Susan Redline, Dermot F. Reilly, Alexander P. Reiner, Sang Youl Rhee, Paul M. Ridker, Michiel Rienstra, Samuli Ripatti, Marylyn D. Ritchie, Dan M. Roden, Frits R. Rosendaal, Jerome I. Rotter, Igor Rudan, Femke Rutters, Charumathi Sabanayagam, Danish Saleheen, Veikko Salomaa, Nilesh J. Samani, Dharambir K. Sanghera, Naveed Sattar, Börge Schmidt, Helena Schmidt, Reinhold Schmidt, Matthias B. Schulze, Heribert Schunkert, Laura J. Scott, Rodney J. Scott, Peter Sever, Eric J. Shiroma, M. Benjamin Shoemaker, Xiao-Ou Shu, Eleanor M. Simonsick, Mario Sims, Jai Rup Singh, Andrew B. Singleton, Moritz F. Sinner, J. Gustav Smith, Harold Snieder, Tim D. Spector, Meir J. Stampfer, Klaus J. Stark, David P. Strachan, Leen M. t Hart, Yasuharu Tabara, Hua Tang, Jean-Claude Tardif, Thangavel A. Thanaraj, Nicholas J. Timpson, Anke Tönjes, Angelo Tremblay, Tiinamaija Tuomi, Jaakko Tuomilehto, Maria-Teresa Tusié-Luna, Andre G. Uitterlinden, Rob M. van Dam, Pim van der Harst, Nathalie Van der Velde, Cornelia M. van Duijn, Natasja van Schoor, Veronique Vitart, Uwe Völker, Peter Vollenweider, Henry Völzke, Scott Vrieze, Niels H. Wacher-Rodarte, Mark Walker, Ya Xing Wang, Nicholas J. Wareham, Richard M. Watanabe, Hugh Watkins, David R. Weir, Thomas M. Werge, Elisabeth Widen, Lynne R. Wilkens, Gonneke Willemsen, Walter C. Willett, James F. Wilson, Tien-Yin Wong, Jeong-Taek Woo, Alan F. Wright, Jer-Yuarn Wu, Huichun Xu, Chittaranjan S. Yajnik, Mitsuhiro Yokota, Jian-Min Yuan, Eleftheria Zeggini, Babette S. Zemel, Wei Zheng, Xiaofeng Zhu, Joseph M. Zmuda, Alan B. Zonderman, John-Anker Zwart, 23andMe Research Team, VA Million Veteran Program, DiscovEHR (DiscovEHR and MyCode Community Health Initiative), eMERGE (Electronic Medical Records and Genomics Network), Lifelines Cohort Study, Regeneron Genetics Center, The PRACTICAL Consortium, Understanding Society Scientific Group, Daniel I. Chasman, Yoon Shin Cho, Iris M. Heid, Mark I. McCarthy, Maggie C. Y. Ng, Christopher J. O’Donnell, Fernando Rivadeneira, Unnur Thorsteinsdottir, Yan V. Sun, E. Shyong Tai, Michael Boehnke, Panos Deloukas, Anne E. Justice, Cecilia M. Lindgren, Ruth J. F. Loos, Karen L. Mohlke, Kari E. North, Kari Stefansson, Robin G. Walters, Thomas W. Winkler, Kristin L. Young, Po-Ru Loh, Jian Yang, Tõnu Esko, Themistocles L. Assimes, Adam Auton, Goncalo R. Abecasis, Cristen J. Willer, Adam E. Locke, Sonja I. Berndt, Guillaume Lettre, Timothy M. Frayling, Yukinori Okada, Andrew R. Wood, Peter M. Visscher, Joel N. Hirschhorn

## Abstract

Common SNPs are predicted to collectively explain 40-50% of phenotypic variation in human height, but identifying the specific variants and associated regions requires huge sample sizes. Here we show, using GWAS data from 5.4 million individuals of diverse ancestries, that 12,111 independent SNPs that are significantly associated with height account for nearly all of the common SNP-based heritability. These SNPs are clustered within 7,209 non-overlapping genomic segments with a median size of ~90 kb, covering ~21% of the genome. The density of independent associations varies across the genome and the regions of elevated density are enriched for biologically relevant genes. In out-of-sample estimation and prediction, the 12,111 SNPs account for 40% of phenotypic variance in European ancestry populations but only ~10%-20% in other ancestries. Effect sizes, associated regions, and gene prioritization are similar across ancestries, indicating that reduced prediction accuracy is likely explained by linkage disequilibrium and allele frequency differences within associated regions. Finally, we show that the relevant biological pathways are detectable with smaller sample sizes than needed to implicate causal genes and variants. Overall, this study, the largest GWAS to date, provides an unprecedented saturated map of specific genomic regions containing the vast majority of common height-associated variants.

## INTRODUCTION

Since 2007, genome-wide association studies (GWAS) have identified thousands of associations between common single nucleotide polymorphisms (SNPs) and height, primarily using studies of European ancestry. The largest GWAS published to date for adult height focussed on common variation and reported up to 3,290 independent associations in 712 loci using a sample size of up to 700,000 individuals.^1^ To date, adult height, which is highly heritable and easily measured, has provided a larger number of common genetic associations than any other human phenotype. In addition, a large collection of genes has been implicated in disorders of skeletal growth, and these are enriched in loci mapped by GWAS of height in the normal range. These features make height an attractive model trait for assessing the role of common genetic variation in defining the genetic and biological architecture of polygenic human phenotypes.

As available sample sizes continue to increase for GWAS of common variants, it becomes important to consider whether these larger samples can “saturate” or nearly completely catalogue the information that can be derived from GWAS. This question of completeness can take several forms, including prediction accuracy compared with heritability attributable to common variation, the mapping of associated genomic regions that account for this heritability, and whether increasing sample sizes continue to provide additional information about the identity of prioritised genes and gene sets. Furthermore, because most GWAS continue to be performed largely in populations of European ancestry, it is important to address these questions of completeness in the context of multiple ancestries. Finally, some have proposed that, when sample sizes become sufficiently large, effectively every gene and genomic region will be implicated by GWAS, rather than implicating specific subsets of genes and biological pathways.^2^

Using data from 5,380,080 individuals, we set out to map common genetic associations with adult height, using variants catalogued in the HapMap 3 project (HM3), and to assess the saturation of this map with respect to variants, genomic regions, and likely causal genes and gene sets. We identify significant variants, explore signal density across the genome, perform out-of-sample estimation and prediction analyses within European and non-European ancestry studies, and prioritise genes and gene sets as likely mediators of the effects on height. We show that this set of common variants reaches predicted limits for prediction accuracy within European-ancestry populations and largely saturates both the genomic regions associated with height and broad categories of likely relevant gene sets; future work remains to extend prediction accuracy to non-European ancestries and to more definitively connect associated regions with individual likely causal genes and variants.

## RESULTS

An overview of our study design and analysis strategy is illustrated in Suppl. Fig. 1.

### Multi-ancestry GWAS meta-analysis identifies 12,111 height-associated SNPs

We performed genetic analysis of up to 5,380,080 individuals from 281 studies from the GIANT consortium and 23andMe, Inc. including 4,080,687 participants of predominantly European ancestries (75.8% of total sample), 472,730 participants with predominantly East-Asian ancestries (8.8%), 455,180 participants of Hispanic ethnicity with typically admixed ancestries (8.5%), 293,593 participants of predominantly African ancestries, mostly African-Americans with admixed African and European ancestries (5.5%) and 77,890 participants of predominantly South-Asian ancestries (1.4%). We refer to these five groups of participants/cohorts by the shorthand EUR, EAS, HIS, AFR, and SAS, respectively, yet recognising that these commonly used groupings oversimplify the actual genetic diversity among participants. Cohort-specific information is provided in Suppl. Tables 1 – 3. We tested the association between standing height and 1,385,132 autosomal bi-allelic SNPs from the HM3 tagging panel^3^, which contains >1,095,888 SNPs with a minor allele frequency (MAF) >1% in each of the five ancestral groups included in our meta-analysis. Suppl. Fig. 2 shows the frequency distribution of HM3 SNPs across all five groups of cohorts.

We first performed individual meta-analyses in each of the five groups of cohorts. We identified 9863, 1888, 918, 493 and 69 quasi-independent genome-wide significant (GWS; *P*<5×10^−8^) SNPs in the EUR, HIS, EAS, AFR and SAS groups, respectively (Table 1; Suppl. Tables 4 – 8). Quasi-independent associations were obtained after performing approximate conditional and joint multiple-SNP (COJO) analyses,^4^ as implemented in GCTA^5^ (**Suppl. Methods**). Previous studies have shown that confounding due to population stratification may remain uncorrected in large EUR GWAS meta-analyses.^6,7^ Therefore, we specifically investigated confounding effects in our EUR GWAS and found no evidence that these GWAS results are driven by population stratification (**Suppl. Note 1**, Suppl. Fig. 3).

**Table 1.** Summary of results from within-ancestry and trans-ancestry GWAS meta-analyses. N denotes the sample size for each SNP. GWS: Genome-Wide Significant (*P*<5×10^−8^). COJO SNPs: near independent GWS SNPs identified using an approximate conditional and Joint analysis implemented in the GCTA software. *P*_GWAS_ : P-value from marginal association test. GWS loci were defined as genomic regions centred around each GWS SNP and including all SNPs within 35 kb on each side of the lead GWS SNP. Overlapping GWS loci were merged so that the number and cumulative length of GWS loci are calculated on non-overlapping GWS loci. Percentage of the genome covered was calculated by dividing the cumulative of GWS loci by 3,039 Mb, i.e. the approximated length of the human genome.

| Cohort Ancestry/Ethnic Group | Number of studies | Max N (Mean N) | Number of GWS COJO SNPs ($P_{\text{GWAS}} < 5 \times 10^{-8}$) | Number of GWS loci (35 kb) | Cumulative length of non-overlapping GWS loci (% genome) |
| --- | --- | --- | --- | --- | --- |
| European (EUR) | 173 | 4,080,687 (3,612,229) | 9,863 (8,382) | 6,386 | 552.5 Mb (18.4%) |
| East-Asian (EAS) | 56 | 472,730 (320,570) | 918 (807) | 821 | 60.5 Mb (2.0%) |
| Hispanic (HIS) | 11 | 455,180 (431,645) | 1,888 (1,332) | 1,599 | 121.4 Mb (4.0%) |
| African (AFR) | 29 | 293,593 (222,981) | 493 (417) | 436 | 32.5 Mb (1.1%) |
| South Asian (SAS) | 12 | 77,890 (59,420) | 69 (65) | 66 | 4.7 Mb (0.2%) |
| Trans-ancestry meta-analysis (META<sub>FE</sub>) | 281 | 5,314,291* (4,611,160) | 12,111 (9,920) | 7,209 | 647.5 Mb (21.6%) |

\*The number of individuals in the trans-ancestry meta-analysis (N=5,314,291) is smaller than the sum of ancestry group specific meta-analyses (N=5,380,080) because of variation in per-SNP sample sizes for SNPs included in the final analysis.

To compare results across the five groups of cohorts, we examined the genetic and physical colocalization between SNPs identified in the largest group (EUR) with those found in the non-EUR groups. We found that over 83% of GWS SNPs detected in non-EUR are in strong linkage disequilibrium 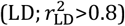 with at least one variant reaching marginal genome-wide significance in EUR (Suppl. Tables 5 – 8) and over 87% of associations detected in non-EUR meta-analyses fall within 100 kb of at least one GWS SNP identified in EUR (Suppl. Fig. 4a). In contrast, a randomly sampled HM3 SNP falls within 100 kb of a EUR GWS SNP only about 68% of the time (standard error; S.E.=0.5% over 10,000 draws). Next, we quantified the cross-ancestry correlation of allele substitution effects (*ρ*_*b*_) at GWS SNPs for all pairs of ancestry groups. We estimated *ρ*_*b*_ using five sets of GWS SNPs identified in each of ancestry group. After correction for winner’s curse,^8,9^ we found *ρ*_*b*_ to range between 0.64 and 0.99 across all pairs of ancestry groups and all sets of GWS SNPs (Suppl. Fig. 5 – 9). Thus, the observed GWS height associations are substantially shared across major ancestral groups, consistent with previous studies based on smaller sample sizes.^10,11^

To find signals that are specific to certain groups, we tested if any individual SNPs detected in non-EUR GWAS are conditionally independent of signals detected in EUR GWAS. We fitted an approximate joint model that includes GWS SNPs identified in EUR and non-EUR, using LD reference panels specific to each ancestry group. After excluding SNPs in strong LD (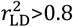 in either ancestry group), we found that 2, 19, 49 and 143 of the GWS SNPs detected in SAS, AFR, EAS and HIS GWAS respectively are conditionally independent of GWS SNPs identified in EUR GWAS (Suppl. Table 9). On average these conditionally independent SNPs have a larger MAF and effect size in non-EUR than in EUR cohorts, which may have contributed to increased statistical power of detection. The largest frequency difference relative to EUR was observed for rs2463169 (height-increasing G allele frequency: 23% in AFR vs. 84% in EUR) within the intron of *PAWR*, which codes for the prostate apoptosis response-4 protein. Interestingly, rs2463169 is located within the 12q21.2 locus, where a strong signal of positive selection in West-African Yoruba populations was previously reported.^12^ The estimated effect at rs2463169 is *β* ~0.034 standard deviation (SD) per G allele in AFR vs. *β*~-0.002 SD/G allele in EUR and the p-value of marginal association in EUR is *P*_*EUR*_ =0.08, suggesting either a true difference in effect size or nearby causal variant(s) with differing LD to rs2463169.

Given that our results demonstrate a strong genetic overlap of GWAS signals across ancestries, we performed a fixed-effect meta-analysis of all five ancestry groups to maximise statistical power for discovering associations due to shared causal variants. The mean Cochran’s heterogeneity Q-statistic is ~34% across SNPs, which indicates moderate heterogeneity of SNP effects between ancestries. The mean chi-square association statistic in our fixed effect meta-analysis (hereafter referred to as META_FE_) is ~36, and ~18% of all HM3 SNPs are marginally GWS. Moreover, we found allele frequencies in our META_FE_ to be very similar to that of EUR (mean F_ST_ across SNPs between EUR and META_FE_ is ~0.001), as expected because our META_FE_ consists of >75% EUR participants and ~14% participants with admixed European and non-European ancestries (i.e. HIS and AFR). To further assess if LD in our META_FE_ could be reasonably approximated by the LD from EUR, we performed LD score regression analysis of our META_FE_ using LD scores estimated in EUR. In this analysis, we focused on the attenuation ratio statistic (R_LDSC-EUR_), for which values >20% classically indicate strong LD inconsistencies between a given reference and GWAS summary statistics. For example, using EUR LD scores in the GWAS of HIS, which is the non-EUR group genetically closest to EUR (F_ST_~0.02), yields an estimated R_LDSC-EUR_ of ~25% (S.E. 1.8%), consistent with strong LD differences between HIS and EUR. By contrast, in our META_FE_, we found an estimated R_LDSC-EUR_ of ~4.5% (S.E. 0.8%), which is significantly lower than 20% and also not statistically different from 3.8% (S.E. 0.8%) in our EUR meta-analysis. Altogether, our LD score regression analyses suggest that LD in our META_FE_ can be reasonably approximated by LD from EUR.

We therefore proceeded to identify quasi-independent GWS SNPs from the multi-ancestry meta-analysis by performing a COJO analysis of our META_FE,_ using genotypes from ~350,000 unrelated EUR participants of the UK Biobank (UKB) as an LD reference. We identified 12,111 quasi-independent GWS SNPs, including 9,920 (82%) primary signals with a GWS marginal effect and 2,191 secondary signals that only reached GWS in a joint regression model (Suppl. Table 10). Of the GWS SNPs obtained from the non-EUR meta-analyses above that were conditionally independent of the EUR GWS SNPs, 0/2 in SAS, 5/19 in AFR, 27/49 in EAS, and 39/143 in HIS remained statistically significant in our META_FE_ (Suppl. Table 9), meaning that a small number of additional signals were only identified in the ancestry-specific analyses.

We next sought replication of the 12,111 META_FE_ signals using GWAS data from 49,160 participants of the Estonian Biobank (EBB). We first re-assessed the consistency of allele frequencies between our META_FE_ and the EBB set. We found a correlation of allele frequencies of ~0.98 between the two datasets and a mean F_ST_ across SNPs of ~0.005, similar to estimates obtained between populations from the same continent. Of the 12,111 GWS SNPs identified through our COJO analysis, 11,847 were available in the EBB dataset, 97% of which (11,529) have MAF>1% (Suppl. Table 10). Given the large difference in sample size between our discovery and replication samples, direct statistical replication of individual associations at GWS is not achievable for most SNPs identified (Suppl. Fig. 10a). Instead, we assessed the correlation of SNP effects between our discovery and replication GWAS as an overall metric of replicability.^1,13^ Over the 11,529/11,847 SNPs with a MAF>1% in the EBB, we found a correlation of marginal SNP effects of *ρ*_*b*_=0.93 (jackknife standard error; S.E. 0.01) and a correlation of conditional SNP effects using the same LD reference panel of *ρ*_*b*_=0.80 (S.E. 0.03; Suppl. Fig. 11). Although we had limited power to replicate associations with 238 GWS variants that are rare in the EBB (MAF<1%), we found, consistent with expectations (**Suppl. Methods**; Suppl. Fig. 10b), that 60% of them have a marginal SNP effect that is sign-consistent with that from our discovery GWAS (Fisher exact test; *P*=0.001). The proportion of sign-consistent SNP effects was >75% (Fisher exact test; *P*<10^−50^) for variants with a MAF>1%, also consistent with expectations (Suppl. Fig. 10b). Altogether, our analyses demonstrate the robustness of our findings and show their replicability in an independent sample.

### Genomic distribution of height-associated SNPs

To examine signal density among the 12,111 GWS SNPs detected in our META_FE_, we defined a measure of local density of association signals for each GWS SNP based on the number of additional independent associations within 100 kb (Suppl. Fig. 12). We observed that 69% of GWS SNPs shared their location with another associated, conditionally independent, GWS SNP (Fig. 1). The mean signal density across the entire genome is 2.0 (LOCO-S.E. = 0.14), consistent with a non-random genomic distribution of GWS SNPs. Next we evaluated signal density around 462 autosomal genes curated from the Online Mendelian Inheritance in Man (OMIM) database^14^ as harbouring pathogenic mutations causing syndromes of abnormal skeletal growth (“OMIM genes”; **Suppl. Methods**; Suppl. Table 11). We found that a high density of height-associated SNPs is significantly correlated with the presence of an OMIM gene nearby (Enrichment fold of OMIM gene when density >1: 2.5×; *P*<0.001; **Suppl. Methods**, Suppl. Fig. 13a).^15,16^ Interestingly, the enrichment of OMIM genes almost linearly increases with the density of height-associated SNPs (Suppl. Fig. 13b). Thus, these 12,111 GWS SNPs nonrandomly cluster near each other and also near known skeletal growth genes.

**Fig. 1.**
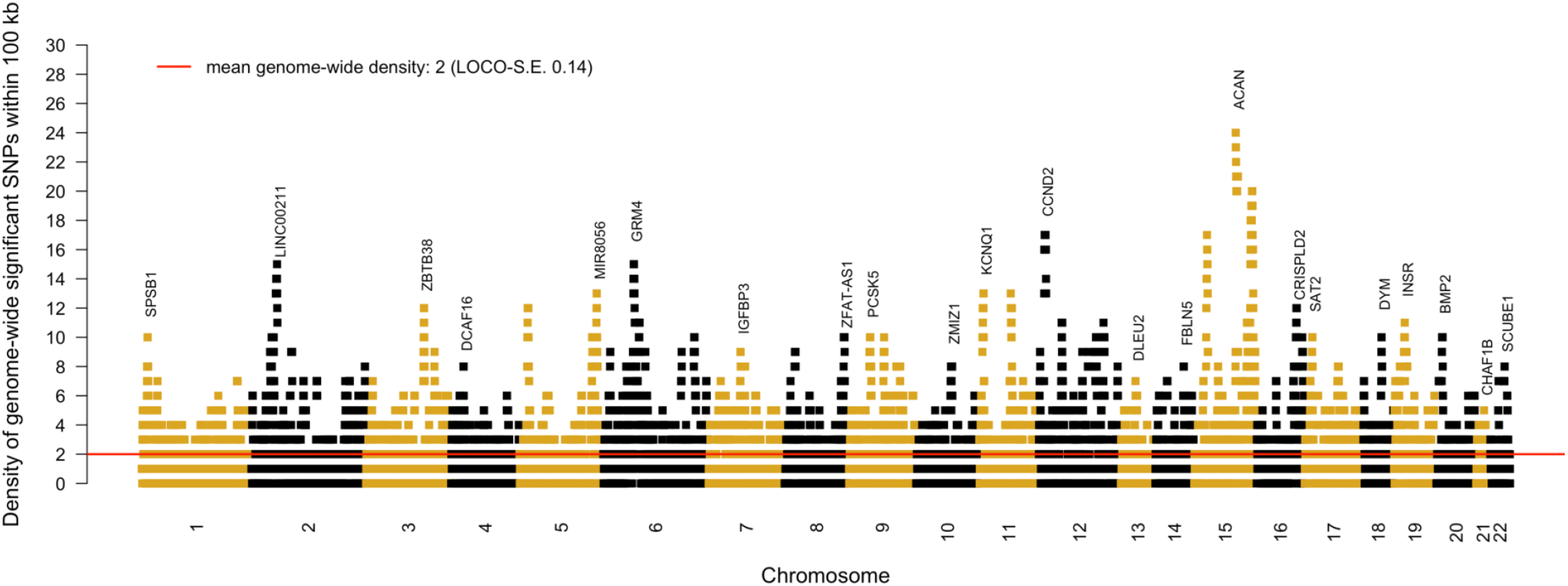
Brisbane plot showing the genomic density of independent genetic associations with height. Each dot represents one of the 12,111 quasi-independent genome-wide significant (GWS; *P*<5×10^−8^) height-associated SNPs identified using approximate conditional and joint multiple-SNP (COJO) analyses of our trans-ancestry GWAS meta-analysis. Density was calculated for each associated SNP as the number of other independent associations within 100 kb. A density of 1 means that a GWS COJO SNP share its location with another independent GWS COJO SNP within <100 kb. The average signal density across the genome is 2 (standard error; S.E. 0.14). S.E. were calculated using a Leave-One-Chromosome-Out jackknife approach (LOCO-S.E.). Sub-significant SNPs are not represented on the figure.

The largest density of conditionally independent associations was observed on chromosome 15 near *ACAN*, a gene mutated in short stature and skeletal dysplasia syndromes, where 25 GWS SNPs co-localise within 100 kb of one another (Fig. 1; Suppl. Fig. 14). We show in **Suppl. Note 2** and Suppl. Figs. 14-15, using haplotype- and simulation-based analyses, that a multiplicity of independent causal variants is the most likely explanation of this observation. Interestingly, we also found that signal density is partially explained by the presence of a recently identified^17,18^ height-associated variable-number-of-tandem-repeat (VNTR) polymorphism at this locus (**Suppl. Note 2**). In fact, the 25 independent GWS SNPs clustered within 100 kb of rs4932198 explain >40% of the VNTR length variation in multiple ancestries (Suppl. Fig. 15e) and an additional ~0.24% (P=8.7 × 10^−55^) phenotypic variance in EUR above what is explained by the VNTR alone (Suppl. Fig. 15f). Altogether, our conclusion is consistent with prior evidence of multiple types of common variation influencing height through *ACAN* gene function, involving multiple enhancers,^19^ missense variants^20^ and tandem repeat polymorphisms.^17,18^

### Variance explained by SNPs within identified loci

To quantify the proportion of height variance explained by GWS SNPs identified in our META_FE_, we stratified all HM3 SNPs into two groups: SNPs in the close vicinity of GWS SNPs, hereafter denoted GWS loci, and all remaining SNPs. We defined GWS loci as non-overlapping genomic segments containing at least 1 GWS SNP, such that GWS SNPs in adjacent loci are >2×35 kb away from each other (i.e. 35 kb window on each side). We chose a 35 kb threshold based on findings from Wu et al.^21^ who previously showed that causal common variants are located within 35 kb of GWS SNPs with >80% probability. Accordingly, we grouped the 12,111 GWS SNPs identified in our META_FE_ into 7,209 non-overlapping loci (Suppl. Table 12) with lengths ranging from 70 kb (for loci containing only 1 signal) to 711 kb (for loci containing up to 25 signals). The average length of GWS loci is ~90 kb (SD 46 kb). The cumulative length of GWS loci represent ~647 Mb, or ~21% of the genome (assuming a genome length of ~3039 Mb).^22^

To estimate what fraction of heritability is explained by common variants within the 21% of the genome overlapping GWS loci, we calculated two genomic relationship matrices (GRMs), one for SNPs within these loci and one for SNPs outside these loci, and then used both matrices to estimate a stratified SNP-based heritability 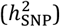 of height in 8 independent samples of all five population groups represented in our META_FE_ (Fig. 2; **Suppl. Methods**). Altogether, our stratified estimation of SNP-based heritability shows that SNPs within these 7,209 GWS loci explain ~100% of 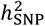 in EUR and >90% of 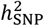 across all non-EUR groups, despite being drawn from less than a quarter of the genome (Fig. 2). We also varied the window size used to define GWS loci and found that 35 kb was the smallest window size for which this level of saturation of SNP-based heritability could be achieved (Suppl. Fig. 16).

**Fig. 2.**
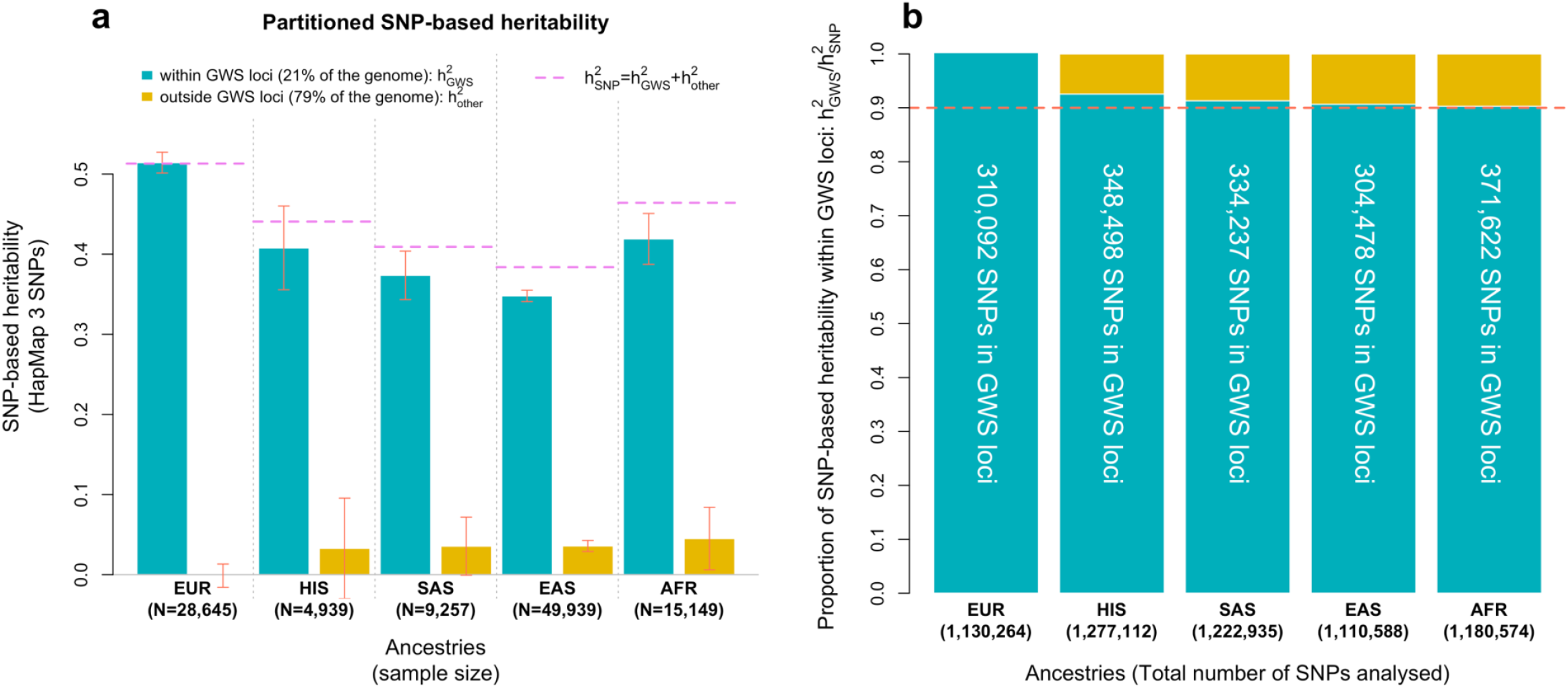
Variance of height explained by HapMap 3 SNP within genome-wide significant (GWS) loci. **Panel a** shows stratified SNP-based heritability 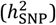 estimates obtained after partitioning the genome into SNPs within 35 kb of a GWS SNP (“GWS loci” label) vs. SNPs >35 kb away from any GWS SNP. Analyses were performed in samples of five different ancestry/ethnic groups: European (EUR: meta-analysis of UK Biobank (UKB) + Lifelines study), African (AFR: meta-analysis of UKB + PAGE study), East-Asian (EAS: meta-analysis of UKB + China Kadoorie Biobank), South-Asian (SAS: UKB) and Hispanic group (HIS: PAGE). **Panel b** shows that >90% of 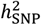 in all ancestries is explained by SNPs within GWS loci identified in this study. The cumulative length of non-overlapping GWS loci is ~647 Mb, i.e. ~21% of the genome assuming a genome length of ~3039 Mb.^22^ The proportion of HapMap 3 SNPs in GWS loci is ~27%.

To further assess the robustness of this key result, we tested if the 7,209 height-associated GWS loci are systematically enriched for trait-heritability. We chose body mass index (BMI) as a control trait given its small genetic correlation with height (*r*_g_=-0.1, ref.^23^) and found no significant enrichment of SNP-based heritability for BMI within height-associated GWS loci (Suppl. Fig. 17). Furthermore, we repeated our analysis using a random set of SNPs with similar EUR MAF and LD scores as the 12,111 height-associated GWS SNPs. We found this control set of SNPs to explain only ~27% of 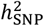 for height, consistent with the proportion of SNPs within the loci defined by this random set of SNPs (Suppl. Figs. 16 - 17). Finally, we extended our stratified estimation of SNP-based heritability to all well-imputed common SNPs (i.e. beyond the HM3 panel) and found, consistently across population groups, that although more genetic variance can be explained by common SNPs not included in the HM3 panel, all information remains concentrated within these 7,209 GWS loci (Suppl. Fig. 18). Thus, with this large GWAS, nearly all of the variability in height that is attributable to common genetic variants can be mapped to regions comprising ~21% of the genome.

### Out-of-sample prediction accuracy

We quantified the accuracy of polygenic scores (PGS) for height based on GWS SNPs in 61,095 unrelated individuals from 3 studies, including 33,001 participants of the UKB who were not included in our discovery GWAS (i.e. 14,587 EUR; 9,257 SAS; 6,911 AFR and 2,246 EAS; **Suppl. Methods**), 14,058 EUR participants from the Lifelines cohort study; and 8,238 HIS and 5,798 AFR participants from the PAGE study. Prediction accuracy 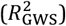 was defined as the squared correlation between the PGS and actual height (corrected for mean and variance sex differences and 20 genotypic principal components). We found that PGS based on 12,111 GWS SNPs from our META_FE_ systematically outperformed those based on GWS identified in ancestry-specific meta-analyses (Fig. 3a). The only exception was in EUR where both PGS performed equally. The largest prediction accuracy was observed in EUR participants 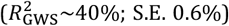 and the smallest one in AFR participants from the UKB 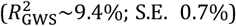. Note that the difference in 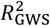 between the EUR and AFR ancestry cohorts is expected because of the over-representation of EUR in our META_FE_ and consistent with a relative accuracy (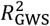 in AFR)/(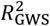 in EUR) of ~25% previously reported.^24^ Nevertheless, we found the accuracy of PGS based on GWS from our multi-ancestry META_FE_ to be consistently larger than that of PGS based on GWS SNPs from a EUR GWAS (Fig. 3a). The largest improvement was observed in AFR, where the meta-analysed accuracy in AFR participants of UKB and PAGE was increased from 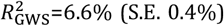 to 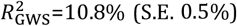, i.e. almost a ~1.6-fold improvement. This observation is partly explained by the increased statistical power but also by the refined estimation of SNP effects due to the inclusion of shorter and ancestry-specific LD blocks in AFR cohorts.

**Fig. 3.**
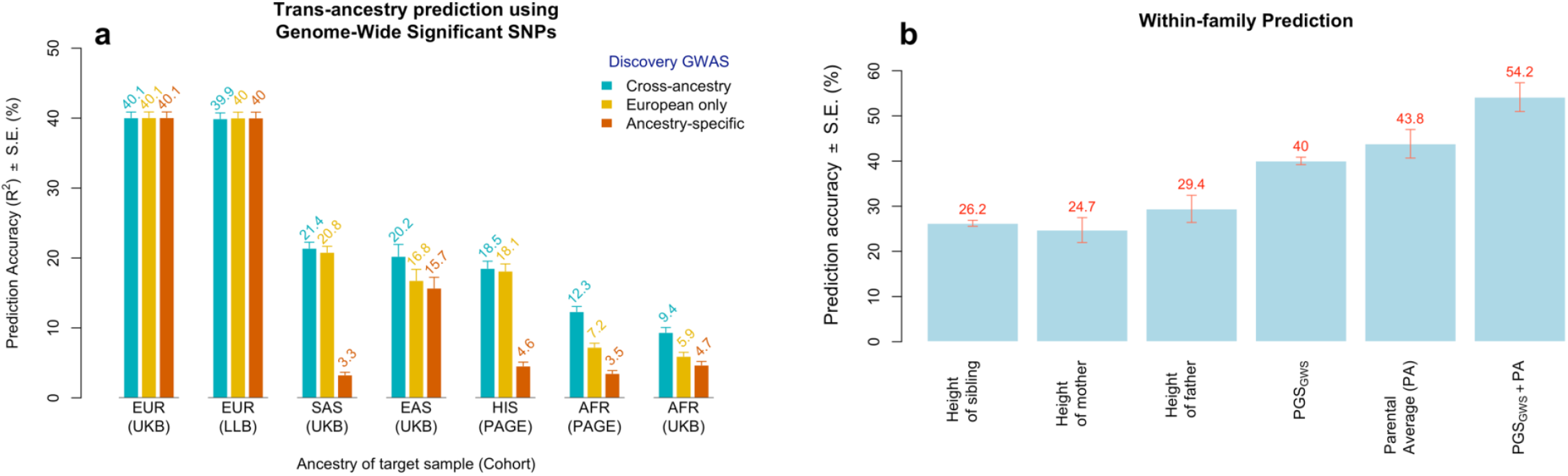
Accuracy of a polygenic predictors of height (PGS) within-family and across ancestries. Prediction accuracy (*R*^2^) was measured as the squared correlation between PGS and actual height adjusted for age, sex and 10 genetic principal components. **Panel a** shows the accuracy of PGSs assessed in participants of 5 different ancestry groups: European (EUR; N=14,587) from the UK Biobank (UKB) and the Lifelines Biobank (LLB; N=14,058) cohorts, South-Asian (SAS; N=9,257) from UKB, East-Asian (EAS; N=2,246) from UKB, Hispanic (HIS; N=8,238) from the PAGE study and admixed African (AFR) from UKB (N=6,911) and PAGE (N=5,798). PGSs used for prediction, in **Panel a**, are based on genome-wide significant (GWS) SNPs identified in (1) cross-ancestry meta-analysis (green bar), (2) EUR meta-analysis (yellow bar) and (3) ancestry-specific meta-analyses (red bars). **Panel b** shows the squared correlation of height between first-degree relatives of EUR participants in UKB and the accuracy of a predictor combining PGS (denoted, PGS_GWS_, as based on GWS SNPs) and familial information. PGS_GWS_ accuracy shown in **Panel b** is the average accuracy in EUR participants from UKB and LLB from **Panel a**. Sibling correlation was calculated in 17,492 independent EUR sibling pairs from the UKB and parent-offspring correlations in 981 EUR unrelated trios (i.e. two parents and 1 child) from the UKB.

Furthermore, we sought to evaluate the prediction accuracy of PGS relative to that of familial information as well as the potential improvement in accuracy gained from combining both sources of information. We analysed 981 unrelated EUR trios (i.e. two parents and one offspring) and 17,492 independent EUR sibling pairs from the UKB, who were excluded from our META_FE_. We found that height of any first-degree relative yields a prediction accuracy between 25% and 30% (Fig. 3b). Moreover, the accuracy of the parental average is ~44% (S.E. 3.2%), which is larger but not significantly different from 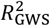 in EUR. In addition, we found that a linear combination of the average height of parents and of the offspring’s PGS yields an unprecedented accuracy of 54% (S.E. 3.2%). This observation reflects the fact that PGS can explain within-family differences between siblings, while average parental height cannot. To show this empirically, we estimate that our PGS based on GWS SNPs explain ~33% (S.E. 0.7%) of height variance between siblings (**Suppl. Methods**). Finally, we demonstrate that the optimal weighting between parental average and PGS can be predicted theoretically as function of 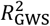, the full narrow sense heritability and the phenotypic correlation between spouses (**Suppl. Note 3**, Suppl. Fig. 19).

In summary, the estimation of variance explained and prediction analyses in European-ancestry samples show that the set of 12,111 GWS SNPs account for nearly all of 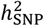 and that combining SNP-based PGS with family history significantly improves prediction accuracy. In contrast, both estimation and prediction results show clear attenuation in samples with non-European ancestry, consistent with previous studies.^24–27^

### Relationship between GWAS discoveries, sample size and ancestry diversity

Our large study offers a unique opportunity to empirically quantify how increasing GWAS sample sizes and ancestry diversity affects discovery of variants, genes and biological pathways. To address this question, we re-analysed 3 previously published GWAS of height^1,15,16^ and also down-sampled our meta-analysis into 4 subsets (including our EUR and META_FE_ GWAS). Altogether we analysed 7 GWAS with a sample size increasing from ~0.13 M up to ~5.3 M individuals (Table 2).

**Table 2.** Overview of 5 European ancestry GWAS re-analysed in our study to quantify the relationship between sample size and discovery. Summary statistics from the 3 published GWAS were imputed using the SSIMP software to maximise coverage of HapMap 3 SNPs (**Suppl. Methods**). GWS loci are defined as in the legend of Table 1.

| Down-sampled GWAS | Max N<br>(Mean N) | Number of GWS<br>COJO SNPs | % of the genome<br>covered by GWS<br>loci (35 kb) |
| --- | --- | --- | --- |
| Lango-Allen et al.<sup>15*</sup> | 130,010<br>(128,942) | 240 | 0.5% |
| Wood et al.<sup>16</sup> | 241,724<br>(239,227) | 633 | 1.4% |
| Yengo et al.<sup>1</sup> | 695,648<br>(688,927) | 2,794 | 5.8% |
| GIANT-EUR<br>(no 23andMe) | 1,632,839<br>(1,502,499) | 4,867 | 9.7% |
| 23andMe-EUR | 2,502,262<br>(2,498,336) | 7,020 | 13.6% |

\*Summary-statistics from the Lango-Allen et al. study, initially over-corrected for population stratification using a double genomic control correction, were re-inflated such that the LD score regression intercept estimated from re- inflated test statistics equals 1.

For each GWAS, we quantified 8 metrics grouped into 4 *variant*- and *locus*-based metrics (number of GWS SNPs, number of GWS loci, prediction accuracy 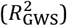 of PGS based on GWS SNPs, the proportion of the genome covered by GWS loci), a *functional annotation*-based metric (enrichment statistics from stratified LDSC^28,29^), 2 *gene*-based metrics (number of genes prioritised by Summary data based Mendelian Randomization^30^ (SMR; **Suppl. Methods**), proximity of variants with OMIM genes), and a *gene-set*-based metric (enrichment within clusters of gene sets/pathways). Overall, we found different patterns for the relationship between those metrics and GWAS sample size and ancestry composition, consistent with varying degrees of saturation achieved at different sample sizes.

We observed the strongest saturation for the *gene-set* and *functional annotation* metrics, which capture how well general biological functions can be inferred from GWAS results using currently available computational methods. Using two popular gene set prioritisation methods (DEPICT^31^ and MAGMA^32^), we found that the same broad clusters of related gene sets (including most of the clusters enriched for OMIM genes) are prioritised at all GWAS sample sizes (Suppl. Figs. 20-21; Suppl. Tables 13 – 15; **Suppl. Note 4**). Similarly, stratified LDSC estimates of heritability enrichment within 97 functional annotations also remain stable across the range of sample sizes (Suppl. Fig. 22). Overall, we found no significant improvement for all these higher-level metrics from adding non-EUR samples to our analyses. The latter observation is consistent with other analyses demonstrating that GWAS expectedly implicate similar biology across major ancestral groups (**Suppl. Note 4**; Suppl. Fig. 23).

For the *gene*-level metric, the excess in the number of OMIM genes that are proximate to a GWS SNP (compared with matched sets of random genes) plateaus at sample sizes of N>1.5M; while the relative enrichment of GWS SNPs near OMIM genes first decreases with sample size, then plateaus when N>1.5M (Suppl. Figs. 24a-c). Interestingly, the decrease observed for N<1.5M reflects the preferential localization of larger effect variants (those identified with smaller sample sizes) closer to OMIM genes (Suppl. Fig. 24d) and, conversely, that more recently identified variants with smaller effects tend to localize further away from OMIM genes (Suppl. Fig. 24e). We also investigated the number of genes prioritised using Summary-data based Mendelian Randomization (hereafter referred to as SMR genes; **Suppl. Methods**) using expression quantitative trait loci (eQTL) as genetic instruments (Suppl. Table 16) as an alternative *gene*-level metric and found it to saturate for N>4M (Suppl. Fig. 24f). Note that saturation of SMR genes is partly affected by the biological relevance and statistical power of eQTL studies.^30^ Therefore, we can expect more genes to be prioritised when integrating GWAS summary statistics from this study with that from larger eQTL studies that may be available in the future and may involve more tissue types. Gene-level metrics were also not substantially affected by adding non-EUR samples, again consistent with broadly similar sets of genes affecting height across ancestries.

At the level of variants and genomic regions, we saw a steady and almost linear increase in the number of GWS SNPs as a function of sample size, as previously reported.^33^ However, given that newly identified variants tend to cluster near ones identified at smaller sample sizes, we also saw a saturation in the number of loci identified for N>2.5M, where the upward trend starts to weaken (Suppl. Fig. 25a). We found a similar pattern for the percentage of the genome covered by GWS loci, with the degree of saturation varying as a function of the window size used to define loci (Suppl. Fig. 25b). The observed saturation in PGS prediction accuracy (both within ancestry, i.e. in EUR; and multi-ancestry) was more noticeable than that of the number and genomic coverage of GWS loci. In fact, increasing sample size from 2.5M to 4M by adding another 1.5M EUR samples increased the number of GWS SNPs from 7,020 to 9,863 (i.e. (9,863-7,020)/7,020 = ~1.4-fold increase) but the absolute increase in prediction accuracy is less than +2.7%. This improvement is mainly observed in EUR but remains lower than +1.3% in EAS and AFR individuals. However, adding another ~1M participants of non-EUR improves the multi-ancestry prediction accuracy by over +3.4% (Suppl. Fig. 25c), highlighting the value of non-EUR populations for this purpose.

Altogether, these analyses show that increasing GWAS sample size not only increases prediction accuracy but also sheds more light on the genomic distribution of causal variants and, at all but the largest sample sizes, the genes proximal to these variants. By contrast, enrichment of higher-level, broadly defined biological categories such as gene sets/pathways and functional annotations can be identified using relatively small sample sizes (N~0.25M for height). Importantly, we confirm that increased genetic diversity in GWAS discovery samples significantly improves the prediction accuracy of PGS in under-represented ancestries.

## DISCUSSION

By performing the largest GWAS to date in 5,380,080 individuals with a primary focus on common genetic variation, we have provided new insights into the genetic architecture of height – including a saturated genomic map of 12,111 genetic associations for height. Consistent with previous studies,^15,16^ we have shown signal density of associations (known and novel) are not randomly distributed across the genome; rather, associated variants are more likely detected around genes previously associated with Mendelian disorders of growth. Furthermore, we observed strong genetic overlap of association across cohorts of various continental ancestries. Effect estimates are moderately to highly correlated (min=0.64, max=0.99), and while there are significant differences in power to detect an association between cohorts with European and non-European ancestries, the majority of genetic associations for height observed in populations with non-European ancestry lie in close proximity and in linkage disequilibrium to associations identified within populations of European ancestry.

By increasing our experimental sample size to >7-times that of previous studies, we have explained up to 40% of the inter-individual variation in height in independent European-ancestry samples using GWS SNPs alone, and >90% of 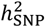 across diverse populations when incorporating all common SNPs within 35 kb of GWS SNPs. This result is important as it highlights that future investigation of common (MAF>1%) genetic variation associated with height in many ancestries will most likely detect signals within the 7,209 GWS loci identified in the present study. An interesting future question is whether rare genetic variants associated with height are also concentrated within the same loci. Of note, previous studies have reported significant enrichment of height heritability near genes as compared to inter-genic regions (e.g. up to >50 kb away from start/stop genomic position of genes).^34^ Our findings are consistent but not reducible to that observation, given that up to ~31% of GWS SNPs identified in this study lie >50 kb away from any gene.

Our study provides a powerful genetic predictor of height based on 12,111 GWS SNPs, for which accuracy reaches ~40% (i.e. 80% of 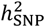) in individuals of European ancestries and up to ~10% in individuals of predominantly African ancestries. Importantly, we show using a new method developed by Wang and colleagues^27^ that LD and MAF differences between European and African ancestries can explain up to ~84% (S.E. 1.5%) of the loss of prediction accuracy between these populations (**Suppl. Methods**), with the remaining loss being presumably explained by heritability differences between populations and/or differences in effect sizes across populations (e.g., due to gene-by-gene or gene-by-environment interactions). This observation is consistent with common causal variants for height being largely shared across ancestries. Therefore, we anticipate that fine-mapping of GWS loci identified in this study, ideally using methods that can accommodate dense sets of signals and large populations with African ancestries, would substantially improve the accuracy of a derived height PGS for non-European ancestry populations. Our study has a large number of participants with African ancestries as compared with previous efforts. However, we emphasise that further increasing the size of GWAS in non-European ancestry populations, including those with diverse African ancestries, is essential to bridge the gap in prediction accuracy, particularly as most studies only partially capture the wide range of ancestral diversity both within Africa and globally. Such increased samples size would help to identify potential ancestry-specific causal variants, to facilitate ancestry-specific fine mapping, and to inform gene by environment/ancestry interactions. Another important finding of our study is to show how individual PGS can be optimally combined with familial information and thereby improve the overall accuracy of height prediction to above 54% in European ancestry populations.

Although large sample sizes are needed to pinpoint the common variants responsible for the heritability of height (and larger samples in multiple ancestries will likely be required to map these at finer scale), the prioritization of relevant genes and gene sets is feasible at smaller sample sizes than that required to account for the common variant heritability. Thus, the sample sizes required for saturation of GWAS are smaller for identifying enriched gene sets, with identification of genes implicated as potentially causal and mapping of genomic regions containing associated variants requiring successively larger sample sizes. Furthermore, unlike prediction accuracy, prioritization of likely causal genes and even mapping of associated regions is consistent across ancestries, reflecting the expected similarity in the biological architecture of human height across populations.

Our study has a number of limitations. First, we focused on SNPs from the HM3 panel, which only partially capture common genetic variation. However, although a significant fraction of height variance can be explained by common SNPs outside the HM3 SNPs panel, we showed that the extra information (also referred to as ‘hidden heritability’) remains concentrated within GWS loci identified from our HM3 SNPs based analyses (Suppl. Fig. 18). This result underlines the widespread allelic heterogeneity at height-associated loci. Another limitation of our study is that we determined conditional associations using an EUR LD reference (N~350,000), which is sub-optimal given that ~24% of our discovery sample is of non-EUR. We emphasise that no analytical tool with an adequately large multi-ancestry reference panel currently is available to properly address how to identify conditionally independent associations in a multi-ancestry study. Fine-mapping of variants remains a particular challenge when attempted across ancestries in loci containing multiple signals (as is often the case for height). A third limitation of our study is our inability to perform well-powered replication analyses of genetic associations specific to populations with non-European ancestries, due to current limited availability of such data. Finally, as with all GWAS, definitive identification of effector genes and the mechanisms by which genes and variants influence phenotype remains a key bottleneck. Therefore, progress towards identifying causal genes from GWAS of height will be mostly driven by the availability of relevant complementary data (e.g., context-specific eQTL in relevant tissues and cell-types) and the power of computational methods that can integrate these data.

In summary, our study has been able to demonstrate empirically that the combined additive effects of tens of thousands of individual variants, detectable with a large enough experimental sample size, can explain substantial variation in a human phenotype. For human height, we show that studies of the order of ~5 million participants of various ancestries provide enough power to map >90% of genetic variance explained by common SNPs down to ~21% of the genome. Height has been used as a model trait for the study of human polygenic traits, including common diseases, because of its high heritability and relative ease of measurement enabling large sample sizes and increased power. Conclusions about the genetic architecture, sample size requirements for additional GWAS discovery, and scope for polygenic prediction that were initially made for height have by-and-large agreed with those for common disease. If the results from this study can also be extrapolated to disease, this would suggest that substantially increased sample sizes could largely resolve the heritability attributed to common variation to a finite set of SNPs (and small genomic regions). These variants and regions would implicate a particular subset of genes, regulatory elements, and pathways that would be most relevant to address questions of function, mechanism and therapeutic intervention.

## Supporting information

Supplementary Information (PDF only) - No Supplementary Table is shared at this point

## ACKNOWLEDGEMENTS

We gratefully acknowledge the participants in each cohorts contributing to this study. Additional acknowledgements are listed in **Supplementary information**. Support for title page creation and format was provided by *AuthorArranger*, a tool developed at the National Cancer Institute. This research was supported by the following funding bodies.

## US National Institutes of Health

75N92021D00001, 75N92021D00002, 75N92021D00003, 75N92021D00004, 75N92021D00005, AA07535, AA10248, AA014041, AA13320, AA13321, AA13326, DA12854, U01 DK062418, HHSN268201800005I, HHSN268201800007I, HHSN268201800003I, HHSN268201800006I, HHSN268201800004I, R01 CA55069, R35 CA53890, R01 CA80205, R01 CA144034, HHSN268201200008I, EY022310, 1X01HG006934-01, R01DK118427, R21DK105913, HHSN268201200036C, HHSN268200800007C, HHSN268200960009C, HHSN268201800001C, N01HC55222, N01HC85079, N01HC85080, N01HC85081, N01HC85082, N01HC85083, N01HC85086, 75N92021D00006, U01HL080295, R01HL085251, R01HL087652, R01HL105756, R01HL103612, R01HL120393, U01HL130114, R01AG023629, UL1TR001881, DK063491, R01 HL095056, 1R01HL139731, 1RO1HL092577 (P.T.E.), K24HL105780 (P.T.E.), HHSC268200782096C, R01 DK087914, R01 DK066358, R01 DK053591, 1K08HG010155 (A.V.K.), 1U01HG011719 (A.V.K.), U01 HG004436, P30 DK072488, HHSN268200782096C, U01 HG 004446, R01 NS45012, U01 NS069208-01, R01-NS114045 (J.W.C.), R01-NS100178 (J.W.C.), R01-NS105150 (J.W.C.), HL043851, HL080467, CA047988, UM1CA182913, U01HG008657, U01HG008685, U01HG008672, U01HG008666, U01HG006379, U01HG008679, U01HG008680, U01HG008684, U01HG008673, U01HG008701, U01HG008676, U01HG008664, U54MD007593, UL1TR001878, R01-DK062370 (M.B.), R01-DK072193 (K.L.M.), intramural project number 1Z01-HG000024 (F.S.C.), N01-HG-65403, DA044283, DA042755, DA037904, AA009367, DA005147, DA036216, 5-P60-AR30701, 5-P60-AR49465, N01-AG-1-2100, HHSN271201200022C, National Institute on Aging Intramural Research Program, R-35-HL135824 (C.J.W.), AA-12502, AA-00145, AA-09203, AA15416, K02AA018755, UM1 CA186107, P01 CA87969, R01 CA49449, U01 CA176726, R01 CA67262, UM1CA167552, CA141298, P01CA055075, CA141298, HL54471, HL54472, HL54473, HL54495, HL54496, HL54509, HL54515, U24 MH068457-06, R01D0042157-01A1, RO1 MH58799-03, MH081802, 1RC2MH089951-01, 1RC2 MH089995, R01 DK092127-04, R01DK110113 (R.J.F.L.), R01DK075787 (R.J.F.L.), R01DK107786 (R.J.F.L.), R01HL142302 (R.J.F.L.), R01HG010297 (R.J.F.L.), R01DK124097 (R.J.F.L.), R01HL151152 (R.J.F.L.), R01-HL046380, KL2-RR024990, R35-HL135818, R01-HL113338, R35HL135818 (Su.R.), HL 046389 (Su.R.), and HL113338 (Su.R.), K01 HL135405 (B.E.C.), R03 HL154284 (B.E.C.), R01HL086718, HG011052 (X.Zhu), N01-HC-25195, HHSN268201500001I, N02-HL-6-4278, R01-DK122503, U01AG023746, U01AG023712, U01AG023749, U01AG023755, U01AG023744, U19AG063893, R01-DK-089256, R01HL117078, R01 HL09135701, R01 HL091357, R01 HL104135, R37-HL045508, R01-HL053353, R01-DK075787, U01-HL054512, R01-HL074166, R01-HL086718, R01-HG003054, U01HG004423, U01HG004446, U01HG004438, DK078150, TW005596, HL085144, RR020649, ES010126, DK056350, R01DK072193, R01 HD30880, R01 AG065357, R01DK104371, R01HL108427, Fogarty grant D43 TW009077, 263 MD 9164, 263 MD 821336, N.1-AG-1-1, N.1-AG-1-2111, HHSN268201800013I, HHSN268201800014I, HHSN268201800015I, HHSN268201800010I, HHSN268201800011I and HHSN268201800012I, KL2TR002490 (L.M.R.), T32HL129982 (L.M.R.), R01AG056477, R01AG034454, R01 HD056465, U01 HL054457, U01 HL054464, U01 HL054481, R01 HL119443, R01 HL087660, U01AG009740, RC2 AG036495, RC4 AG039029, U01AG009740 (W.Zhao.), RC2 AG036495 (W.Zhao.), RC4 AG039029 (W.Zhao.), 75N92020D00001, HHSN268201500003I, N01-HC-95159, 75N92020D00005, N01-HC-95160, 75N92020D00002, N01-HC-95161, 75N92020D00003, N01-HC-95162, 75N92020D00006, N01-HC-95163,

75N92020D00004, N01-HC-95164, 75N92020D00007, N01-HC-95165, N01-HC-95166, N01-HC-95167, N01-HC-95168, N01-HC-95169, UL1-TR-000040, UL1-TR-001079, UL1-TR-001420, N02-HL-64278, UL1TR001881, DK063491, R01-HL088457, R01-HL-60030, R01-HL067974, R01-HL-55005, R01-HL 067974, R01HL111249, R01HL111249-04S1, U01HL54527, U01HL54498, EY014684, EY014684-03S1, EY014684-04S1, DK063491, S10OD017985, S10RR025141, UL1TR002243, UL1TR000445, UL1RR024975, U01HG004798, R01NS032830, RC2GM092618, P50GM115305, U01HG006378, U19HL065962, R01HD074711, 5K08HL135275 (R.W.M.), R01 HL77398 (B.L.), NR013520 (Y.V.S.), DK125187 (Y.V.S.), HHSN268201700001I, HHSN268201700002I, HHSN268201700003I, HHSN268201700004I, HHSN268201700005I, R01HL087641, R01HL086694, U01HG004402, HHSN268200625226C, UL1RR025005, U01HG007416, R01DK101855, 15GRNT25880008, N01-HC65233, N01-HC65234, N01-HC65235, N01-HC65236, N01-HC65237, U01HG007376, HHSN268201100046C, HHSN268201100001C, HHSN268201100002C, HHSN268201100003C, HHSN268201100004C, HHSN271201100004C, N01-AG-6-2101, N01-AG-6-2103, N01-AG-6-2106, R01-AG028050, R01-NR012459, P30AG10161, P30AG72975, R01AG17917, RF1AG15819, R01AG30146, U01AG46152, U01AG61256, AG000513, R01 HD58886, R01 HD100406, N01-HD-1-3228, -3329, -3330, -3331, -3332, -3333, UL1 TR000077, R01 HD056465 (S.F.A.G.), R01 HG010067 (S.F.A.G.), R01CA64277, R01CA15847, UM1CA182910, R01CA148677, R01CA144034, UM1 CA182876, R01DK075787, R01DK075787 (J.N.H.), ZIA CP010152-20, U19 CA 148537-01, U01 CA188392, X01HG007492, HHSN268201200008I, Z01CP010119, R01-CA080122, R01-CA056678, R01-CA082664, R01-CA092579, K05-CA175147, P30-CA015704, CA063464, CA054281, CA098758, CA164973, R01CA128813, K25 HL150334 (R.E.Mu.), DP2 ES030554 (P.-R.L.), U19 CA148065, CA128978, 1U19 CA148537, 1U19 CA148065, 1U19 CA148112, U01 DK062418, U01-DK105535 (M.I.M.), R01HL24799 NIHHLB, U01 DK105556, DK093757 (K.L.M.).

## Wellcome Trust

068545/Z/02, 076113/B/04/Z, Strategic Award 079895, 090532/Z/09/Z, 203141/Z/16/Z, 201543/B/16/Z, 084723/Z/08/Z, 090532, 098381, 217065/Z/19/Z, WT088806, WT092830/Z/10/Z, 202802/Z/16/Z (N.J.T.), 217065/Z/19/Z (N.J.T.), 216767/Z/19/Z, 104036/Z/14/Z, 098051, WT098051, 212946/Z/18/Z, 202922/Z/16/Z, 104085/Z/14/Z, 088158/Z/09/Z, 221854/Z/20/Z, 212904/Z/18/Z, WT095219MA, 068545/Z/02, 076113, 090532 (M.I.M.), 098381 (M.I.M.), 106130 (M.I.M.), 203141 (M.I.M.), 212259 (M.I.M.), 072960/Z/03/Z, 084726/Z/08/Z, 084727/Z/08/Z, 085475/Z/08/Z, 085475/B/08/Z.

## UK Medical Research Council

G0000934, MR/N013166/1 (P.R.H.J.T.), MR/N013166/1 (K.A.K.), U. MC_UU_00007/10, G0601966, G0700931, MRC Integrative Epidemiology Unit MC_UU_00011/1 (N.J.T., R.E.Mi.), MC_UU_00019/1, G9521010D (the BRIGHT Study), MC_UU_12015/1, MC_PC_13046, MC_PC_13049, MC-PC-14135, MC_UU_00017/1, MC_UU_12026/2, MC_U137686851, K013351, R024227, MC_UU_00007/10, MR/M016560/1, G1001799, MC_PC_20026 (L.J.Sm.).

## Cancer Research UK

CRUK Integrative Cancer Epidemiology Programme C18281/A29019 (N.J.T.), C16077/A29186, C500/A16896, C5047/A7357, C1287/A10118, C1287/A16563, C5047/A3354, C5047/A10692, C16913/A6135, C5047/A1232, C490/A10124, C1287/A16563, C1287/A10118, C1287/A10710, C12292/A11174, C1281/A12014, C5047/A8384, C5047/A15007, C5047/A10692, C8197/A16565.

## Australian Research Council

DP0770096 (P.M.Vis.), DP1093502, FL180100072 (P.M.Vis.), DE200100425 (Lo.Y.)

## Australian National Health and Medical Research Council

241944, 389875, 389891, 389892, 389938, 442915, 442981, 496739, 496688, 552485, 613672, 613601, 1011506, APP1172917 (S.E.M.),, 572613, 403981, 1059711, 1027449, 1044840, 1021858, 974159, 211069, 457349, 512423, 302010, 571013, GNT1154518 (D.A.M.), GNT1103329 (A.W.H.), 1186500 (D.F.), 209057, 396414, 1074383, 390130, 1009458, Career Development Fellowship, 1113400 (P.M.Vis., Jian Y.).

## UK National Institute for Health Research Centres

Barts Biomedical Research Centre (Pa.D., S.K.), Comprehensive Biomedical Research Centre Imperial College Healthcare NHS Trust, Health Protection Research Unit on Health Impact of Environmental Hazards, RP-PG-0407-10371, Official Development Assistance award 16/136/68, the University of Bristol NIHR Biomedical Research Centre BRC-1215-2001 (N.J.T.), Academic Clinical Fellowship (S.J.H.), Leicester Cardiovascular Biomedical Research Centre BRC-1215-20010 (C.P.N., P.S.B., N.J.S.), Barts Biomedical Research Centre and Queen Mary University of London, Exeter Clinical Research Facility, Clinical Research Facility and Biomedical Research Centre based at Guy’s and St Thomas’ NHS Foundation Trust and King’s College London (M.Man., C.Par.), Biomedical Research Centre at The Institute of Cancer Research and The Royal Marsden NHS Foundation Trust, Biomedical Research Centre at the University of Cambridge, Oxford Biomedical Research Centre.

## European Union

018996, LSHG-CT-2006-018947, HEALTH-F2-2013-601456, ERA-CVD program grant 01KL1802 (S.W.v.d.L.), 305739, 727565, FP/2007-2013 ERC Grant Agreement number 310644 MACULA, LSHM-CT-2007-037273, SOC 95201408 05F02, SOC 98200769 05F02, LSHM-CT-2006-037593, 279143, iHealth-T2D 643774, 223004, Marie Sklodowska-Curie grant agreement number 786833 (J.R.), 810645, FP7-HEALTH-F4-2007 grant number 201413 and 9602768, QLG1-CT-2001-01252, LSHG-CT-2006-01894 (I.R., A.F.W., V.V.), 733100, HEALTH-F2-2009-223175, LSHG-CT-2006-01947), HEALTH-F4-2007-201413, QLG2-CT-2002-01254, FP7 project number 602633, H2020 project numbers 634935 and 633784, HEALTH-F2-2009-223175, IMI-SUMMIT program, H2020 grants 755320 and 848146 (S.W.v.d.L.), BigData@Heart grant EU IMI 116074 (P.Ki.).

## European Regional Development Fund

2014-2020.4.01.15-0012, 2014-2020.4.01.16-0125 (A.Me.), 539/2010 A31592, 2014-2020.4.01.16-0030.

## Netherlands Heart Foundation

CVON 2011/B019 (S.W.v.d.L.), CVON 2017-20 (S.W.v.d.L.), NHS2010B233, NHS2010B280, CVON 2014–9 (M.R.).

## British Heart Foundation

Centre for Research Excellence (H.W.), RG/14/5/30893 (Pa.D.), FS/14/66/3129 (O.G.), SP/04/002, SP/16/4/32697 (C.P.N.), CH/1996001/9454, 32334 (M.Ki.), RG/17/1/32663, FS/13/43/30324 (P.Ki.), PG/17/30/32961 (P.Ki.), PG/20/22/35093 (P.Ki.).

## US Department of Veterans Affairs

Baltimore Geriatrics Research, Education, and Clinical Center; IK2-CX001780 (S.M.D.), I01-BX004821, MVP 001, IK2-CX001907 (Sr.R.).

## American Heart Association

18SFRN34250007 (S.A.Lu.), 18SFRN34110082 (P.T.E.), 17IBDG33700328 (J.W.C.), 15GPSPG23770000 (J.W.C.), 15POST24470131 (C.N.S.), 17POST33650016 (C.N.S.).

## Leducq Fondation

‘PlaqOmics’ (Ather-Express, S.W.v.d.L), 14CVD01 (P.T.E.).

## Netherlands Organization for Scientific Research NWO

GB-MW 940-38-011, ZonMW Brainpower grant 100-001-004, ZonMw Risk Behavior and Dependence grant 60-60600-97-118, ZonMw Culture and Health grant 261-98-710, GB-MaGW 480-01-006, GB-MaGW 480-07-001, GB-MaGW 452-04-314, GB-MaGW 452-06-004, 175.010.2003.005, 481-08-013, 481-11-001, Vici 016.130.002, 453-16-007/2735, Gravitation 024.001.003, 480-05-003, NWO/SPI 56-464-14192, 480-15-001/674, ZonMW grant number 916.19.151 (H.H.H.A.), ZonMw grant 95103007, 175.010.2005.011, 911-03-012, ZonMw grant 6130.0031, VIDI 016-065-318 (D.P.), Vidi 016.096.309.

## European Research Council

ERC-2017-STG-757364, ERC-CoG-2015-681466, CoG-2015_681742_NASCENT (I.J.), ERC-2011-StG 280559-SEPI, ERC-STG-2015-679242, 742927, ERC-230374.

## Swedish Research Council

2017-02554, 349-2006-237, 2009-1039, Linné grant number 349-2006-237, 2016-06830 (G.H.), 2017-00641, grant for the Swedish Infrastructure for Medical Population-based Life-course Environmental Research.

## Novo Nordisk Foundation

12955 (B.F.), NNF18CC0034900, NNF15OC0015896, NNF18CC0034900, NNF15CC0018486.

## Academy of Finland

77299, 124243, 285547 EGEA, 100499, 205585, 118555, 141054, 264146, 308248, 312073, 265240, 263278, Center of Excellence in Complex Disease Genetics grant number 312062, 329202 (M.Ki.), 322098, 206374, 251360, 276861, 322098, 286284, 134309 (Eye), 126925, 121584, 124282, 129378 (Salve), 117787 (Gendi), and 41071 (Skidi), 263401 (Le. G.), 267882 (Le. G.), 312063 (Le. G.), 336822 (Le. G.), 312072 (T.T.), 336826 (T.T.).

## German Federal Ministry of Education and Research

01ZZ9603, 01ZZ0103, 01ZZ0403, 03IS2061A, 03ZIK012, 01EA1801A (G.E.D.), 01ER0804 (K.-U.E.), BMBF 01ER1206 and BMBF 01ER1507 (I.M.H.), BMBF projects 01EG0401, 01GI0856, 01GI0860, 01GS0820_WB2-C, 01ER1001D, 01GI0205.

## Additional funding

The University of Newcastle Strategic Initiatives Fund; the Gladys M Brawn Senior Research Fellowship scheme; Vincent Fairfax Family Foundation; The Hunter Medical Research Institute; the Nagahama City Office and the Zeroji Club; the Center of Innovation Program, the Global University Project from the Ministry of Education, Culture, Sports, Science and Technology of Japan; the Practical Research Project for Rare/Intractable Diseases (ek0109070, ek0109283, ek0109196, ek0109348), and the Program for an Integrated Database of Clinical and Genomic Information (kk0205008), from the Japan Agency for Medical Research and Development; Takeda Medical Research Foundation; Astellas Pharma, Inc.; Daiichi Sankyo Co., Ltd.; Mitsubishi Tanabe Pharma Corporation; Otsuka Pharmaceutical Co., Ltd.; Taisho Pharmaceutical Co., Ltd.; and Takeda Pharmaceutical Co., Ltd.; Type 1 Diabetes Genetics Consortium; the French Ministry of Research; the Chief Scientist Office of the Scottish Government #CZB/4/276 and #CZB/4/710; Arthritis Research UK; Royal Society URF (J.F.W.); the Atlantic Philanthropies; the UK Economic and Social Research Council awards ES/L008459/1 and ES/L008459/1; the UKCRC Centre of Excellence for Public Health Northern Ireland; the Centre for Ageing Research and Development in Ireland; the Office of the First Minister and Deputy First Minister; the Health and Social Care Research and Development Division of the Public Health Agency; the Wellcome Trust/Wolfson Foundation; and Queen’s University Belfast; the Science Foundation Ireland-Department for the Economy Award 15/IA/3152 (NICOLA); NI HSC R&D division STL/5569/19 (L.J.Sm.); the Italian Ministry of Education, University and Research (MIUR) number 5571/DSPAR/2002 (OGP study); GlaxoSmithKline; the Faculty of Biology and Medicine of Lausanne; the Swiss National Science Foundation grants 33CSCO-122661, 33CS30-139468, 33CS30-148401 and 33CS30_177535/1; the Montreal Heart Institute Biobank; the Canadian Institutes of Health Research PJT #156248; the Canada Research Chair Program, Genome Quebec and Genome Canada, and the Montreal Heart Institute Foundation (G.L.); the Strategic Priority CAS Project grant number XDB38000000, Shanghai Municipal Science and Technology Major Project grant number 2017SHZDZX01, and the National Natural Science Foundation of China grant number 81970684; the National Medical Research Council (grants 0796/2003, 1176/2008, 1149/2008, STaR/0003/2008, 1249/2010, CG/SERI/2010, CIRG/1371/2013, and CIRG/1417/2015) and the Biomedical Research Council (grants 08/1/35/19/550 and 09/1/35/19/616) of Singapore; the Ministry of Health, Singapore; the National University of Singapore and the National University Health System, Singapore; the Agency for Science, Technology and Research, Singapore; Merck Sharp & Dohme Corp., Whitehouse Station, NJ, USA; Kuwait Foundation for Advancements of Sciences (The KODGP); the Oogfonds, MaculaFonds, Landelijke Stichting voor Blinden en Slechtzienden, Stichting Blindenhulp, Stichting A.F. Deutman Oogheelkunde Researchfonds; in Mexico the Fondo Sectorial de Investigación en Salud y Seguridad Social SSA/IMSS/ISSSTECONACYT project 150352; Temas Prioritarios de Salud Instituto Mexicano del Seguro Social 2014-FIS/IMSS/PROT/PRIO/14/34; the Fundación IMSS; Compute Ontario (www.computeontario.ca) and Compute Canada (www.compute.canada.ca); CIHR Operating grants and a CIHR New Investigator Award (E.J.P.); the Westlake Education Foundation (Jian Y.); AstraZeneca; a Miguel Servet contract from the ISCIII Spanish Health Institute number CP17/00142 and co-financed by the European Social Fund (M.S.-L.); the Dutch Ministry of Justice; the European Science Foundation EuroSTRESS project FP-006; Biobanking and Biomolecular Resources Research Infrastructure BBMRI-NL award CP 32; Accare Centre for Child and Adolescent Psychiatry; and the Dutch Brain Foundation; the Federal Ministry of Science, Germany award 01 EA 9401; German Cancer Aid award 70-2488-Ha I; the participating Departments, the Division and the Board of Directors of the Leiden University Medical Centre and the Leiden University, Research Profile Area ‘Vascular and Regenerative Medicine’; Research Project For Excellence IKϒ/SIEMENS; the Wake Forest School of Medicine grant M01 RR07122 and Venture Fund; the Greek General Secretary of Research and Technology award PENED 2003; the MRC-PHE Centre for Environment and Health; the Singapore Ministry of Health’s National Medical Research Council under its Singapore Translational Research Investigator (STaR) Award NMRC/STaR/0028/2017 (J.C.C); the German Research Foundation Project-ID 431984000 - SFB 1453 (M.Wu., Anna K.); the KfH Foundation for Preventive Medicine, and Bayer Pharma AG; the German Research Foundation grant KO 3598/5-1 (Anna K.); the Leipzig Research Center for Civilization Diseases; the Medical Faculty of the University of Leipzig; the Free State of Saxony; the Medical Research Funds from Kangbuk Samsung Hospital (H.-N.K.); the Division of Adult and Community Health, Centers for Disease Control and Prevention; Astra-Zeneca (P.M.R., D.I.C.); Amgen (P.M.R., D.I.C.); a gift from the Smilow family; the Perelman School of Medicine at the University of Pennsylvania; the University of Bristol; a comprehensive list of grants funding is available on the ALSPAC website; the US Centers for Disease Control and Prevention/Association of Schools of Public Health awards S043, S1734, and S3486, and US Centers for Disease Control and Prevention awards U01 DP003206 and U01 DP006266; the Ministry of Cultural Affairs and the Social Ministry of the Federal State of Mecklenburg-West Pomerania; Hjartavernd (the Icelandic Heart Association), and the Althingi (the Icelandic Parliament); Bristol-Myers Squibb; the Netherlands Genomics Initiative’s Netherlands Consortium for Healthy Aging grant 050-060-810; the Netherlands Heart Foundation grant 2001 D 032 (J.W.J.); the Chief Scientist Office of the Scottish Government Health Directorates award CZD/16/6, the Scottish Funding Council award HR03006; the Stiftelsen Kristian Gerhard Jebsen; Faculty of Medicine and Health Sciences, Norwegian University of Science and Technology; Central Norway Regional Health Authority; the Medical Research Council of Canada and the Canadian Institutes of Health Research grant FRN- CCT-83028 (The Quebec Family Study); Pfizer, New York, NY, USA; the Servier Research Group, Paris, France; Leo Laboratories, Copenhagen, Denmark; Estonian Research Council grants PUT 1371, EMBO Installation grant 3573, and The European Regional Development Fund (Kr.L.); the Estonian Research Council grants PUT PRG687, PRG1291 (EstBB, T.E.); the University of Oulu grant number 24000692, Oulu University Hospital grant number 24301140; the Austrian Science Fond grant numbers P20545-P05 and P13180, the Austrian National Bank Anniversary Fund award number P15435, the Austrian Ministry of Science under the aegis of the EU Joint Programme-Neurodegenerative Disease Research (www.jpnd.eu), the Austrian Science Fund P20545-B05, and the Medical University of Graz (ASPS); Wellcome Trust Sanger Institute; the Broad Institute; the Grant of National Center for Global Health and Medicine; the Core Research for Evolutional Science and Technology (CREST) from the Japan Science Technology Agency; the Program for Promotion of Fundamental Studies in Health Sciences, National Institute of Biomedical Innovation Organization; the Grant of National Center for Global Health and Medicine; the German Research Foundation awards HE 3690/7-1 (I.M.H.) and BR 6028/2-1 (Ca.B.); funds from THL and various domestic foundations (The FINRISK surveys); Business Finland through the Personalized Diagnostics and Care program, SalWe Ltd grant number 3986/31/2013; the Finnish Foundation for Cardiovascular Research, the Sigrid Juselius Foundation and University of Helsinki HiLIFE Fellow and Grand Challenge grants (Sa.R.); the Finnish innovation fund Sitra and Finska Läkaresällskapet (E.W.); Netherlands Twin Registry Repository and the Biobanking and Biomolecular Resources Research Infrastructure awards BBMRI–NL, 184.021.007 and 184.033.111; Amsterdam Public Health and Neuroscience Campus Amsterdam; the Avera Institute for Human Genetics, Sioux Falls, South Dakota, USA (The Netherlands Twin Register); the KNAW Academy Professor Award PAH/6635 (D.I.B.); the Netherlands Organization for Scientific Research Geestkracht program grant 10-000-1002; the Center for Medical Systems Biology, Biobanking and Biomolecular Resources Research Infrastructure; VU University’s Institutes for Health and Care Research and Neuroscience Campus Amsterdam; University Medical Center Groningen; Leiden University Medical Center; the Genetic Association Information Network of the Foundation for the National Institutes of Health; the BiG Grid, the Dutch e-Science Grid; The Lundbeck Foundation; the Stanley Medical Research Institute; the Aarhus and Copenhagen universities and university hospitals; the Danish National Biobank resource supported by the Novo Nordisk Foundation; the Robert Dawson Evans Endowment of the Department of Medicine at Boston University School of Medicine and Boston Medical Center; the Economic & Social Research Council award ES/H029745/1; American Diabetes Association Innovative and Clinical Translational Award 1-19-ICTS-068 (J.M.M.); SIGMA; Consejo Naconal de Ciencia y Tecnologia CONACYT grants 2092, M9303, F677M9407, 251M 2005COI (C.G.-V.); the Danish National Research Foundation; the Danish Pharmacists’ Fund; the Egmont Foundation; the March of Dimes Birth Defects Foundation; the Augustinus Foundation; the Health Fund of the Danish Health Insurance Societies; the Oak Foundation fellowship (B.F.); the Nordic Center of Excellence in Health-Related e-Sciences (Xueping.L.); Grants-in-Aid from MEXT numbers 24390169, 16H05250, 15K19242, 16H06277, 19K19434, 20K10514, 21H03206, and a grant from the Funding Program for Next-Generation World-Leading Researchers number LS056; Council of Scientific and Industrial Research, Ministry of Science and Technology, Govt. of India, New Delhi, India; the Lundbeck Foundation grant number R16-A1694; The Danish Ministry of Health grant number 903516; the Danish Council for Strategic Research grant number 0603-00280B; and The Capital Region Research Foundation; the Danish Research Council; the Danish Centre for Health Technology Assessment; Novo Nordisk Inc.; Research Foundation of Copenhagen County; Danish Ministry of Internal Affairs and Health; the Danish Heart Foundation; the Danish Pharmaceutical Association; the Ib Henriksen Foundation; the Becket Foundation; and the Danish Diabetes Association; the Velux Foundation; The Danish Medical Research Council; Danish Agency for Science, Technology and Innovation; The Aase and Ejner Danielsens Foundation; ALK-Abello A/S, Hørsholm, Denmark; and Research Centre for Prevention and Health, the Capital Region of Denmark; the Timber Merchant Vilhelm Bang’s Foundation; the Danish Heart Foundation grant number 07-10-R61-A1754-B838-22392F; the Health Insurance Foundation (Helsefonden) grant number 2012B233 (Health2008); TrygFonden grant number 7-11-0213, the Lundbeck Foundation award R155-2013-14070; the Danish Research Council for Independent Research and by Region of Southern Denmark; the Heinz Nixdorf Foundation; the German Research Council DFG projects EI 969/2-3, ER 155/6-1;6-2, HO 3314/2-1;2-2;2-3;4-3, INST 58219/32-1, JO 170/8-1, KN 885/3-1, PE 2309/2-1, SI 236/8-1;9-1;10-1; the Ministry of Innovation, Science, Research and Technology, North Rhine-Westphalia; Academia Sinica; the Office of Population Studies Foundation in Cebu; the China-Japan Friendship Hospital; Ministry of Health, Chinese National Human Genome Center at Shanghai; Beijing Municipal Center for Disease Prevention and Control; the National Institute for Nutrition and Health, China Center for Disease Control and Prevention; the Canadian Institutes of Health Research grant MOP-82893; WA Health, Government of Western Australia Future Health WA grant G06302; Safe Work Australia; the University of Western Australia (UWA); Curtin University; Women and Infants Research Foundation; Telethon Kids Institute; Edith Cowan University; Murdoch University; The University of Notre Dame Australia; The Raine Medical Research Foundation; the Italian Ministry of Health award ICS110.1/RF97.71; Hong Kong Kadoorie Charitable Foundation; National Natural Science Foundation of China award 91846303; National Key Research and Development Program of China awards 2016YFC 0900500, 0900501, 0900504, 1303904; the KfH Stiftung Präventivmedizin e.V. (C.A.B.); the Else Kröner-Fresenius-Stiftung (2012_A147); the University Hospital Regensburg; the Deutsche Forschungsgemeinschaft (DFG, German Research Foundation) Project-ID 387509280 – SFB 1350 (Subproject C6); the European Union/EFPIA/ JDRF Innovative Medicines Initiative 2 Joint Undertaking grant number 115974; German Research Foundation DFG BO 3815/4-1 (C.A.B.); the Swedish Foundation for Strategic Research; the Swedish Heart-Lung Foundation; Swedish Heart Lung Foundation (A.Po.); VIAgenomics number SP/19/2/344612; the Strategic Cardiovascular Program of Karolinska Institutet and Stockholm County Council; the Foundation for Strategic Research and the Stockholm County Council number 560283; the ALF/LUA research grant in Gothenburg; the Torsten Soderberg Foundation; the ESRC grants ES/S007253/1, ES/T002611/1, and ES/T014083/1 (M.Ku.); Beijing Municipal of Health Reform and Development Project #2019-4 (Beijing Eye Study); the Children’s Hospital of Philadelphia; a Research Development Award from the Cotswold Foundation; the Childreńs Hospital of Philadelphia Endowed Chair in Genomic Research; the Daniel B. Burke Endowed Chair for Diabetes Research; the Italian Ministry of Universities grant IDF SHARID ARS01_01270; the Assessorato Ricerca Regione Campania grant POR CAMPANIA 2000/2006 MISURA 3.16; the Dutch Ministry of Health, Welfare and Sport; the Dutch Ministry of Economic Affairs; the University Medical Center Groningen (UMCG the Netherlands); University of Groningen and the Northern Provinces of the Netherlands; the UMCG Genetics Lifelines Initiative supported by a Spinoza Grant from NWO; University of Michigan discretionary funds; National Institute of Health, Republic of Korea grants 4845–301, 4851–302, 4851–307; Korea National Institute of Health intramural grant 2019-NG-053-02; the Korea Healthcare Technology R&D Project, Ministry of Health and Welfare, Republic of Korea award A102065; the National Research Foundation of Korea Grant 2020R1I1A2075302 (Y.S.C.); the National Research Foundation of Korea Grant NRF-2020R1A2C1012931; the Republic of Croatia Ministry of Science, Education and Sports research grant 108-1080315-0302; the Eye Birth Defects Foundation Inc.; the National Science Council, Taiwan grant NSC 98-2314-B-075A-002-MY3; the Taichung Veterans General Hospital, Taichung, Taiwan grant TCVGH-1003001C; AFNET; EHRA; German Centre for Cardiovascular Research (DZHK); German heart Foundation (DSF); the State of Brandenburg DZD grant 82DZD00302; Sanofi; Abbott; the Victor Chang Cardiac Research Institute; NSW Health; the Center for Translational Molecular Medicine, the University Medical Center Groningen; the Dutch Kidney Foundation grant E0.13; the Netherlands Cardiovascular Research Initiative; the Dell Loy Hansen Heart Foundation (M.J.Cu.); Biosense Webster, ImriCor, and ADAS software (S.N.); the Swedish Heart-Lung Foundation grant 2019-0526; Swedish Foundation for Strategic Research grant IRC15-0067; Skåne University Hospital; governmental funding of clinical research within the Swedish National Health Service; the Knut and Alice Wallenberg Foundation (J.G.S.); the Boettcher Foundation Webb Waring Biomedical Research Award (Sr.R.); the Translational Genomics Research Institute; the Singapore National Medical Research Council grant 1270/2010, and the National Research Foundation, Singapore project 370062002; the Genetic Laboratory of the Department of Internal Medicine, Erasmus MC; the Research Institute for Diseases in the Elderly grant 014-93-015; the Netherlands Genomics Initiative (NGI)/Netherlands Organisation for Scientific Research (NWO) Netherlands Consortium for Healthy Aging project 050-060-810; the Dutch Dairy Association NZO; Netherlands Consortium Healthy Aging, Ministry of Economic Affairs, Agriculture and Innovation project KB-15-004-003; Wageningen University; VU University Medical Center; and Erasmus MC; The Folkhalsan Research Foundation; Nordic Center of Excellence in Disease Genetics; Finnish Diabetes Research Foundation; Foundation for Life and Health in Finland; Finnish Medical Society; Helsinki University Central Hospital Research Foundation; Perklén Foundation; Ollqvist Foundation; Narpes Health Care Foundation; Municipal Heath Care Center and Hospital in Jakobstad; and Health Care Centers in Vasa, Narpes and Korsholm; the Institute of Cancer Research and The Everyman Campaign; The Prostate Cancer Research Foundation; Prostate Research Campaign UK (now PCUK); The Orchid Cancer Appeal; Rosetrees Trust; The National Cancer Research Network UK; The National Cancer Research Institute (NCRI) UK; the Movember Foundation grants D2013-36 and D2013-17; the Morris and Horowitz Families Endowed Professorship; the Swedish Cancer Foundation; Ligue Nationale Contre le Cancer, Institut National du Cancer (INCa); Fondation ARC; Fondation de France; Agence Nationale de sécurité sanitaire de l’alimentation, de l’environnement et du travail (ANSES); Ligue départementale du Val de Marne; the Baden Württemberg Ministry of Science, Research and Arts; The Ronald and Rita McAulay Foundation; Cancer Australia; AICR Netherlands A10-0227; Cancer Council Tasmania; Cancer Councils of Victoria and South Australia; Philanthropic donation to Northshore University Health System; FWO Vlaanderen grants G.0684.12N and G.0830.13N; the Belgian federal government grant KPC_29_023; a Concerted Research Action of the KU Leuven grant GOA/15/017; the Spanish Ministry Council Instituto de Salud Carlos III-FEDER grants PI08/1770, PI09/00773-Cantabria, PI11/01889-FEDER, PI12/00265, PI12/01270, PI12/00715, PI15/00069,and RD09/0076/00036; the Fundación Marqués de Valdecilla grant API 10/09; the Spanish Association Against Cancer (AECC) Scientific Foundation; the Catalan Government DURSI grant 2009SGR1489; the Xarxa de Bancs de Tumors de Catalunya sponsored by Pla Director d’Oncologia de Catalunya (XBTC); the Spanish Ministry of Science and Innovation grant CEX2018-000806-S; the Generalitat de Catalunya; the VicHealth and Cancer Council Victoria; Programa Grupos Emergentes; Cancer Genetics Unit, CHUVI Vigo Hospital; Instituto de Salud Carlos III, Spain; Cancer Australia PdCCRS and Cancer Council Queensland; the California Cancer Research Fund grant 99-00527V-10182; US Public Health Service grants U10CA37429 and 5UM1CA182883; Canadian Cancer Society Research Institute Career Development Award in Cancer Prevention grant 2013-702108; the German Cancer Aid (Deutsche Krebshilfe); The Anthony DeNovi Fund; the Donald C. McGraw Foundation; and the St. Louis Men’s Group Against Cancer; UK Biobank project 12505; the Australian Research Council grant DE200100425 (Lo.Y.); the Australian Research Council grant FL180100072 (P.M.Vis.); Westlake Education Foundation (Jian Y.); a Burroughs Wellcome Fund Career Award, the Next Generation Fund at the Broad Institute of MIT and Harvard, and a Sloan Research Fellowship (P.-R.L.); the Consortium for Systems Biology (NCSB), the Netherlands Genomics Initiative (NGI)/Netherlands Organisation for Scientific Research (NWO); the Government of Canada through Genome Canada and the Canadian Institutes of Health Research grant GPH-129344; the Ministère de l’Économie et de l’Innovation du Québec through Genome Québec grant PSRSIIRI-701; the Quebec Breast Cancer Foundation; the US Department of Defence grant W81XWH-10-1-0341; the Canadian Institutes of Health Research (CIHR) for the CIHR Team in Familial Risks of Breast Cancer; Komen Foundation for the Cure; the Breast Cancer Research Foundation; and the Ovarian Cancer Research Fund; the Economic and Social Research Council grant number ES/M001660/1; Wellcome Investigator and NIHR Senior Investigator (M.I.M.); Council of Scientific and Industrial Research, Government of India grant number BSC0122; the Department of Science and Technology, Government of India through PURSE II CDST/SR/PURSE PHASE II/11 provided to Jawaharlal Nehru University, New Delhi, INDIA; the Deutsche Forschungsgemeinschaft (DFG, German Research Foundation) Projektnummer 209933838 – SFB 1052; B03, C01; SPP 1629 TO 718/2-1; the Competitive Research Funding of the Tampere University Hospital grants 9M048 and 9N035; the Finnish Cultural Foundation; the Finnish Foundation for Cardiovascular Research; the Emil Aaltonen Foundation, Finland; Juho Vainio Foundation; Finnish Cardiac Research Foundation; Finnish Ministry of Education and Culture; Yrjö Jahnsson Foundation; C.G. Sundell Foundation; Special Governmental Grants for Health Sciences Research, Turku University Hospital; Foundation for Pediatric Research; and Turku University Foundation; the Social Insurance Institution of Finland; Competitive State Research Financing of the Expert Responsibility area of Kuopio, Tampere and Turku University Hospitals grant X51001; Paavo Nurmi Foundation; Signe and Ane Gyllenberg Foundation; Diabetes Research Foundation of Finnish Diabetes Association; Tampere University Hospital Supporting Foundation; and Finnish Society of Clinical Chemistry; the Italian Ministry of Health—RC 01/21 (M.P.C.) and D70-RESRICGIROTTO (G.G.); 5 per mille 2015 senses CUP: C92F17003560001 (P.G.); the Helmholtz Zentrum München – German Research Center for Environmental Health, which is funded by the German Federal Ministry of Education and Research (BMBF) and by the State of Bavaria; the Department of Innovation, Research, and University of the Autonomous Province of Bolzano-South Tyrol; the Croatian National Center of Research Excellence in Personalized Healthcare grant number KK.01.1.1.01.0010 (O.Po.) and the Center of Competence in Molecular Diagnostics grant number KK.01.2.2.03.0006 (O.Po.); the Norwegian Research Council Mobility Grant 24014) and Young Research Talent grant 287086; the South-Eastern Health Authorities PhD-grant 2019122; Vestre Viken Hospital Trust PhD-grant; afib.no - the Norwegian Atrial Fibrillation Research Network; “Indremedisinsk Forskningsfond” at Bærum Hospital.

## AUTHOR CONTRIBUTIONS

**Steering committee:**

G.R.A., T.L.A., S.I.B., M.B., D.I.C., Y.S.C., T.E., T.M.F., I.M.H., J.N.H., G.L., C.M.L., A.E.L., R.J.F.L., M.I.M., K.L.M., M.C.Y.N., K.E.N., C.J.O., Y.O., F.Ri., Y.V.S., E.S.T., C.J.W., U.T., P.M.Vis., R.G.W.

**Conveners of GIANT working groups:**

S.I.B., Pa.D., J.N.H., A.E.J., G.L., C.M.L., R.J.F.L., E.M., K.L.M., K.E.N., Y.O., C.N.S., R.G.W., C.J.W., A.R.Wo., Lo.Y.

**Writing Group (drafted, edited, and commented on manuscript):**

E.Ba., J.N.H., G.L., E.M., Y.O., Sr.R., Sa.S., Sa.V., P.M.Vis, A.R.Wo., Lo.Y.

**Coordinated or supervised data collection or analysis specific to manuscript:**

Ad.A., Pa.D., T.E., T.M.F., J.N.H., A.E.J., G.L., A.E.L., P.-R.L., Y.O., K.S., U.T., P.M.Vis., R.G.W., A.R.Wo., Jian Y., Lo.Y.

**Data preparation group (checked and prepared data from contributing cohorts for meta-analyses):**

J.Ar., S.I.B., S.-H.C., T.F., S.E.G., M.Gr., Yi.J., A.E.J., Tu.K., A.E.L., Kr.L., D.E.M., E.M., C.M.-G., M.Ma., A.Moo., Si.R., C.N.S., Sa.V., T.W.W., X.Y., Kr.Y.

**Meta-analysis working group:**

J.N.H., E.M., Sa.V., Lo.Y.

**Primary height analysis working group (post meta-analysis):**

E.Ba., A.D.B., M.Gr., Yu.J., M.Kan., Ku.L., Je.M., E.M., R.E.Mu., Sr.R., Sa.S., Ju.S., Sa.V., A.R.Wo., Lo.Y.

**All other authors were involved in the design, management, coordination, or analysis of contributing studies**

## COMPETING FINANCIAL INTERESTS

Yu.J. is employed by and hold stock or stock options in 23andMe, Inc. T.S.A. is a shareholder in Zealand Pharma A/S and Novo Nordisk A/S. G.C-P is an employee of 23andMe Inc. M.E.K. is employed by SYNLAB Holding Deutschland GmbH. H.L.L. receives support from a consulting contract between Data Tecnica International and the National Institute on Aging (NIA), National Institutes of Health (NIH). As of January 2020, A.Mah. is an employee of Genentech, and a holder of Roche stock. I.N. is an employee and stock owner of Gilead Sciences; this work was conducted before employment by Gilead Sciences. Ji.S. is employed by and hold stock or stock options in 23andMe, Inc. Ca.S. is an employee of Regeneron, Inc. Va.S. is employed by deCODE Genetics/Amgen inc. Since completing the work contributed to this paper, D.J.T. has left the University of Cambridge and is now employed by Genomics plc. G.T. is employed by deCODE Genetics/Amgen inc. H.B. has consultant arrangements with Chiesi Pharmaceuticals and Boehringer Ingelheim. M.J.Ca. is Chief Scientist for Genomics England, a UK Government company. M.J.Cu. has served on Advisory Board or Consulted for Biosense Webster, Janssen Scientific Affairs and Johnson & Johnson. S.M.D. receives research support from RenalytixAI and personal consulting fees from Calico Labs, outside the scope of the current research. P.T.E. receives sponsored research support from Bayer AG and IBM Health, and he has served on advisory boards or consulted for Bayer AG, Quest Diagnostics, MyoKardia and Novartis. P.Ki. has received suppport from several drug and device companies active in atrial fibrillation, and has received honoraria from several such companies in the past, but not in the last three years. P.Ki. is listed as inventor on two patents held by University of Birmingham (Atrial Fibrillation Therapy WO 2015140571, Markers for Atrial Fibrillation WO 2016012783). G.D.K. has given talks, attended conferences and participated in trials sponsored by Amgen, MSD, Lilly, Vianex, Sanofi, and have also accepted travel support to conferences from Amgen, Sanofi, MSD and Elpen. S.A.Lu. receives sponsored research support from Bristol Myers Squibb / Pfizer, Bayer AG, Boehringer Ingelheim, Fitbit, and IBM, and has consulted for Bristol Myers Squibb / Pfizer, Bayer AG, and Blackstone Life Sciences. W.M. reports grants and personal fees from AMGEN, BASF, Sanofi, Siemens Diagnostics, Aegerion Pharmaceuticals, Astrazeneca, Danone Research, Numares, Pfizer, Hoffmann LaRoche: personal fees from MSD, Alexion; grants from Abbott Diagnostics, all outside the submitted work. W.M. is employed with Synlab Holding Deutschland GmbH.

M.A.N. receives support from a consulting contract between Data Tecnica International and the National Institute on Aging (NIA), National Institutes of Health (NIH). S.N. is a scientific advisor to Circle software, ADAS software, CardioSolv, and ImriCor and recieves grant support from Biosense Webster, ADAS software, and ImriCor. Her.S. has received honoraria for consulting from AstraZeneca, MSD/Merck, Daiichi, Servier, Amgen and Takeda Pharma. He has further received honoraria for lectures and/or chairs from AstraZeneca, BayerVital, BRAHMS, Daiichi, Medtronic, Novartis, Sanofi and Servier. P.S. has received research awards from Pfizer Inc. 23andMe Research team are employed by and hold stock or stock options in 23andMe, Inc. The views expressed in this article are those of the author(s) and not necessarily those of the NHS, the NIHR, or the Department of Health. M.I.McC. has served on advisory panels for Pfizer, NovoNordisk and Zoe Global, has received honoraria from Merck, Pfizer, Novo Nordisk and Eli Lilly, and research funding from Abbvie, Astra Zeneca, Boehringer Ingelheim, Eli Lilly, Janssen, Merck, NovoNordisk, Pfizer, Roche, Sanofi Aventis, Servier, and Takeda. As of June 2019, MMcC is an employee of Genentech, and a holder of Roche stock. C.J.O. is a current employee of Novartis Institute of Biomedical Research. U.T. is employed by deCODE Genetics/Amgen inc. K.S. is employed by deCODE Genetics/Amgen inc. Ad.A. is employed by and hold stock or stock options in 23andMe, Inc. C.J.W.’s spouse is employed by Regeneron. A.E.L. is currently employed by and holds stock in Regeneron Pharmaceuticals, Inc. J.N.H. holds equity in Camp4 Therapeutics.

