## Supplementary Information (PDF only) - No Supplementary Table is shared at this point for "A Saturated Map of Common Genetic Variants Associated with Human Height from 5.4 Million Individuals of Diverse Ancestries"

**OVERVIEW OF THE SUPPLEMENTARY INFORMATION**

ACKNOWLEDGEMENTS (p. 3)

SUPPLEMENTARY METHODS (p. 41)
Quality control checks of individuals studies
Genome-wide association study meta-analysis
Hold-out sample from the UK Biobank
Conditional and joint association analyses
$F_{ST}$ calculation and stratified LD score regression
Density of GWS signal and enrichment near OMIM genes
Replication analyses
Variance explained by GWS SNPs and loci
Prediction analyses
Samples used for prediction analyses and estimation of variance explained by SNPs in GWS loci
Contribution of LD and MD to the loss of prediction accuracy in African ancestry participants
Down-sampled GWAS analyses
Gene prioritisation and Summary data-based Mendelian Randomization
Selection of OMIM genes

SUPPLEMENTARY FIGURES (p. 46)
Suppl. Fig. 1. Overview of study design and analytical strategy.
Suppl. Fig. 2. Minor allele frequency distribution of HapMap 3 SNPs across 5 ancestries.
Suppl. Fig. 3. Quantification of confounding due to population stratification.
Suppl. Fig. 4. Colocalization of height-associated signals across ancestries.
Suppl. Fig. 5 - 9. Correlation of marginal SNP effects across ancestry groups.
Suppl. Fig. 10. Replication of marginal associations in the Estonian Biobank.
Suppl. Fig. 11. Correlation of SNPs effects between discovery and replication GWAS.
Suppl. Fig. 12. Schematic representation of the measure of signal density.
Suppl. Fig. 13. Colocalization of high density of GWS SNPs and OMIM genes.
Suppl. Fig. 14. Independent signal density at the *ACAN* gene locus across ancestries.
Suppl. Fig. 15. Haplotypic analysis at the *ACAN* locus.
Suppl. Fig. 16 – 18. Variance explained by SNPs within GWS loci.
Suppl. Fig. 19. Optimal weighting of PGS and parental information in simulated data.
Suppl. Fig. 20. Selection of the number of gene set clusters using the "Elbow method".
Suppl. Fig. 21. Enrichment of height-associated genes identified at various GWAS sample size.
Suppl. Fig. 22. Annotation-level saturation of GWAS discoveries as a function of sample size.
Suppl. Fig. 23. Heritability enrichment within gene sets prioritised with MAGMA and DEPICT
Suppl. Fig. 24. Gene-level saturation of GWAS discoveries as a function of sample size.
Suppl. Fig. 25. Variant-level saturation of GWAS discoveries as a function of sample size.

SUPPLEMENTARY NOTES (p. 72)
Note 1. Investigation of population stratification in a large European ancestry GWAS of height
Note 2. Distinguishing loss of tagging from multiplicity of causal variants at the ACAN locus
Note 3. Optimal weighting of PGS and parental information for height prediction
Note 4. Saturation of GWAS signals within pathways and gene sets

SUPPLEMENTARY TABLES (additional data - spreadsheet)
Suppl. Tab 1. Study descriptive (and samples QC)
Suppl. Tab 2. Genotyping QC
Suppl. Tab 3. Phenotype Summary
Suppl. Tab 4 – 8. List of COJO SNPS from ancestry specific GWAS
Suppl. Tab 9. List of non-EUR ancestry specific COJO SNPS

Suppl. Tab 10. List of COJO-SNPs from trans-ancestry fixed-effect meta-analysis
Suppl. Tab 11. List of OMIM genes
Suppl. Tab 12. Lis of genome-wide significant loci
Suppl. Tab 13. List of Top gene sets per cluster of pathways
Suppl. Tab 14. List of OMIM genes per cluster of pathways
Suppl. Tab 15. Similarity in gene prioritisation within and across ancestries
Suppl. Tab 16. List of genes prioritised by summary data-based Mendelian Randomization

SUPPLEMENTARY REFERENCES (p. 83)

**ACKNOWLEDGEMENTS**

**Cohort acknowledgements and detailed funding information**

**Hunter Community Study (HCS)**
The authors would like to thank the men and women participating in the HCS as well as all the staff, investigators and collaborators who have supported or been involved in the project to date. The University of Newcastle provided $300 000 from its Strategic Initiatives Fund, and $600 000 from the Gladys M Brawn Senior Research Fellowship scheme; Vincent Fairfax Family Foundation, a private philanthropic trust, provided $195 000; The Hunter Medical Research Institute provided media support during the initial recruitment of participants.

**Women's Health Initiative (WHI)**
The WHI program is funded by the National Heart, Lung, and Blood Institute, National Institutes of Health, U.S. Department of Health and Human Services through contracts 75N92021D00001, 75N92021D00002, 75N92021D00003, 75N92021D00004, 75N92021D00005.

**The Nagahama study**
We are extremely grateful to the Nagahama City Office and the nonprofit organization Zeroji Club for their help in performing the Nagahama study. The work was supported by a university grant, the Center of Innovation Program, the Global University Project from the Ministry of Education, Culture, Sports, Science and Technology of Japan; the Practical Research Project for Rare/Intractable Diseases (ek0109070, ek0109283, ek0109196, ek0109348), and the Program for an Integrated Database of Clinical and Genomic Information (kk0205008), from the Japan Agency for Medical Research and Development (AMED), Takeda Medical Research Foundation.

**QIMR**
QIMR gratefully acknowledges the contributions of the participants and their families. QIMR gratefully acknowledges funding from the Australian National Health and Medical Research Council (NHMRC; 241944, 389875, 389891, 389892, 389938, 442915, 442981, 496739, 496688, 552485, 613672, 613601, 1011506 and APP1172917), the US National Institutes of Health (AA07535, AA10248, AA014041, AA13320, AA13321, AA13326 and DA12854) and the Australian Research Council (ARC; DP0770096 and DP1093502). SEM was supported by NHMRC Investigator Grant APP1172917.

**Korea National Diabetes Program (KNDP)**
The KNDP gratefully acknowledges the contributions of the participants and of the study staff. This study was supported by a grant from the Korea Healthcare Technology R&D Project, Ministry of Health and Welfare, Republic of Korea (A102065).

**Japan PGx Data Science Consortium (JPDSC)**
The authors thank the Japan PGx Data Science Consortium (JPDSC) for kindly providing genotype and phenotype data. The JPDSC was comprised of six pharmaceutical companies in Japan, namely Astellas Pharma, Inc.; Daiichi Sankyo Co., Ltd.; Mitsubishi Tanabe Pharma Corporation; Otsuka Pharmaceutical Co., Ltd.; Taisho Pharmaceutical Co., Ltd.; and Takeda Pharmaceutical Co., Ltd.

**British 1958 birth cohort (B58C)**
We acknowledge use of phenotype and genotype data from the British 1958 Birth Cohort DNA collection, funded by the Medical Research Council grant G0000934 and the Wellcome Trust grant 068545/Z/02. Genotyping for the B58C-WTCCC subset was funded by the Wellcome Trust grant 076113/B/04/Z. The B58C-T1DGC genotyping utilized resources provided by the Type 1 Diabetes Genetics Consortium, a collaborative clinical study sponsored by the National Institute of Diabetes and Digestive and Kidney Diseases (NIDDK), National Institute of Allergy and Infectious Diseases (NIAID), National Human Genome Research Institute (NHGRI), National Institute of Child Health and Human

Development (NICHD), and Juvenile Diabetes Research Foundation International (JDRF) and supported by U01 DK062418. B58C-T1DGC GWAS data were deposited by the Diabetes and Inflammation Laboratory, Cambridge Institute for Medical Research (CIMR), University of Cambridge, which is funded by Juvenile Diabetes Research Foundation International, the Wellcome Trust and the National Institute for Health Research Cambridge Biomedical Research Centre; the CIMR is in receipt of a Wellcome Trust Strategic Award (079895). The B58C-GABRIEL genotyping was supported by a contract from the European Commission Framework Programme 6 (018996) and grants from the French Ministry of Research.

**Aneurysm-Express (AAA)**
Claudia Tersteeg, Krista den Ouden, Mirjam B. Smeets, and Loes B. Collé are graciously acknowledged for their work on the DNA extraction. Astrid E.M.W. Willems, Evelyn Velema, Kristy M. J. Vons, Sara Bregman, Timo R. ten Brinke, Sara van Laar, Louise M. Catanzariti, Joyce E.P. Vrijenhoek, Sander M. van de Weg, Arjan H. Schoneveld, Arnold Koekman, Arjan Boltjes, Petra H. Homoed-van der Kraak, and Aryan Vink are graciously acknowledged for their past and continuing work on the Aneurysm-Express Biobank Study. We would also like to thank all the (former) employees involved in the Aneurysm-Express Biobank Study of the Departments of Surgery of the St. Antonius Hospital Nieuwegein and University Medical Center Utrecht for their continuing work. Lastly, we would like to thank all participants of the Aneurysm-Express Biobank Study; without you these kinds of studies would not be possible.

**Ather-Express (AE)**
Claudia Tersteeg, Krista den Ouden, Mirjam B. Smeets, and Loes B. Collé are graciously acknowledged for their work on the DNA extraction. Astrid E.M.W. Willems, Evelyn Velema, Kristy M. J. Vons, Sara Bregman, Timo R. ten Brinke, Sara van Laar, Louise M. Catanzariti, Joyce E.P. Vrijenhoek, Sander M. van de Weg, Arjan H. Schoneveld, Arnold Koekman, Arjan Boltjes, Petra H. Homoed-van der Kraak, and Aryan Vink are graciously acknowledged for their past and continuing work on the Athero-Express Biobank Study. We would also like to thank all the (former) employees involved in the Athero-Express Biobank Study of the Departments of Surgery of the St. Antonius Hospital Nieuwegein and University Medical Center Utrecht for their continuing work. Lastly, we would like to thank all participants of the Athero-Express Biobank Study; without you these kinds of studies would not be possible.

**Orkney Complex Disease Study (ORCADES)**
DNA extractions were performed at the Edinburgh Clinical Research Facility, University of Edinburgh. We would like to acknowledge the invaluable contributions of the research nurses in Orkney, the administrative team in Edinburgh and the people of Orkney. PRHJT acknowledges funding from the Medical Research Council Doctoral Training Programme in Precision Medicine (MR/N013166/1). The Orkney Complex Disease Study (ORCADES) was supported by the Chief Scientist Office of the Scottish Government (CZB/4/276, CZB/4/710), a Royal Society URF to J.F.W., the MRC Human Genetics Unit quinquennial programme "QTL in Health and Disease", Arthritis Research UK and the European Union framework program 6 EUROSPAN project (contract no. LSHG-CT-2006-018947).

**Viking Health Study - Shetland (VIKING)**
DNA extractions and genotyping were performed at the Edinburgh Clinical Research Facility, University of Edinburgh. We would like to acknowledge the invaluable contributions of the research nurses in Shetland, the administrative team in Edinburgh and the people of Shetland. KAK acknowledges funding from the Medical Research Council Doctoral Training Programme in Precision Medicine (MR/N013166/1). VIKING was supported by the MRC Human Genetics Unit programme grant, "Quantitative traits in health and disease" (U. MC_UU_00007/10).

**Northern Ireland Cohort for Longitudinal Study of Ageing (NICOLA)**
We are grateful to all the participants of the NICOLA Study, and the whole NICOLA team, which includes nursing staff, research scientists, clerical staff, computer and laboratory technicians, managers and

receptionists. The authors alone are responsible for the interpretation of the data and any views or opinions presented are solely those of the authors and do not necessarily represent those of the NICOLA Study team. Support for the study came from The Atlantic Philanthropies, the Economic and Social Research Council, the UKCRC Centre of Excellence for Public Health Northern Ireland, the Centre for Ageing Research and Development in Ireland, the Office of the First Minister and Deputy First Minister, the Health and Social Care Research and Development Division of the Public Health Agency, the Wellcome Trust/Wolfson Foundation and Queen's University Belfast provide core financial support for NICOLA. The analysis of molecular biomarkers for NICOLA's Wave 1 was funded by the Economic and Social Research Council, award reference ES/L008459/1. Generic analysis of data was supported by ESRC (ES/L008459/1) and the Science Foundation Ireland-Department for the Economy (SFI-DfE) Investigator Program Partnership Award (15/IA/3152). LJS is supported by an award from NI HSC R&D division STL/5569/19; UKRI (Medical Research Council) MC_PC_20026.

**Ogliastra Genetic Park (OGP)**
The OGP expresses its gratitude to all the study participants for their contributions and to the municipal administrations for their economic and logistic support. The OGP study was supported by grant from the Italian Ministry of Education, University and Research (MIUR) n°: 5571/DSPAR/2002.

**CARDIA**
The Coronary Artery Risk Development in Young Adults Study (CARDIA) is conducted and supported by the National Heart, Lung, and Blood Institute (NHLBI) in collaboration with the University of Alabama at Birmingham (HHSN268201800005I & HHSN268201800007I), Northwestern University (HHSN268201800003I), University of Minnesota (HHSN268201800006I), and Kaiser Foundation Research Institute (HHSN268201800004I).

**LURIC**
We thank the LURIC study team who were either temporarily or permanently involved in patient recruitment as well as sample and data handling, in addition to the laboratory staff at the Ludwigshafen General Hospital and the Universities of Freiburg, Ulm, and Heidelberg, Germany. LURIC was supported by the 7th Framework Program RiskyCAD (grant agreement number 305739) of the European Union and the H2020 Program TO_AITION (grant agreement number 848146) of the European Union. The work of G.E.D. is supported by the European Union's Horizon 2020 research and innovation programme under the ERA-Net Cofund action N° 727565 (OCTOPUS project) and the German Ministry of Education and Research (grant number 01EA1801A).

**Cohorte Lausannoise (CoLaus)**
The authors would like to thank all the people who participated in the recruitment of the participants, data collection and validation, particularly Nicole Bonvin, Yolande Barreau, Mathieu Firmann, François Bastardot, Julien Vaucher, Panagiotis Antiochos, Cédric Gubelmann, Marylène Bay and Benoit Delabays. The CoLaus study was and is supported by research grants from GlaxoSmithKline, the Faculty of Biology and Medicine of Lausanne, and the Swiss National Science Foundation (grants 33CSCO-122661, 33CS30-139468, 33CS30-148401 and 33CS30_177535/1).

**Montreal Heart Institute Biobank (MHIBB)**
We thank all participants and staff of the André and France Desmarais MHI Biobank. The MHIBB was funded by the Montreal Heart Institute Biobank. GL funded by the Canadian Institutes of Health Research (PJT #156248), the Canada Research Chair Program, Genome Quebec and Genome Canada, and the Montreal Heart Institute Foundation.

**Nutrition and Health of Aging Population in China (NHAPC)**
We are grateful to all participants of the NHAPC study, and also thank our colleagues at the laboratory and local CDC staffs of Beijing and Shanghai for their assistance with data collection. The NHPAC study are supported by the Strategic Priority CAS Project (grant number XDB38000000), Shanghai Municipal

Science and Technology Major Project (grant number 2017SHZDZX01), and the National Natural Science Founda-tion of China (grant number 81970684).

**Diabetic Cohort (DC)**
The Diabetic Cohort (DC) was supported by the individual research grant from the National Medical Research Council (NMRC) and the Biomedical Research Council (BMRC) of Singapore.

**Living Biobank**
The Living Biobank was supported by grants from the Ministry of Health, Singapore, the National University of Singapore and the National University Health System, Singapore. In addition, genotyping for Living Biobank was funded by the Agency for Science, Technology and Research, Singapore, and Merck Sharp & Dohme Corp., Whitehouse Station, NJ, USA.

**SEED (SCES + SiMES + SINDI)**
The Singapore Chinese Eye Study (SCES), the Singapore Malay Eye Study (SiMES), and the Singapore Indian Eye Study (SINDI) are supported by the National Medical Research Council (NMRC), Singapore (grants 0796/2003, 1176/2008, 1149/2008, STaR/0003/2008, 1249/2010, CG/SERI/2010, CIRG/1371/2013, and CIRG/1417/2015), and Biomedical Research Council (BMRC), Singapore (08/1/35/19/550 and 09/1/35/19/616).

**Singapore Chinese Health Study (SCHS-T2D)**
The Singapore Chinese Health Study was supported by U.S. NIH/NCI grants: RO1 CA55069, R35 CA53890, R01 CA80205, and R01 CA144034.

**Singapore Prospective Study Program (SP2)**
The Singapore Prospective Study Program (SP2) were supported by the individual research grant and clinician scientist award schemes from the National Medical Research Council (NMRC) and the Biomedical Research Council (BMRC) of Singapore.

**Kuwait Obesity and Diabetes Genetics Programme (KODGP )**
The KODGP gratefully acknowledges the contributions of the participants and of the study staff. The KODGP was supported by institutional funding by Kuwait Foundation for Advancements of Sciences.

**EUGENDA**
The EUGENDA cohort was funded by grants from the Oogfonds, MaculaFonds, Landelijke Stichting voor Blinden en Slechtzienden, Stichting Blindenhulp, Stichting A.F. Deutman Oogheelkunde Researchfonds, the Dutch Research Council (Vidi Innovational Research Award 016.096.309), and the European Research Council under the European Union's Seventh Framework Programme (FP/2007-2013) (ERC Grant Agreement no. 310644 MACULA). The EUGENDA samples were genotyped as part of the IAMDGC exome chip project supported by CIDR (contract number HHSN268201200008I) and funded by EY022310 (to Jonathan L. Haines, Case Western Reserve University, Cleveland) and 1 × 01HG006934-01 (to Gonçalo R. Abecasis, University of Michigan, Department of Biostatistics).

**Mexico City 1 and Mexico City 2 (MC1 & MC2)**
MC1 & MC2 gratefully acknowledge the contributions of the participants. We thank Miguel Alexander Vazquez Moreno, Daniel Locia and Araceli Méndez Padrón for technical support in Mexico. The Mexico City 1 and Mexico City 2 studies were supported in Mexico by the Fondo Sectorial de Investigación en Salud y Seguridad Social (SSA/IMSS/ISSSTECONACYT, project 150352), Temas Prioritarios de Salud Instituto Mexicano del Seguro Social (2014-FIS/IMSS/PROT/PRIO/14/34), and the Fundación IMSS. In Canada, this research was enabled in part by two CIHR Operating grants to EJP, a CIHR New Investigator Award to EJP and by support provided by Compute Ontario (www.computeontario.ca), and Compute Canada (www.compute.canada.ca).

**Wenzhou Medical University Biobank - Tibetans (WMUB-T)**
The WMUB-T gratefully acknowledges the contributions of the participants and of the study staff. Jian Y. is funded by the Westlake Education Foundation

**Precocious Coronary Artery Disease (PROCARDIS)**
PROCARDIS was supported by the European Community Sixth Framework Program (LSHM-CT-2007-037273), AstraZeneca, the Swedish Research Council, the Knut and Alice Wallenberg Foundation, the Swedish Heart-Lung Foundation, the Torsten and Ragnar Soderberg Foundation, the Strategic Cardiovascular Program of Karolinska Institutet and Stockholm County Council, the Foundation for Strategic Research and the Stockholm County Council (560283). Wellcome Trust core award (090532/Z/09/Z, 203141/Z/16/Z, 201543/B/16/Z); HEALTH-F2-2013-601456 (CVGenes@Target), the TriPartite Immunometabolism Consortium [TrIC]-Novo Nordisk Foundation's Grant number NNF15CC0018486, VIAgenomics (SP/19/2/344612) and support from the NIHR Oxford Biomedical Research Centre. The views expressed are those of the author(s) and not necessarily those of the NHS, the NIHR or the Department of Health. HW is supported by the British Heart Foundation Centre for Research Excellence. Maria Sabater-Lleal is supported by a Miguel Servet contract from the ISCIII Spanish Health Institute (CP17/00142) and co-financed by the European Social Fund.

**Asian Indian Diabetic Heart Study/Sikh Diabetes Study (AIDHS/SDS)**
The AIDHS/SDS gratefully acknowledges the contributions of the participants and of the study staff. AIDHS/SDS was funded by NIH Grants (R01DK118427, R21DK105913)

**Cardiovascular Health Study (CHS)**
This CHS research was supported by NHLBI contracts HHSN268201200036C, HHSN268200800007C, HHSN268200960009C, HHSN268201800001C, N01HC55222, N01HC85079, N01HC85080, N01HC85081, N01HC85082, N01HC85083, N01HC85086, 75N92021D00006; and NHLBI grants U01HL080295, R01HL085251, R01HL087652, R01HL105756, R01HL103612, R01HL120393, and U01HL130114 with additional contribution from the National Institute of Neurological Disorders and Stroke (NINDS). Additional support was provided through R01AG023629 from the National Institute on Aging (NIA). A full list of principal CHS investigators and institutions can be found at CHS-NHLBI.org. The provision of genotyping data was supported in part by the National Center for Advancing Translational Sciences, CTSI grant UL1TR001881, and the National Institute of Diabetes and Digestive and Kidney Disease Diabetes Research Center (DRC) grant DK063491 to the Southern California Diabetes Endocrinology Research Center. The content is solely the responsibility of the authors and does not necessarily represent the official views of the National Institutes of Health.

**Hoorn DCS Cohort**
The authors thank participants and staff of the Diabetes Care System West-Friesland.

**Tracking Adolescents' Individual Lives Survey - Population Cohort (TRAILS Pop) and Clinical Cohort (TRAILS CC)**
We are grateful to all adolescents who participated in this research and to everyone who worked on this project and made it possible. TRAILS has been financially supported by grants from the Netherlands Organization for Scientific Research NWO (Medical Research Council program grant GB-MW 940-38-011; ZonMW Brainpower grant 100-001-004; ZonMw Risk Behavior and Dependence grant 60-60600-97-118; ZonMw Culture and Health grant 261-98-710; Social Sciences Council medium-sized investment grants GB-MaGW 480-01-006 and GB-MaGW 480-07-001; Social Sciences Council project grants GB-MaGW 452-04-314 and GB-MaGW 452-06-004; NWO large-sized investment grant 175.010.2003.005; NWO Longitudinal Survey and Panel Funding 481-08-013 and 481-11-001; NWO Vici 016.130.002 and 453-16-007/2735; NWO Gravitation 024.001.003), the Dutch Ministry of Justice (WODC), the European Science Foundation (EuroSTRESS project FP-006), the European Research Council (ERC-2017-STG-757364 en ERC-CoG-2015-681466), Biobanking and Biomolecular Resources Research Infrastructure BBMRI-NL (CP 32), the participating universities, and Accare Centre for Child

and Adolescent Psychiatry. Statistical analyses were carried out on the Genetic Cluster Computer (http://www.geneticcluster.org), which is financially supported by the Netherlands Scientific Organization (NWO 480-05-003) along with a supplement from the Dutch Brain Foundation.

**Gene-Lifestyle Interactions and Complex Traits Involved in Elevated Disease Risk V2 (GLACIERV2)**
We are grateful to the study participants, health professionals, investigators, data managers and support staff who have contributed to GLACIERV2 and to the Northern Sweden Health and Disease Study. Individual investigator effort was funded by Swedish Research Council, Novo Nordisk Foundation, Swedish Heart Lung Foundation, and European Research Council (CoG-2015_681742_NASCENT).

**European Prospective Investigation into Cancer and Nutrition (EPIC) - Potsdam cohort**
We thank the Human Study Centre (HSC) of the German Institute of Human Nutrition Potsdam-Rehbrücke, namely, the trustee and the data hub for the processing, and the participants for the provision of the data, the biobank for the processing of the biological samples, and the head of the HSC, Manuela Bergmann, for the contribution to the study design and leading the underlying processes of data generation. The recruitment phase of the EPIC-Potsdam Study was supported by the Federal Ministry of Science, Germany (01 EA 9401) and the European Union (SOC 95201408 05F02). The follow-up of the EPIC-Potsdam Study was supported by German Cancer Aid (70-2488-Ha I) and the European Community (SOC 98200769 05F02). This work was supported by a grant from the German Ministry of Education and Research (BMBF) and the State of Brandenburg (DZD grant 82DZD00302).

**Netherlands Epidemiology of Obesity Study (NEO)**
We greatly appreciate all participants of the Netherlands Epidemiology of Obesity study, and all participating general practitioners for inviting eligible individuals. We furthermore thank P.R. van Beelen and all research nurses for collecting the data, P.J. Noordijk and her team for sample handling and storage and I. de Jonge, MSc for all data management of the NEO study. The NEO study is supported by the participating Departments, the Division and the Board of Directors of the Leiden University Medical Centre, and by the Leiden University, Research Profile Area 'Vascular and Regenerative Medicine.

**Tromso-6 Migraine (TromsoMig)**
We thank all participants that attended the Tromsø Study.

**Mexican hypertriglyceridemia (MexTG)**
The MexTG cohort gratefully acknowledges the contributions of the participants of the study staff. The MexTG cohort was funded by NIH grant R01 HL095056.

**MGH Cardiology and Metabolic Patient Cohort (CAMP)**
The MGH Cardiology and Metabolic Patient Cohort is comprised of 3850 subjects recruited from the ambulatory MGH Cardiology Practice between 2009 and 2012. Dr. Lubitz is supported by NIH grant 1R01HL139731 and American Heart Association 18SFRN34250007. Dr. Ellinor is supported by the Fondation Leducq (14CVD01), the NIH (1RO1HL092577, K24HL105780) and the American Heart Association (18SFRN34110082).

**Greek Recurrent Myocardial Infarction Cohort (GRMIC)**
The GRMIC thanks all study participants who contributed to this study and all individuals who have contributed to patients' recruitments and sample collection. This work was supported by the British Heart Foundation (BHF) grant RG/14/5/30893 (P.D.) and forms part of the research themes contributing to the translational research portfolios of the Barts Biomedical Research Centre funded by the UK National Institute for Health Research (NIHR). O.G. received funding from the British Heart Foundation (BHF) (FS/14/66/3129).

**NAFLD study (NAFLD)**
The NAFLD study is partially supported by "Research Project For Excellence IKY/SIEMENS". O.G. received funding from the British Heart Foundation (BHF) (FS/14/66/3129).

**Wake Forest School of Medicine Study (WFSM)**
WFSM gratefully acknowledges the contributions of the participants and of the study staff. Genotyping services were provided by the Center for Inherited Disease Research (CIDR). CIDR is fully funded through a federal contract from the National Institutes of Health to The Johns Hopkins University, contract number HHSC268200782096C. This work was supported by National Institutes of Health grants R01 DK087914, R01 DK066358, R01 DK053591, U01 DK105556 and by the Wake Forest School of Medicine grant M01 RR07122 and Venture Fund.

**The Hellenic study of Interactions between Single nucleotide polymorphisms and Eating in Atherosclerosis Susceptibility (THISEAS)**
THISEAS is partially supported by a research grant (PENED 2003) from the Greek General Secretary of Research and Technology. Work supported by the Barts Biomedical Research Centre funded by the UK National Institute for Health Research (NIHR).

**CARDIOGENICS**
The main sponsor of Cardiogenics was the EU (LSHM-CT-2006-037593). Work supported by the Barts Biomedical Research Centre funded by the UK National Institute for Health Research (NIHR).

**London Life Sciences Prospective Population Study (LOLIPOP)**
The LOLIPOP study is supported by the National Institute for Health Research (NIHR) Comprehensive Biomedical Research Centre Imperial College Healthcare NHS Trust. We acknowledge support of the MRC-PHE Centre for Environment and Health, and the NIHR Health Protection Research Unit on Health Impact of Environmental Hazards. The work was carried out in part at the NIHR/Wellcome Trust Imperial Clinical Research Facility. The views expressed are those of the author(s) and not necessarily those of the Imperial College Healthcare NHS Trust, the NHS, the NIHR or the Department of Health. We thank the participants and research staff who made the study possible. The LOLIPOP was funded by the British Heart Foundation (SP/04/002), the Medical Research Council (G0601966, G0700931), the Wellcome Trust (084723/Z/08/Z, 090532 & 098381) the NIHR (RP-PG-0407-10371), the NIHR Official Development Assistance (ODA, award 16/136/68), the European Union FP7 (EpiMigrant, 279143) and H2020 programs (iHealth-T2D, 643774). JC is supported by the Singapore Ministry of Health's National Medical Research Council under its Singapore Translational Research Investigator (STaR) Award (NMRC/STaR/0028/2017).

**German Chronic Kidney Disease Study (GCKD)**
We are grateful for the willingness of the patients to participate in the GCKD study. The enormous effort of the study personnel of the various regional centers is highly appreciated. We thank the large number of nephrologists who provide routine care for the patients and collaborate with the GCKD study. The GCKD study was/is funded by grants from the Federal Ministry of Education and Research (BMBF, grant number 01ER0804, K.U.E.) and the KfH Foundation for Preventive Medicine. Genotyping was supported by Bayer Pharma AG. The work of A.K. was supported by the German Research Foundation (DFG) - Project-ID 431984000 - SFB 1453, and DFG grant KO 3598/5-1. The work of M.W. was supported by the German Research Foundation (DFG) - Project-ID 431984000 - SFB 1453.

**LIFE-Adult**
We thank all participants of the LIFE-Adult study for spending their time and blood samples. LIFE-Adult genotyping was performed at the Cologne Center for Genomics (CCG, University of Cologne, Peter Nuernberg and Mohammad R. Toliat). For genotype imputation, compute infrastructure provided by ScaDS (Dresden/Leipzig Competence Center for Scalable Data Services and Solutions) at the Leipzig

University Computing Centre was used. LIFE-Adult is funded by the Leipzig Research Center for Civilization Diseases (LIFE). LIFE is an organizational unit affiliated to the Medical Faculty of the University of Leipzig. LIFE is funded by means of the European Union, by the European Regional Development Fund (ERDF) and by funds of the Free State of Saxony within the framework of the excellence initiative.

**PROMIS**
AVK was funded by NHGRI 1K08HG010155. GH was funded by the Swedish Research Council grant 2016-06830.

**Ewha Womans University Hospital PCOS Study (pcosN)**
The pcosN gratefully acknowledges the contributions of the participants and of the study staff. YSC acknowledges support from the National Research Foundation of Korea (NRF) Grant (2020R1I1A2075302).

**Catholic University Incheon ST Mary's Hospital Eye Study (EYE)**
The EYE gratefully acknowledges the contributions of the participants and of the study staff. YSC acknowledges support from the National Research Foundation of Korea (NRF) Grant (2020R1I1A2075302).

**Yonsei Avellino Corneal Dystrophy Study (ACD)**
The ACD gratefully acknowledges the contributions of the participants and of the study staff. We are thankful for the computing resources provided by the Global Science experimental Data hub Center (GSDC) Project and the Korea Research Environment Open NETwork (KREONET) of the Korea Institute of Science and Technology Information (KISTI).

**Kangbuk Samsung Cohort Study (KSCS)**
The KSCS gratefully acknowledges the contributions of the participants and of the study staff. We are thankful for the computing resources provided by the Global Science experimental Data hub Center (GSDC) Project and the Korea Research Environment Open NETwork (KREONET) of the Korea Institute of Science and Technology Information (KISTI). HNK acknowledges support from the National Research Foundation of Korea (NRF) Grant (NRF-2020R1A2C1012931) and the Medical Research Funds from Kangbuk Samsung Hospital.

**Genetics of Early-Onset Stroke (GEOS)**
The research team gratefully acknowledges the contributions of the participants in the GEOS study. The GEOS Study was supported by the National Institutes of Health Genes, Environment and Health Initiative (GEI) Grant U01 HG004436, as part of the GENEVA consortium under GEI, with additional support provided by the Mid-Atlantic Nutrition and Obesity Research Center (P30 DK072488); and the Office of Research and Development, Medical Research Service, and the Baltimore Geriatrics Research, Education, and Clinical Center of the Department of Veterans Affairs. Genotyping services were provided by the Johns Hopkins University Center for Inherited Disease Research (CIDR), which is fully funded through a federal contract from the National Institutes of Health to the Johns Hopkins University (contract number HHSN268200782096C). Assistance with data cleaning was provided by the GENEVA Coordinating Center (U01 HG 004446; PI Bruce S Weir). Study recruitment and assembly of datasets were supported by a Cooperative Agreement with the Division of Adult and Community Health, Centers for Disease Control and Prevention, and by grants from the National Institute of Neurological Disorders and Stroke (NINDS) and the NIH Office of Research on Women's Health (R01 NS45012, U01 NS069208-01). Dr. Cole was partially supported by an American Heart Association (AHA)-Bayer Discovery Grant (Grant 17IBDG33700328), the AHA Cardiovascular Genome-Phenome Study (Grant-15GPSPG23770000), NIH (Grants: R01-NS114045; R01-NS100178; R01-NS105150), and the US Department of Veterans Affairs.

**Justification for the Use of Statins in Primary Prevention: an Intervention Trial Evaluating Rosuvastatin (JUPITER)**
The JUPITER trial (PI: Ridker) and its substudy for genetics (Pis: Chasman and Ridker) were funded by Astra-Zeneca.

**Women's Genome Health Study (WGHS)**
The WGHS is supported by the National Heart, Lung, and Blood Institute (HL043851 and HL080467) and the National Cancer Institute (CA047988 and UM1CA182913) to Julie Buring and I-Min Lee, with funding for genotyping provided by Amgen. (Ridker, Chasman, PIs).

**Electronic Medical Records and Genomics Network (eMERGE)**
The eMERGE network gratefully acknowledges the contributions of the participants and of the study staff at each eMERGE site. This phase of the eMERGE Network was initiated and funded by the NHGRI through the following grants: U01HG008657 (Group Health Cooperative/University of Washington); U01HG008685 (Brigham and Women's Hospital); U01HG008672 (Vanderbilt University Medical Center); U01HG008666 (Cincinnati Children's Hospital Medical Center); U01HG006379 (Mayo Clinic); U01HG008679 (Geisinger Clinic); U01HG008680 (Columbia University Health Sciences); U01HG008684 (Children's Hospital of Philadelphia); U01HG008673 (Northwestern University); U01HG008701 (Vanderbilt University Medical Center serving as the Coordinating Center); U01HG008676 (Partners Healthcare/Broad Institute); U01HG008664 (Baylor College of Medicine); and U54MD007593 (Meharry Medical College). eMERGE Network Banner author list: Murray Brilliant, Wendy Chung, Paul Crane, Damien Croteau-Chonka, Josh Denny, Todd Edwards, Geoff Hayes, Scott Hebbring, George Hripsak, Krzysztok Kiryluk, Terrie Kitchner, Iftikhar Kullo, Bahram Namjou, Peggy Peissig, Ning Shang, Digna Velez Edwards, Chunhua Weng.

**Penn Medicine Biobank (PMBB)**
The PMBB gratefully acknowledges the contributions of the participants and of the study staff. The Penn Medicine BioBank is funded by a gift from the Smilow family, the National Center for Advancing Translational Sciences of the National Institutes of Health under CTSA Award Number UL1TR001878, and the Perelman School of Medicine at the University of Pennsylvania. SMD is supported by IK2-CX001780. This publication does not represent the views of the Department of Veterans Affairs or the United States Government.

**Avon Longitudinal Study of Parents and Children (ALSPAC)**
The UK Medical Research Council and Wellcome (Grant ref: 217065/Z/19/Z) and the University of Bristol provide core support for ALSPAC. A comprehensive list of grants fundingis available on the ALSPAC website.This research was specifically funded by Wellcome (Grant ref: WT088806 and WT092830/Z/10/Z). NJT is a Wellcome Trust Investigator (202802/Z/16/Z), is the PI of the Avon Longitudinal Study of Parents and Children (MRC & WT 217065/Z/19/Z), is supported by the University of Bristol NIHR Biomedical Research Centre (BRC-1215-2001), the MRC Integrative Epidemiology Unit (MC_UU_00011/1) and works within the CRUK Integrative Cancer Epidemiology Programme (C18281/A29019). REM is a member of the MRC Integrative Epidemiology Unit at the University of Bristol funded by the MRC (MC_UU_00011/1). SH is supported by the UK National Institute for Health Research Academic Clinical Fellowship.

**National Survey of Health and Development (NSHD)**
We thank the NSHD participants for their continued support and lifelong contribution to the study. The NSHD is funded by the UKRI Medical Research Council [MC_UU_00019/1].

**Finland-United States Investigation of NIDDM Genetics (FUSION)**
We would like to thank the Finnish volunteers who generously participated in the FUSION study. Support for FUSION was provided by NIH grants R01-DK062370 (M.B.), R01-DK072193 (K.L.M.), and

intramural project number 1Z01-HG000024 (F.S.C.). Genome-wide genotyping was conducted by the Johns Hopkins University Genetic Resources Core Facility SNP Center at the Center for Inherited Disease Research (CIDR), with support from CIDR NIH contract no. N01-HG-65403.

**Metabolic Syndrome in Men (METSIM)**
The METSIM study was funded by the Academy of Finland (grant no.77299 and 124243)

**Minnesota Center for Twin and Family Research (MCTFR)**
Work conducted within the MCTFR was supported by grants from the National Institutes of Health DA044283, DA042755, DA037904, AA009367, DA005147, and DA036216.

**Johnston County Osteoarthritis Project (JoCoOA)**
The investigators wish to thank the staff and participants in the Johnston County Osteoarthritis Project, without whom this work would not be possible. The JoCoOA is supported in part by S043, S1734, & S3486 from the CDC/Association of Schools of Public Health; 5-P60-AR30701 & 5-P60-AR49465 from NIAMS/NIH, and U01 DP003206, U01 DP006266 from the CDC.

**Study of Health in Pomerania (SHIP)**
SHIP is part of the Community Medicine Research net of the University of Greifswald, Germany, which is funded by the Federal Ministry of Education and Research (grants no. 01ZZ9603, 01ZZ0103, and 01ZZ0403), the Ministry of Cultural Affairs as well as the Social Ministry of the Federal State of Mecklenburg-West Pomerania, and the network 'Greifswald Approach to Individualized Medicine (GANI_MED)' funded by the Federal Ministry of Education and Research (grant 03IS2061A). Genome-wide data have been supported by the Federal Ministry of Education and Research (grant no. 03ZIK012) and a joint grant from Siemens Healthineers, Erlangen, Germany and the Federal State of Mecklenburg-West Pomerania. The University of Greifswald is a member of the Caché Campus program of the InterSystems GmbH. SHIP was funded by the Federal Ministry of Education and Research grants no. 01ZZ9603, 01ZZ0103, 01ZZ0403, 03IS2061A, and 03ZIK012

**Age, gene/environment suceptibility Reykjavik study (AGES-RS)**
We want to thank the study participants for their participation. Age, Gene/Environment Susceptibility-Reykjavik Study (AGES) was funded by NIH contract N01-AG-1-2100 and HHSN271201200022C, the NIA Intramural Research Program, Hjartavernd (the Icelandic Heart Association), and the Althingi (the Icelandic Parliament).

**PROspective Study of Pravastatin in the Elderly at Risk for vascular disease (PROSPER)**
The PROSPER study was supported by an investigator initiated grant obtained from Bristol-Myers Squibb. Prof. Dr. J. W. Jukema is an Established Clinical Investigator of the Netherlands Heart Foundation (grant 2001 D 032). Support for genotyping was provided by the seventh framework program of the European commission (grant 223004) and by the Netherlands Genomics Initiative (Netherlands Consortium for Healthy Aging grant 050-060-810).

**deCODE**
We thank participants in deCODE cardiovascular- and obesity studies and collaborators for their cooperation.

**Generation Scotland (GS)**
We are grateful to all the families who took part, the general practitioners and the Scottish School of Primary Care for their help in recruiting them, and the whole Generation Scotland team, which includes interviewers, computer and laboratory technicians, clerical workers, research scientists, volunteers, managers, receptionists, healthcare assistants and nurses. Generation Scotland received core support from the Chief Scientist Office of the Scottish Government Health Directorates [CZD/16/6] and the Scottish Funding Council [HR03006] and is currently supported by the Wellcome Trust

[216767/Z/19/Z]. Genotyping of the GS:SFHS samples was carried out by the Genetics Core Laboratory at the Edinburgh Clinical Research Facility, University of Edinburgh, Scotland and was funded by the Medical Research Council UK and the Wellcome Trust (Wellcome Trust Strategic Award "STratifying Resilience and Depression Longitudinally" (STRADL) Reference 104036/Z/14/Z).

**Cardiovascular Health Improvement Project (CHIP)**
CJW is supported by NIH grant R-35-HL135824.

**The Trondelag Health Study (HUNT)**
The genetic investigations of the HUNT Study, is a collaboration between investigators from the HUNT study and University of Michigan Medical School and the University of Michigan School of Public Health. The K.G. Jebsen Center for Genetic Epidemiology is financed by Stiftelsen Kristian Gerhard Jebsen; Faculty of Medicine and Health Sciences, NTNU, Norwegian University of Science and Technology (NTNU) and Central Norway Regional Health Authority.

**GRAPHIC (Genetic Regulation of Arterial Pressure In humans in the Community)**
C.P.N is funded by the BHF (SP/16/4/32697). C.P.N., P.S.B. and N.J.S. are supported by the National Institute for Health Research (NIHR) Leicester Cardiovascular Biomedical Research Centre (BRC-1215-20010)

**Quebec Family Study (QFS)**
Thanks are expressed to the participants in the Québec Family Study and the staff of the Physical Activity Sciences Laboratory at Université Laval for their contribution in this study. The Quebec Family Study was funded by multiple grants from the Medical Research Council of Canada and the Canadian Institutes of Health Research. This work was supported by a team grant from the Canadian Institutes of Health Research (FRN-CCT-83028)

**Anglo-Scandinavian Cardiac Outcomes Trial UK (ASCOT UK)**
We thank all ASCOT trial participants, physicians, nurses, and practices in the participating countries for their important contribution to the study. In particular we thank Clare Muckian and David Toomey for their help in DNA extraction, storage, and handling. We also acknowledge support from the NIHR Barts Biomedical Research Centre and Queen Mary University of London, UK. This work was supported by Pfizer, New York, NY, USA, for the ASCOT study and the collection of the ASCOT DNA repository; by Servier Research Group, Paris, France; and by Leo Laboratories, Copenhagen, Denmark. HRW acknowledges funding from the NIHR Cardiovascular Biomedical Research Centre at Barts and QMUL. We also acknowledge support from the NIHR Barts Biomedical Research Centre and Queen Mary University of London, UK for genotyping. JR acknowledges funding from the European Union's Horizon 2020 research and innovation programme under the Marie Sklodowska-Curie grant agreement No 786833.

**British Genetics of Hypertension (BRIGHT) study**
The BRIGHT study is extremely grateful to all the patients who participated in the study and the BRIGHT nursing team. This work formed part of the research themes contributing to the translational research portfolio for the NIHR Barts Cardiovascular Biomedical Research Centre. The funders had no role in study design, data collection and analysis. This work was funded by the Medical Research Council of Great Britain (grant number: G9521010D).

**Estonian Biobank (EstBB)**
The EstBB gratefully acknowledges the contributions of the participants. Data analyses were carried out in part in the High-Performance Computing Center of University of Tartu. We would like to thank participants and support staff of Estonian Biobank. K.L was supported by Estonian Research Council grants PUT 1371, EMBO Installation grant 3573, and The European Regional Development Fund. This study was funded by the European Union through the European Regional Development Fund (Project

No. 2014-2020.4.01.15-0012 and Project No. 2014-2020.4.01.16-0125), by the European Union through Horizon 2020 grant no. 810645 and by the Estonian Research Council grants PUT (PRG687, PRG1291). T.E was supported by Estonian Research Council grant PUT (PRG1291). A.M was supported by the European Union through the European Regional Development Fund for the development of CoEs (Project No. 2014-2020.4.01.16-0125). K.P. was supported by the European Union through the European Regional Development Fund (Project No. 2014-2020.4.01.16-0030).

**Northern Finland Birth cohort 1966 (NFBC1966)**
The authors are grateful to the late professor Paula Rantakallio (launch of NFBC1966), the participants in the 31- and 46-years-old study and the NFBC project center (www.oulu.fi/nfbc). We thank all cohort members and researchers who participated in the 46 yrs study. We also wish acknowledge the work of the NFBC project center. NFBC1966 received financial support from University of Oulu Grant no. 24000692, Oulu University Hospital Grant no. 24301140, ERDF European Regional Development Fund Grant no. 539/2010 A31592, Academy of Finland ( 285547, EGEA).

**Austrian Stroke Prevention Study (ASPS), Austrian Stroke Prevention Family Study (ASPS-Fam)**
The Austrian Stroke Prevention Study (ASPS): The authors thank the staff and the participants for their valuable contributions. We thank Birgit Reinhart for her long-term administrative commitment, Elfi Hofer for the technical assistance at creating the DNA bank, Ing. Johann Semmler and Anita Harb for DNA sequencing and DNA analyses by TaqMan assays and Irmgard Poelzl for supervising the quality management processes after ISO9001 at the biobanking and DNA analyses. The research reported in this article was funded by the Austrian Science Fond (FWF) grant number P20545-P05 and P13180 and supported by the Austrian National Bank Anniversary Fund, P15435, the Austrian Ministry of Science under the aegis of the EU Joint Programme-Neurodegenerative Disease Research (JPND)-www.jpnd.eu and by the Austrian Science Fund P20545-B05. The Medical University of Graz supports the databank of the ASPS.

**Finnish Twin Cohort Study (FTC)**
We thank the twins for active participations and the staff of the study for their hard work. Phenotype and genotype data collection in the twin cohort has been supported by the Wellcome Trust Sanger Institute, the Broad Institute, ENGAGE – European Network for Genetic and Genomic Epidemiology, FP7-HEALTH-F4-2007, grant agreement number 201413, National Institute of Alcohol Abuse and Alcoholism (grants AA-12502, AA-00145, and AA-09203 to R J Rose and AA15416 and K02AA018755 to D M Dick) and the Academy of Finland (grants 100499, 205585, 118555, 141054, 264146, 308248, and 312073 to JKaprio). JKaprio acknowledges support by the Academy of Finland (grants 265240, 263278).

**TwinGene**
TwinGene is part of the Swedish Twin Registry, managed by Karolinska Institutet and receiving funding through the Swedish Research Council under the grant no 2017-00641.

**Ragama Health Study (RHS)**
The RHS was supported by a Grant from the National Center for Global Health and Medicine (NCGM).

**Cardio-metabolic Genome Epidemiology Network, GWAS1 (CAGE_GWAS1) and Cardio-metabolic Genome Epidemiology Network, Amagasaki Study (CAGE-Amagasaki)**
The CAGE Network studies were supported by grants for the Core Research for Evolutional Science and Technology (CREST) from the Japan Science Technology Agency; the Program for Promotion of Fundamental Studies in Health Sciences, National Institute of Biomedical Innovation Organization (NIBIO); and the Grant of National Center for Global Health and Medicine (NCGM).

**AugUR**

The AugUR study was supported by grants from the German Federal Ministry of Education and Research (BMBF 01ER1206, BMBF 01ER1507 to I.M.H.) and by the German Research Foundation (DFG; HE 3690/7-1 to I.M.H., BR 6028/2-1 to CB).

**Nurses' Health Study (NHS)**
The authors would like to thank the participants and staff of the Nurses Health Study for their valuable contributions. The Nurses' Health Study was supported by NIH grants UM1 CA186107, P01 CA87969 and R01 CA49449.

**Nurses' Health Study II (NHS II)**
The authors would like to thank the participants and staff of the Nurses' Health Study II for their valuable contributions. The Nurses' Health Study II was supported by NIH Grants U01 CA176726 and R01 CA67262.

**Health Professionals Follow-up Study (HPFS)**
We are grateful to the participants and staff of the Health Professionals Follow-Up Study for their valuable contributions. The Health Professionals Follow-up Study was supported by grants UM1CA167552, CA141298, and P01CA055075.

**Physicians Health Study (PHS)**
We are grateful to the participants and staff of the Physicians' Health Study for their valuable contributions. The Physicians Health Study was supported by grant CA141298.

**HyperGEN - Genetic Epidemiology of Hypertension Network**
HyperGEN was funded by NIH cooperative agreement grants HL54471, HL54472, HL54473, HL54495, HL54496, HL54509, HL54515.

**Cooperative health research in the region of Augsburg (KORA)**
The KORA study was initiated and financed by the Helmholtz Zentrum München –German Research Center for Environmental Health, which is funded by the German Federal Ministry of Education and Research (BMBF) and by the State of Bavaria.

**FINRISK1-7**
The FINRISK surveys have been mainly funded by budgetary funds from THL. Additional funding has been obtained from the Academy of Finland and various domestic foundations.

**GeneRISK**
The GeneRISK study was funded by Business Finland through the Personalized Diagnostics and Care program coordinated by SalWe Ltd (Grant No 3986/31/2013).
SR was supported by the Academy of Finland Center of Excellence in Complex Disease Genetics (Grant No 312062), the Finnish Foundation for Cardiovascular Research, the Sigrid Juselius Foundation and University of Helsinki HiLIFE Fellow and Grand Challenge grants. EW was supported by the Finnish innovation fund Sitra (EW) and Finska Läkaresällskapet.

**The Netherlands Twin Register (NTR)**
We thank all twins and family members for their participation. We also thank SURFsara for the support in using the Lisa Compute Cluster. See also: http://www.tweelingenregister.vu.nl/ for more information. Studies were approved by the Central Ethics Committee on Research Involving Human Subjects of the VU University Medical Centre, Amsterdam, an Institutional Review Board certified by the U.S. Office of Human Research Protections (IRB number IRB00002991 under Federal-wide Assurance- FWA00017598; IRB/institute codes, NTR 03-180). The Netherlands Twin Register acknowledges funding from the Netherlands Organization for Scientific research (NWO), including NWO-Grants NWO/SPI 56-464-14192 and 480-15-001/674: Netherlands Twin Registry Repository

and the Biobanking and Biomolecular Resources Research Infrastructure (BBMRI–NL, 184.021.007 and 184.033.111); Amsterdam Public Health (APH) and Neuroscience Campus Amsterdam (NCA); the European Community 7th Framework Program (FP7/2007-2013): ENGAGE (HEALTH-F4-2007-201413) and ACTION (9602768) and European Research Council (ERC-230374). We also acknowledge The Rutgers University Cell and DNA Repository cooperative agreement (NIMH U24 MH068457-06); the Collaborative Study of the Genetics of DZ twinning (NIH R01D0042157-01A1); the Developmental Study of Attention Problems in Young Twins (NIMH, RO1 MH58799-03); Major depression: stage 1 genome-wide association in population-based samples (MH081802); Determinants of Adolescent Exercise Behavior (NIDDK R01 DK092127-04); Grand Opportunity grants Integration of Genomics and Transcriptomics (NIMH 1RC2MH089951-01) and Developmental Trajectories of Psychopathology (NIMH 1RC2 MH089995); and the Avera Institute for Human Genetics, Sioux Falls, South Dakota (USA). DIB acknowledges her KNAW Academy Professor Award (PAH/6635).

**The Netherlands Study of Depression and Anxiety (NESDA)**
For NESDA, funding was obtained from the Netherlands Organization for Scientific Research (Geestkracht program grant 10-000-1002); the Center for Medical Systems Biology (CSMB, NWO Genomics), Biobanking and Biomolecular Resources Research Infrastructure (BBMRI-NL), VU University's Institutes for Health and Care Research (EMGO+) and Neuroscience Campus Amsterdam, University Medical Center Groningen, Leiden University Medical Center, National Institutes of Health (NIH, R01D0042157-01A, MH081802, Grand Opportunity grants 1RC2 MH089951 and 1RC2 MH089995). Part of the genotyping and analyses were funded by the Genetic Association Information Network (GAIN) of the Foundation for the National Institutes of Health. Computing was supported by BiG Grid, the Dutch e-Science Grid, which is financially supported by NWO. We

**BioMe**
RJFL is supported by the NIH (R01DK110113; R01DK075787; R01DK107786; R01HL142302; R01HG010297; R01DK124097; R01HL151152).

**iPSYCH**
Analyses of the iPSYCH cohort was conducted on the GenomeDenmark and Computerome High Performance Computing facilities. iPSYCH was supported by The Lundbeck Foundation, the Stanley Medical Research Institute, the Aarhus and Copenhagen universities and university hospitals; the research was conducted using the Danish National Biobank resource, supported by the Novo Nordisk Foundation.

**Cleveland Family Study**
The Cleveland Family Study has been supported by National Institutes of Health grants [R01-HL046380, KL2-RR024990, R35-HL135818, and R01-HL113338]. SR was funded by NHLBI R35HL135818; HL 046389; HL113338. BEC was funded by NIH grants [K01 HL135405, R03 HL154284].

**HyperGen-Axiom (Hypertension Genetic Epidemiology Network Axiom chip data)**
The HyperGen-Axiom was funded by NIH grant R01HL086718. XZ was funded by NHGRI HG011052.

**Framingham Heart Study**
This research was conducted in part using data and resources from the Framingham Heart Study of the National Heart Lung and Blood Institute of the National Institutes of Health and Boston University School of Medicine. The analyses reflect intellectual input and resource development from the Framingham Heart Study investigators participating in the SNP Health Association Resource (SHARe) project. This work was partially supported by the National Heart, Lung and Blood Institute's Framingham Heart Study (Contract Nos. N01-HC-25195 and HHSN268201500001I) and its contract with Affymetrix, Inc for genotyping services (Contract No. N02-HL-6-4278). A portion of this research utilized the Linux Cluster for Genetic Analysis (LinGA-II) funded by the Robert Dawson Evans

Endowment of the Department of Medicine at Boston University School of Medicine and Boston Medical Center. This research was partially supported by grant R01-DK122503 from the National Institute of Diabetes and Digestive and Kidney.

**Long Life Family Study (LLFS)**
LLFS acknowledges the contributions of the participants and of the study staff. LLFS was funded by NIA grants U01AG023746, U01AG023712, U01AG023749, U01AG023755, U01AG023744, and U19AG063893.

**Family Heart Study (FamHS)**
FamHS acknowledges the contributions of the participants and of the study staff. FamHS was funded by NIDDK R01-DK-089256 and NHLBI R01HL117078.

**Genetics of Lipid Lowering Drugs and Diet Network (GOLDN)**
GOLDN acknowledges the contributions of the participants and of the study staff. GOLDN was funded by the NHLBI grants R01 HL09135701, R01 HL091357 and R01 HL104135.

**Hellenic Isolated Cohorts MANOLIS (HELIC-MANOLIS)**
The MANOLIS cohort is named in honour of Manolis Giannakakis, 1978-2010. We thank the residents of Anogia and surrounding Mylopotamos villages, and of the Pomak villages, for taking part. The HELIC study has been supported by many individuals who have contributed to sample collection (including Antonis Athanasiadis, Olina Balafouti, Christina Batzaki, Georgios Daskalakis, Eleni Emmanouil, Chrisoula Giannakaki, Margarita Giannakopoulou, Anastasia Kaparou, Vasiliki Kariakli, Stella Koinaki, Dimitra Kokori, Maria Konidari, Hara Koundouraki, Dimitris Koutoukidis, Vasiliki Mamakou, Eirini Mamalaki, Eirini Mpamiaki, Maria Tsoukana, Dimitra Tzakou, Katerina Vosdogianni, Niovi Xenaki, Eleni Zengini), data entry (Thanos Antonos, Dimitra Papagrigoriou, Betty Spiliopoulou), sample logistics (Sarah Edkins, Emma Gray), genotyping (Robert Andrews, Hannah Blackburn, Doug Simpkin, Siobhan Whitehead), research administration (Anja Kolb-Kokocinski, Carol Smee, Danielle Walker) and informatics (Martin Pollard, Josh Randall). This work was funded by the Wellcome Trust (098051) and the European Research Council (ERC-2011-StG 280559-SEPI).

**Hellenic Isolated Cohorts Pomak (HELIC-Pomak)**
This work was funded by the Wellcome Trust (098051) and the European Research Council (ERC-2011-StG 280559-SEPI).

**Genetic Overlap between Metabolic and Psychiatric traits (GOMAP)**
We thank all participants for their important contribution. We are grateful to Georgia Markou, Laiko General Hospital Diabetes Centre, Maria Emetsidou and Panagiota Fotinopoulou, Hippokratio General Hospital Diabetes Centre, Athina Karabela, Dafni Psychiatric Hospital, Eirini Glezou and Marios Mangioros, Dromokaiteio Psychiatric Hospital, Angela Rentari, Harokopio University of Athens, and Danielle Walker, Wellcome Trust Sanger Institute. This work was funded by the Wellcome Trust (098051).

**Understanding Society: The UK Household Longitudinal Study (UKHLS)**
These data are from Understanding Society: The UK Household Longitudinal Study, which is led by the Institute for Social and Economic Research at the University of Essex and funded by the Economic and Social Research Council. The data were collected by NatCen and the genome wide scan data were analysed by the Wellcome Trust Sanger Institute. The Understanding Society DAC have an application system for genetics data and all use of the data should be approved by them. The application form is at: https://www.understandingsociety.ac.uk/about/health/data. The UK Household Longitudinal Study was funded by grants from the Economic & Social Research Council (ES/H029745/1) and the Wellcome Trust (WT098051).

**Nigeria Cohort**
The authors acknowledge the assistance of the research staff and participants in Igbo-Ora, Oyo State, Nigeria. Nigerian cohort study was supported by NIH grants R37-HL045508, R01-HL053353, R01-DK075787 and U01-HL054512.

**Maywood Cohort**
Maywood cohort study was supported by NIH grants R37-HL045508, R01-HL074166, R01-HL086718 and R01-HG003054.

**Epidemiologisch Preventief Onderzoek Zoetermeer (EPOZ)**
We are grateful to the contributions of the EPOZ study participants. HHHA was supported by ZonMW grant number 916.19.151.

**The Slim Initiative in Genomic Medicine for the Americas (SIGMA) T2D Consortium**
Acknowledges the contribution of participants and study staff. Additional funding from SIGMA, MCDS RO1 24799 NIHHLB ,CONACYT 2092,CONACYT M9303,CONACYT F677,CONACYT 251 M. J.M.Mercader is supported by American Diabetes Association Innovative and Clinical Translational Award 1-19-ICTS-068. The Mexico City Diabetes Study (part of The SIGMA consortium) has been funded by The National Heart, Lung and Blood Institutes RO1 HL 2479 , and by Consejo Naconal de Ciencia y Tecnologia CONACYT grants 2092, M9303, F677M9407, 251M 2005COI.

**Danish National Birth Cohort (DNBC)**
We are very grateful to all DNBC families who took part in the study. We would also like to thank everyone involved in data collection and biological material handling. The Danish National Birth Cohort (DNBC) is a result of major grants from the Danish National Research Foundation, the Danish Pharmacists' Fund, the Egmont Foundation, the March of Dimes Birth Defects Foundation, the Augustinus Foundation, and the Health Fund of the Danish Health Insurance Societies. The DNBC biobank is a part of the Danish National Biobank resource, which is supported by the Novo Nordisk Foundation. The generation of GWAS genotype data for the DNBC samples was carried out within the Gene Environment Association Studies (GENEVA) consortium with funding provided through the National Institutes of Health's Genes, Environment, and Health Initiative (U01HG004423; U01HG004446; U01HG004438). BF received support from an Oak Foundation fellowship and a Novo Nordisk Foundation grant (12955). XL received support from the Nordic Center of Excellence in Health-Related e-Sciences.

**Kita-Nagoya Genome Epidemiology Study (CAGE-KING)**
CAGE-KING was supported in part by Grants-in-Aid from MEXT (nos. 24390169, 16H05250, 15K19242, 16H06277, 19K19434, 20K10514, 21H03206) as well as by a grant from the Funding Program for Next-Generation World-Leading Researchers (NEXT Program, no. LS056).

**Indian Diabetes Prevention Study 3 (IDPP3)**
The IDPP3 gratefully acknowledges the participants of the volunteers in the study and the field staff who followed the volunteers on regular basis. Investigator support from the Council of Scientific and Industrial Research, Ministry of Science and Technology, Govt. of India, New delhi, India.

**Wellcome Genetic (WELLGEN)**
The WELLGEN is grateful and obliged to the study participants and other staff members who contributed to the study. Investigator support from the Council of Scientific and Industrial Research, Ministry of Science and Technology, Govt. of India, New delhi, India.

**Pune maternal Nutrition Study (PMNS)**

The PMNS gratefully acknowledges the contributions of the participants and of the study staff. Investigator support from the Council of Scientific and Industrial Research, Ministry of Science and Technology, Govt. of India, New delhi, India.

**The Fenland Study (Fenland)**
We are grateful to all the volunteers and to the General Practitioners and practice staff for assistance with recruitment. We thank the Fenland Study Investigators, Fenland Study Co-ordination team and the Epidemiology Field, Data and Laboratory teams. The Fenland Study (10.22025/2017.10.101.00001)is funded by the Medical Research Council (MC_UU_12015/1). We further acknowledge support for genotyping from the Medical Research Council (MC_PC_13046).

**COPSAC2000**
We express our deepest gratitude to the children and families of the COPSAC2000 cohort for all their support and commitment. All funding received by COPSAC is listed on www.copsac.com. The Lundbeck Foundation (Grant no. R16-A1694); The Ministry of Health (Grant no. 903516); Danish Council for Strategic Research (Grant no. 0603-00280B), and The Capital Region Research Foundation have provided core support to the COPSAC research center.

**Inter99**
The Inter99 was initiated by Torben Jørgensen (PI), Knut Borch-Johnsen (co-PI), Hans Ibsen and Troels F. Thomsen. The steering committee comprises the former two and Charlotta Pisinger. The study was financially supported by research grants from the Danish Research Council, the Danish Centre for Health Technology Assessment, Novo Nordisk Inc., Research Foundation of Copenhagen County, Ministry of Internal Affairs and Health, the Danish Heart Foundation, the Danish Pharmaceutical Association, the Augustinus Foundation, the Ib Henriksen Foundation, the Becket Foundation, and the Danish Diabetes Association. Novo Nordisk Foundation Center for Basic Metabolic Research is an independent Research Center, based at the University of Copenhagen, Denmark and partially funded by an unconditional donation from the Novo Nordisk Foundation (www.cbmr.ku.dk) (Grant number NNF18CC0034900).

**Health**
The Health2006 was financially supported by grants from the Velux Foundation; The Danish Medical Research Council, Danish Agency for Science, Technology and Innovation; The Aase and Ejner Danielsens Foundation; ALK-Abello A/S, Hørsholm, Denmark, and Research Centre for Prevention and Health, the Capital Region of Denmark. Health2008 was supported by the Timber Merchant Vilhelm Bang's Foundation, the Danish Heart Foundation (Grant number 07-10-R61-A1754-B838-22392F), and the Health Insurance Foundation (Helsefonden) (Grant number 2012B233).

**DanFund**
This study was supported by TrygFonden (7-11-0213), the Lundbeck Foundation (R155-2013-14070), Novo Nordisk Foundation (NNF15OC0015896).

**SDC**
Novo Nordisk Foundation Center for Basic Metabolic Research is an independent Research Center, based at the University of Copenhagen, Denmark and partially funded by an unconditional donation from the Novo Nordisk Foundation (www.cbmr.ku.dk) (Grant number NNF18CC0034900).

**Vejle Biobank**
The Vejle Diabetes Biobank was supported by The Danish Research Council for Independent Research and by Region of Southern Denmark.

**Heinz Nixdorf Recall Study (HNR)**

The HNR study is indebted to all study participants and to both the dedicated personnel of the study center of the Heinz Nixdorf Recall study and to the investigative group. The HNR study thanks the Heinz Nixdorf Foundation for their generous support of this study. Parts of the study were also supported by the German Research Council (DFG) [DFG project: EI 969/2-3, ER 155/6-1;6-2, HO 3314/2-1;2-2;2-3;4-3, INST 58219/32-1, JO 170/8-1, KN 885/3-1, PE 2309/2-1, SI 236/8-1;9-1;10-1,], the German Ministry of Education and Science [BMBF project: 01EG0401, 01GI0856, 01GI0860, 01GS0820_WB2-C, 01ER1001D, 01GI0205], the Ministry of Innovation, Science, Research and Technology, North Rhine-Westphalia (MIWFT-NRW).

**LBR (Leicester BioResource)**
C.P.N is funded by the BHF (SP/16/4/32697). C.P.N., P.S.B. and N.J.S. are supported by the National Institute for Health Research (NIHR) Leicester Cardiovascular Biomedical Research Centre (BRC-1215-20010).

**Nijmegen Biomedical Study (NBS)**
The Nijmegen Biomedical Study is a population-based survey conducted at the Department for Health Evidence and the Department of Laboratory Medicine of the Radboud university medical center. Principal investigators of the Nijmegen Biomedical Study are L.A.L.M. Kiemeney, A.L.M. Verbeek, D.W. Swinkels en B. Franke.

**Taiwan Type 2 Diabetes Study Consortium (TWT2D)**
The TWT2D gratefully acknowledges the contributions of the participants and of the study staff. The TWT2D was funded by Academia Sinica.

**Cebu Longitudinal Health and Nutrition Survey (CLHNS)**
We thank the entire staff of the Office of Population Studies (OPS) Foundation in Cebu for their long-term work on the CLHNS. The CLHNS was supported by US National Institutes of Health grants DK078150, TW005596, HL085144; pilot funds from RR020649, ES010126, and DK056350; and the Office of Population Studies Foundation in Cebu. Additional support for data analysis was provided by US NIH R01DK072193. CNS was supported by American Heart Association postdoctoral fellowships 15POST24470131 and 17POST33650016.

**China Health and Nutrition Survey (CHNS)**
We thank the participants, researchers and staff who contributed to the China Health and Nutrition Survey (CHNS). We are grateful to research grant funding from the National Institute for Health (NIH), the Eunice Kennedy Shriver National Institute of Child Health and Human Development (NICHD) for R01 HD30880, National Institute on Aging (NIA) for R01 AG065357, National Institute of Diabetes and Digestive and Kidney Diseases (NIDDK) for R01DK104371 and R01HL108427, the NIH Fogarty grant D43 TW009077 since 1989, and the China-Japan Friendship Hospital, Ministry of Health for support for CHNS 2009, Chinese National Human Genome Center at Shanghai since 2009, and Beijing Municipal Center for Disease Prevention and Control since 2011. We thank the National Institute for Nutrition and Health, China Center for Disease Control and Prevention. Additional support for data analysis was provided by US NIH R01DK072193. CNS was supported by American Heart Association postdoctoral fellowships 15POST24470131 and 17POST33650016.

**The Raine Study**
The authors are grateful to the Raine Study participants and their families, and to the Raine Study team for cohort coordination and data collection. The authors gratefully acknowledge the NHMRC for their long-term funding to the study over the last 30 years and also the following institutes for providing funding for Core Management of the Raine Study: The University of Western Australia (UWA), Curtin University, Women and Infants Research Foundation, Telethon Kids Institute, Edith Cowan University, Murdoch University, The University of Notre Dame Australia and The Raine Medical Research Foundation. This work was supported by resources provided by the Pawsey Supercomputing Centre

with funding from the Australian Government and Government of Western Australia. The Raine Study was supported by the National Health and Medical Research Council of Australia [grant numbers 572613, 403981, 1059711, 1027449, 1044840, 1021858], the Canadian Institutes of Health Research [grant number MOP-82893], and WA Health, Government of Western Australia (WADOH) [Future Health WA G06302]. Funding was also generously provided by Safe Work Australia.

**Exeter 10,000 Study (EXTEND)**
EXTEND is supported by the National Institute for Health Research Exeter Clinical Research Facility.

**Invecchiare in Chianti (InCHIANTI)**
The InCHIANTI study baseline (1998-2000) was supported as a "targeted project" (ICS110.1/RF97.71) by the Italian Ministry of Health and in part by the U.S. National Institute on Aging (Contracts: 263 MD 9164 and 263 MD 821336); the InCHIANTI Follow-up 1 (2001-2003) was funded by the U.S. National Institute on Aging (Contracts: N.1-AG-1-1 and N.1-AG-1-2111).

**Relationship between Insulin Sensitivity and Cardiovascular disease Study (RISC)**
The RISC Study was supported by European Union grant QLG1-CT-2001-01252 and AstraZeneca.

**China Kadoorie Biobank (CKB)**
CKB acknowledges the contribution of participants in the study and the members of the survey teams in each of the 10 regional centres, as well as the project development and management teams based at Beijing, Oxford, and the 10 regional centres. China Kadoorie Biobank was supported as follows: Baseline survey: Hong Kong Kadoorie Charitable Foundation; long-term follow-up, study support and maintenance: UK Wellcome Trust (212946/Z/18/Z, 202922/Z/16/Z, 104085/Z/14/Z, 088158/Z/09/Z), National Natural Science Foundation of China (91846303), and National Key Research and Development Program of China (2016YFC 0900500, 0900501, 0900504, 1303904). DNA extraction and genotyping: GlaxoSmithKline, UK Medical Research Council (MC_PC_13049, MC-PC-14135). The UK Medical Research Council (MC_UU_00017/1, MC_UU_12026/2, MC_U137686851), Cancer Research UK (C16077/A29186, C500/A16896) and the British Heart Foundation (CH/1996001/9454), provide core funding to the Clinical Trial Service Unit and Epidemiological Studies Unit at Oxford University for the project.

**DIACORE**
Cohort recruiting and management was funded by the KfH Stiftung Präventivmedizin e.V. (Carsten A. Böger). Genome-wide genotyping was funded the Else Kröner-Fresenius-Stiftung (2012_A147), the KfH Stiftung Präventivmedizin and the University Hospital Regensburg. The Deutsche Forschungsgemeinschaft (DFG, German Research Foundation) supported this work – Project-ID 387509280 – SFB 1350 (Subproject C6 to I.M.H.) and Iris Heid and Carsten Böger received funding by DFG BO 3815/4-1. This project has received funding from the Innovative Medicines Initiative 2 Joint Undertaking under grant agreement No 115974. This Joint Undertaking receives support from the European Union's Horizon 2020 research and innovation programme and EFPIA with JDRF (BEAT-DKD).

**Jackson Heart Study (JHS)**
The Jackson Heart Study (JHS) is supported and conducted in collaboration with Jackson State University (HHSN268201800013I), Tougaloo College (HHSN268201800014I), the Mississippi State Department of Health (HHSN268201800015I) and the University of Mississippi Medical Center (HHSN268201800010I, HHSN268201800011I and HHSN268201800012I) contracts from the National Heart, Lung, and Blood Institute (NHLBI) and the National Institute on Minority Health and Health Disparities (NIMHD). The authors also wish to thank the staffs and participants of the JHS. The views expressed in this manuscript are those of the authors and do not necessarily represent the views of the National Heart, Lung, and Blood Institute; the National Institutes of Health; or the U.S. Department of Health and Human Services. The project described was supported by the National Center for Advancing

Translational Sciences, National Institutes of Health, through Grant KL2TR002490 (LMR). LMR was also supported by T32HL129982.

**MrOS Gothenburg**
MrOS in Sweden is supported by the Swedish Research Council, the Swedish Foundation for Strategic Research, the ALF/LUA research grant in Gothenburg, the Lundberg Foundation, the Knut and Alice Wallenberg Foundation, the Torsten Soderberg Foundation, and the Novo Nordisk Foundation.

**Whitehall II (WHII)**
We thank all the participating civil service departments and their welfare, personnel, and establishment officers; the British Occupational Health and Safety Agency; the British Council of Civil Service Unions; all participating civil servants in the Whitehall II study; and all members of the Whitehall II study team. The Whitehall II Study team comprises research scientists, statisticians, study coordinators, nurses, data managers, administrative assistants, and data entry staff, all of whom made this study possible. The Whitehall II study is supported by grants from the Wellcome Trust (221854/Z/20/Z), the UK Medical Research Council (MRC, K013351 and R024227), the US National Institute on Aging (R01AG056477; R01AG034454), and the British Heart Foundation (32334). Kivimaki is supported by grants from the Wellcome Trust (221854/Z/20/Z), the UK Medical Research Council (MRC, K013351 and R024227), the US National Institute on Aging (R01AG056477; R01AG034454) and Academy of Finland (329202). Kumari is supported by grants from ESRC (ES/S007253/1 ; ES/T002611/1; ES/T014083/1).

**Multiethnic cohort - African American Breast Cancer (MEC-AABC); Multiethnic cohort - African American Prostate Cancer (MEC-AAPC); Multiethnic cohort - Latina American Breast Cancer (MEC-LABC); Multiethnic cohort - Latino American Prostate Cancer (MEC-LAPC)**
The Multiethnic Cohort (MEC) is a population-based prospective cohort study including approximately 215,000 men and women from Hawaii and California. All participants were 45-75 years of age at baseline, and primarily of 5 ancestries: Japanese Americans, African Americans, European Americans, Hispanic/Latinos, and Native Hawaiians. (PMIDs: 10695593; 23449381) MEC was funded by the National Cancer Institute in 1993 to examine lifestyle risk factors and genetic susceptibility to cancer. All eligible cohort members completed baseline and follow-up questionnaires.

**Beijing Eye Study**
Supported by Beijing Municipal of Health Reform and Development Project #2019-4.

**Blue Mountains Eye Study (BMES) / Lions Eye Institute (LEI)**
The Westmead/Sydney samples were collected in three studies that were supported by the National Health and Medical Research Council (NHMRC), Australia: Grant IDs 974159, 211069, 457349 and 512423 supported the Blue Mountains Eye Study that provided population-based controls; Grant ID 302010 supported the Cataract Surgery and Risk of Age-related Macular Degeneration study that provided clinic-based early and late AMD cases and controls; and Grant ID 571013 supported the Genes and Environment in late AMD study that provided clinic-based late AMD cases. The NHMRC had no role in the design or conduct of these studies. DAM is supported by an NHMRC Practitioner Fellowship (GNT1154518). AWH is supported by an NHMRC Practitioner Fellowship (GNT1103329).

**Children's Hospital of Philadelphia - CHOP**
This research was financially supported by an Institute Development Award from the Children's Hospital of Philadelphia, a Research Development Award from the Cotswold Foundation, the Childrens´s Hospital of Philadelphia Endowed Chair in Genomic Research, the Daniel B. Burke Endowed Chair for Diabetes Research and NIH grant R01 HD056465. The authors thank the network of primary care clinicians and the patients and families for their contribution to this project and to clinical research facilitated by the Pediatric Research Consortium (PeRC) at The Children's Hospital of Philadelphia. R. Chiavacci, E. Dabaghyan, A. (Hope) Thomas, K. Harden, A. Hill, C. Johnson-Honesty, C. Drummond, S.

Harrison, F. Salley, C. Gibbons, K. Lilliston, C. Kim, E. Frackelton, F. Mentch, G. Otieno, K. Thomas, C. Hou, K. Thomas and M.L. Garris provided expert assistance with genotyping and/or data collection and management. The authors would also like to thank S. Kristinsson, L.A. Hermannsson and A. Krisbjörnsson of Raförninn ehf for extensive software design and contributions.

**Genetic Park of Cilento and Vallo di Diano Project (Cilento)**
The Cilento gratefully acknowledges the population for their participation and the study staff. The Cilento was funded by grants from the Italian Ministry of Universities (IDF SHARID ARS01_01270), the Assessorato Ricerca Regione Campania (POR CAMPANIA 2000/2006 MISURA 3.16).

**Epihealth**
Funding from the Swedish research council.

**Lifelines Cohort Study**
The authors wish to acknowledge the services of the Lifelines Cohort Study, the contributing research centers delivering data to Lifelines, and all the study participants. The Lifelines Biobank initiative has been made possible by funding from the Dutch Ministry of Health, Welfare and Sport, the Dutch Ministry of Economic Affairs, the University Medical Center Groningen (UMCG the Netherlands), University of Groningen and the Northern Provinces of the Netherlands. The generation and management of GWAS genotype data for the Lifelines Cohort Study is supported by the UMCG Genetics Lifelines Initiative (UGLI). UGLI is partly supported by a Spinoza Grant from NWO, awarded to Cisca Wijmenga.

**Genetic Epidemiology Network of Arteriopathy (GENOA)**
Genotyping was performed at the Mayo Clinic (Stephen Turner, Mariza de Andrade, Julie Cunningham) and was made possible by the University of Texas Health Sciences Center (Eric Boerwinkle, Megan Grove-Gaona). We would also like to thank the families that participated in the GENOA study. Support for GENOA was provided by the National Heart, Lung and Blood Institute (U01 HL054457, U01 HL054464, U01 HL054481, R01 HL119443, and R01 HL087660) of the National Institutes of Health.

**Health and Retirement Study (HRS)**
HRS is supported by the National Institute on Aging (NIA U01AG009740). The genotyping was funded separately by the National Institute on Aging (RC2 AG036495, RC4 AG039029). Our genotyping was conducted by the NIH Center for Inherited Disease Research (CIDR) at Johns Hopkins University. Genotyping quality control and final preparation of the data were performed by the Genetics Coordinating Center at the University of Washington and the School of Public Health at the University of Michigan. Additional funding from US National Institutes of Health grants U01AG009740, RC2 AG036495, RC4 AG039029.

**Genes for Good**
Genes for Good gratefully acknowledges the generous contributions from its research participants and the work of its administrative staff for making the study possible. Genes for Good was funded through University of Michigan discretionary funds.

**Korea Association REsource (KARE)**
This study was conducted with bioresources from National Biobank of Korea, the Korea Disease Control and Prevention Agency, Republic of Korea. The Korean Association Resource (KARE) was supported by grants from National Institute of Health, Republic of Korea (4845–301, 4851–302, 4851–307) and intramural grants from the Korea National Institute of Health (2019-NG-053-02).

**CROATIA-KORCULA**
The CROATIA studies gratefully acknowledge the contributions of the participants and of the many students and members of staff who supported the field work, including but not limited to those from The University of Split and Zagreb Medical Schools, the Institute for Anthropological Research in

Zagreb, the Croatian Institute for Public Health. We are grateful to administrative teams in Croatia and Edinburgh. We acknowledge the Helmholtz Zentrum München, Neuherberg, Germany where the array-genotyping was performed. CROATIA studies were funded by grants from the Medical Research Council UK (https://mrc.ukri.org/), in particular MRC Programme Grants to the Human Genetics Unit, "QTL in health and disease", currently MC_UU_00007/10 , the Republic of Croatia Ministry of Science, Education and Sports research grants to I.R. (108-1080315-0302), and European Commission Framework 6 project EUROSPAN (Contract No. LSHG-CT-2006-018947). VV is supported by the MRC Programme Grant to the Human Genetics Unit, "QTL in health and disease", currently MC_UU_00007/10. O.Po. is supported by the Croatian National Center of Research Excellence in Personalized Healthcare (grant number KK.01.1.1.01.0010); and the Center of Competence in Molecular Diagnostics (KK.01.2.2.03.0006).

**CROATIA-VIS**

The CROATIA studies gratefully acknowledge the contributions of the participants and of the many students and members of staff who supported the field work, including but not limited to those from The University of Split and Zagreb Medical Schools, the Institute for Anthropological Research in Zagreb, the Croatian Institute for Public Health. We are grateful to administrative teams in Croatia and Edinburgh and acknowledge staff at the Wellcome Trust Clinical Research Facility, Edinburgh, UK for performing the array-genotyping. A.F. W. was supported by the MRC (https://mrc.ukri.org/) Programme Grant to the Human Genetics Unit, "QTL in health and disease".

**Myocardial Infarction Genetics Consortium (MIGEN)**

Funding support was provided by grants 1K08HG010155 and 1U01HG011719 (to A.V.K.) from the National Human Genome Research Institute

**Sardinia**

Funding support from Intramural Research Program of the NIH, National Institute on Aging, with contracts N01-AG-1-2109 and HHSN271201100005C. Sardinian Autonomous Region (L.R. no. 7/2009) grant cRP3-154. Fondazione di Sardegna (grant U1301.2015/AI.1157. BE Prat. 2015-1651).

**TwinsUK (TUK)**

We are grateful to the twins who took part in TwinsUK and the whole TwinsUK team, which includes academic researchers, clinical staff, laboratory technicians, administrative staff and research managers. TwinsUK receives funding from the Wellcome Trust (212904/Z/18/Z), Medical Research Council (AIMHY; MR/M016560/1) and European Union (H2020 contract #733100). TwinsUK and M.M. are supported by the National Institute for Health Research (NIHR)-funded BioResource, Clinical Research Facility and Biomedical Research Centre based at Guy's and St Thomas' NHS Foundation Trust in partnership with King's College London. P.C. is funded by the European Union (H2020 contract #733100).

**MESA (Multi-Ethnic Study of Atherosclerosis)**

MESA and the MESA SHARe projects are conducted and supported by the National Heart, Lung, and Blood Institute (NHLBI) in collaboration with MESA investigators. Support for MESA is provided by contracts 75N92020D00001, HHSN268201500003I, N01-HC-95159, 75N92020D00005, N01-HC-95160, 75N92020D00002, N01-HC-95161, 75N92020D00003, N01-HC-95162, 75N92020D00006, N01-HC-95163, 75N92020D00004, N01-HC-95164, 75N92020D00007, N01-HC-95165, N01-HC-95166, N01-HC-95167, N01-HC-95168, N01-HC-95169, UL1-TR-000040, UL1-TR-001079, and UL1-TR-001420. Funding for SHARe genotyping was provided by NHLBI Contract N02-HL-64278. Genotyping was performed at Affymetrix (Santa Clara, California, USA) and the Broad Institute of Harvard and MIT (Boston, Massachusetts, USA) using the Affymetrix Genome-Wide Human SNP Array 6.0. Also supported in part by the National Center for Advancing Translational Sciences, CTSI grant UL1TR001881, and the National Institute of Diabetes and Digestive and Kidney Disease Diabetes

Research Center (DRC) grant DK063491 to the Southern California Diabetes Endocrinology Research Center.

**Mexican-American Coronary Artery Disease (MACAD) Study**
This research was supported in part by the Mexican-American Coronary Artery Disease (MACAD) National Heart, Lung, and Blood Institute, contracts R01-HL088457, R01-HL-60030.

**Hypertension and Insulin Resistance (HTN-IR)**
Supported in part by the Hypertension and Insulin Resistance (HTN-IR) contracts R01-HL067974, R01-HL-55005, R01-HL 067974.

**The Rare Variants for Hypertension in Taiwan Chinese (THRV)**
The Rare Variants for Hypertension in Taiwan Chinese (THRV) is supported by the National Heart, Lung, and Blood Institute (NHLBI) grant (R01HL111249) and its participation in TOPMed is supported by an NHLBI supplement (R01HL111249-04S1). THRV is a collaborative study between Washington University in St. Louis, The Lundquist Institute at Harbor UCLA, University of Texas in Houston, Taichung Veterans General Hospital, Taipei Veterans General Hospital, Tri-Service General Hospital, National Health Research Institutes, National Taiwan University, and Baylor University. THRV is based (substantially) on the parent SAPPHIRe study, along with additional population-based and hospital-based cohorts. SAPPHIRe was supported by NHLBI grants (U01HL54527, U01HL54498) and Taiwan funds, and the other cohorts were supported by Taiwan funds.

**Taiwan U.S. Diabetic Retinopathy Study (TUDR)**
This study was supported by the National Eye Institute of the National Institutes of Health (EY014684) and ARRA Supplement (EY014684-03S1, -04S1), the National Institute of Diabetes and Digestive and Kidney Disease grant DK063491 to the Southern California Diabetes Endocrinology Research Center, the Eye Birth Defects Foundation Inc., the National Science Council, Taiwan (NSC 98-2314-B-075A-002-MY3) and the Taichung Veterans General Hospital, Taichung, Taiwan (TCVGH-1003001C).

**AFNET**
We would like to thank all investigators nad patients participating in the EAST - AFNEt 4 trial. EAST - AFNET 4 was supported by AFNET, EHRA, German Centre for Cardiovascular Research (DZHK), German heart Foundation (DSF), Sanofi, and Abbott.

**BioVU**
Vanderbilt University Medical Center's BioVU projects are supported by numerous sources: institutional funding, private agencies, and federal grants. These include the NIH funded Shared Instrumentation Grant S10OD017985 and S10RR025141; CTSA grants UL1TR002243, UL1TR000445, and UL1RR024975. Genomic data have also been supported by investigator-led projects that include U01HG004798, R01NS032830, RC2GM092618, P50GM115305, U01HG006378, U19HL065962, R01HD074711; and additional funding sources listed at https://victr.vumc.org/biovu-funding/.

**DFAust**
Dr Fatkin is supported by the Victor Chang Cardiac Research Institute, NSW Health, and the National health and Medical Research Council of Australia (1186500).

**CATHGEN**
Dr. McGarrah is supported by NIH grant 5K08HL135275.

**GENAF**
The GENAF study is funded by the Norwegian Research Council with a Mobility Grant (240149) and Young Research Talent grant (287086); the South-Eastern Health Authorities with a PhD-grant

(2019122); Vestre Viken Hospital Trust with a PhD-grant; afib.no - the Norwegian Atrial Fibrillation Research Network; "Indremedisinsk Forskningsfond" at Bærum Hospital.

**GGAF**
The GGAF is supported by funding to the 5 sources that form GGAF. The AF RISK study is supported by the Netherlands Heart Foundation (grant NHS2010B233), and the Center for Translational Molecular Medicine. Both the Young-AF and Biomarker-AF studies are supported by funding from the University Medical Center Groningen. The GIPS-III trial was supported by grant 95103007 from ZonMw, the Netherlands Organization for Health Research and Development. The PREVEND study is supported by the Dutch Kidney Foundation (grant E0.13) and the Netherlands Heart Foundation (grant NHS2010B280). Prof.Dr. Rienstra acknowledges support from the Netherlands Cardiovascular Research Initiative: an initiative supported by the Netherlands Heart Foundation, CVON 2014–9: "Reappraisal of Atrial Fibrillation: interaction between hyperCoagulability, Electrical remodelling, and Vascular destabilization in the progression of AF (RACE V)".

**GRADE**
Supported by NIH-NHLBI R01 HL77398.

**INSPIRE_AF**
Dr Cutler is supported by funding from the Dell Loy Hansen Heart Foundation.

**JHU_AF**
Dr. Nazarian is supported by grants from the US NIH/NHLBI, as well as Biosense Webster, ImriCor, and ADAS software.

**MGH_AF**
Dr. Ellinor is supported by the Fondation Leducq (14CVD01), the NIH (1RO1HL092577, and K24HL105780) and the American Heart Association (18SFRN34110082).

**MGH Stroke**
Dr. Lubitz is supported by NIH grant 1R01HL139731 and American Heart Association 18SFRN34250007.

**MPP-AF, MPP-echo**
J. Gustav Smith was supported by grants from the Swedish Heart-Lung Foundation (2019-0526), the Swedish Research Council (2017-02554), the European Research Council (ERC-STG-2015-679242), Skåne University Hospital, governmental funding of clinical research within the Swedish National Health Service, a generous donation from the Knut and Alice Wallenberg foundation to the Wallenberg Center for Molecular Medicine in Lund, and funding from the Swedish Research Council (Linnaeus grant Dnr 349-2006-237, Strategic Research Area Exodiab Dnr 2009-1039) and Swedish Foundation for Strategic Research (Dnr IRC15-0067) to the Lund University Diabetes Center.

**The Cooperative Health Research In South Tyrol Study (CHRIS)**
The CHRIS study was funded by the Department of Innovation, Research, and University of the Autonomous Province of Bolzano-South Tyrol. Full acknowledgements for the CHRIS study are reported here: https://translational-medicine.biomedcentral.com/articles/10.1186/s12967-015-0704-9#Sec33.

**The Atherosclerosis Risk in Communities Study (ARIC)**
ARIC is a prospective epidemiologic study conducted in four U.S. communities (Williams OD. The Atherosclerosis Risk in Communities (Aric) Study - Design and Objectives. American Journal of Epidemiology. 1989;129(4):687-702. PubMed PMID: WOS:A1989T805200005.). It is designed to investigate the causes of atherosclerosis and its clinical outcomes, and variation in cardiovascular risk

factors, medical care, and disease by race, gender, location, and date. ARIC includes two parts: the Cohort Component and the Community Surveillance Component. The Cohort Component began in 1987, and each ARIC field center randomly selected and recruited a cohort sample of approximately 4,000 individuals aged 45-64 from a defined population in their community, to receive extensive examinations, including medical, social, and demographic data. Follow-up currently occurs semi-annually, by telephone, to maintain contact and to assess health status of the cohort. ARIC: The Atherosclerosis Risk in Communities study has been funded in whole or in part with Federal funds from the National Heart, Lung, and Blood Institute, National Institutes of Health, Department of Health and Human Services (contract numbers HHSN268201700001I, HHSN268201700002I, HHSN268201700003I, HHSN268201700004I and HHSN268201700005I), R01HL087641, R01HL086694; National Human Genome Research Institute contract U01HG004402; and National Institutes of Health contract HHSN268200625226C. The authors thank the staff and participants of the ARIC study for their important contributions. Infrastructure was partly supported by Grant Number UL1RR025005, a component of the National Institutes of Health and NIH Roadmap for Medical Research.

**The Hispanic Community Health Study / Study of Latinos (HCHS/SOL)**

The Hispanic Community Health Study / Study of Latinos (HCHS/SOL) is a multi-center study of Hispanic/Latino populations with the goal of determining the role of acculturation in the prevalence and development of diseases, and to identify other traits that impact Hispanic/Latino health. 2 The study is sponsored by the National Heart, Lung, and Blood Institute (NHLBI) and other institutes, centers, and offices of the National Institutes of Health (NIH). Recruitment began in 2006 with a target population of 16,000 persons of Cuban, Puerto Rican, Dominican, Mexican or Central/South American origin. Household sampling was employed as part of the study design. Participants were recruited through four sites affiliated with San Diego State University, Northwestern University in Chicago, Albert Einstein College of Medicine in Bronx, New York, and the University of Miami. Researchers from seven academic centers provided scientific and logistical support. Study participants who were self-identified Hispanic/Latino and aged 18-74 years underwent extensive psycho-social and clinical assessments during 2008-2011. A re-examination of the HCHS/SOL cohort is conducted during 2015-2017. Annual telephone follow-up interviews are ongoing since study inception to determine health outcomes of interest.(dbGaP study accession number: phs000555). HCHS/SOL: Primary funding support to Dr. North and colleagues is provided by U01HG007416. Additional support was provided via R01DK101855 and 15GRNT25880008. The HCHS/SOL study was carried out as a collaborative study supported by contracts from the National Heart, Lung, and Blood Institute (NHLBI) to the University of North Carolina (N01-HC65233), University of Miami (N01-HC65234), Albert Einstein College of Medicine (N01-HC65235), Northwestern University (N01-HC65236), and San Diego State University (N01-HC65237). The following Institutes/Centers/Offices contribute to the HCHS/SOL through a transfer of funds to the NHLBI: NIMHD, National Institute on Deafness and Other Communication Disorders, National Institute of Dental and Craniofacial Research, National Institute of Diabetes and Digestive and Kidney Diseases, National Institute of Neurological Disorders and Stroke, NIH Institution-Office of Dietary Supplements.

**The Health, Aging and Body Composition Study (Health ABC Study)**

This research was supported by National Institute on Aging (NIA) Contracts N01-AG-6-2101; N01-AG-6-2103; N01-AG-6-2106; NIA grant R01-AG028050, and NINR grant R01-NR012459. This research was funded in part by the Intramural Research Program of the NIH, National Institute on Aging.

**The Religious Orders Study and the Rush Memory and Aging Project batch 1 (ROSMAP1)**

The ROSMAP are grateful to the participants of the studies, and the faculty and staff of the Rush Alzheimer's Disease Center. Funding from NIA grant P30AG10161, P30AG72975, R01AG17917, RF1AG15819, R01AG30146, U01AG46152, U01AG61256; Translational Genomics Research Institute.

**Healthy Aging in Neighborhooeds of Diversity across the Life Span (HANDLS)**

We thank all HANDLS participants for their commitment to and participation in the study. HANDLS is funded by the NIA Intrmaural Research Program Project Number AG000513.

**Bone Mineral Density in Childhood Study (BMDCS)**
We appreciate the dedication of the BMDCS study participants and their families, and the support of Dr. Karen Winer, Scientific Director of this effort. This study was funded by R01 HD58886, R01 HD100406, the Eunice Kennedy Shriver National Institute of Child Health and Human Development (NICHD) contracts (N01-HD-1-3228, -3329, -3330, -3331, -3332, -3333), and the CTSA program Grant 8 UL1 TR000077. S.F.A.G. is supported by the Daniel B. Burke Endowed Chair for Diabetes Research, R01 HD056465 and R01 HG010067.

**Shanghai breast cancer study (SBCS)**
The generation and management of GWAS genotype data for the SBCS was supported by R01CA64277 and R01CA15847.

**Shanghai Women's Health Study(SWHS)**
The generation and management of GWAS genotype data for the SWHS was supported by UM1CA182910 and R01CA148677.

**Singapore Chinese Health Study - Coronary Artery Disease (SCHS CAD)**
We thank Siew-Hong Low of the National University of Singapore for supervising the field work of the Singapore Chinese Health Study and the Ministry of Health in Singapore for assistance with the identification of AMI cases via database linkages. We also acknowledge the founding, longstanding principal investigator of the Singapore Chinese Health Study, Mimi C. Yu. The Singapore Chinese Health Study (SCHS) was supported by the U.S. National Institutes of Health (Grant Numbers R01CA144034 and UM1 CA182876), and by the Singapore National Medical Research Council(Grant Number 1270/2010). Genotyping of the SCHS CAD subset was funded by the HUJ-CREATE Programme of the National Research Foundation, Singapore (Project Number 370062002).

**Rotterdam Study I, II, III**
The generation and management of GWAS genotype data for the Rotterdam Study (RS I, RS II, RS III) was executed by the Human Genotyping Facility of the Genetic Laboratory of the Department of Internal Medicine, Erasmus MC, Rotterdam. We thank Pascal Arp, Mila Jhamai, Marijn Verkerk, Lizbeth Herrera and Marjolein Peters, PhD, and Carolina Medina-Gomez, PhD, for their help in creating the GWAS database and for the creation and analysis of imputed data. The GWAS datasets are supported by the Netherlands Organisation of Scientific Research NWO Investments (nr. 175.010.2005.011, 911-03-012), the Genetic Laboratory of the Department of Internal Medicine, Erasmus MC, the Research Institute for Diseases in the Elderly (014-93-015; RIDE2), the Netherlands Genomics Initiative (NGI)/Netherlands Organisation for Scientific Research (NWO) Netherlands Consortium for Healthy Aging (NCHA), project nr. 050-060-810.

**BPROOF**
The authors gratefully thank all study participants, and all co-workers who helped to succeed this trial, especially P.H. in 't Veld, M. Hillen-Tijdink, A. Nicolaas-Merkus, N. Pliester, S. Oliai Araghi, and S. Smits. They also thank Prof. Dr. H.A.P. Pols for obtaining funding. The generation and management of GWAS genotype data for theBPROOF study was executed by the Human Genotyping Facility of the Genetic Laboratory of the Department of Internal Medicine, Erasmus MC, Rotterdam. We thank Carolina Medina, Ph.D. and Jard de Vries for their help in creating the GWAS database and for the creation and analysis of imputed data. B-PROOF is supported and funded by The Netherlands Organization for Health Research and Development (ZonMw, Grant 6130.0031), The Hague; unrestricted grant from NZO (Dutch Dairy Association), Zoetermeer; NCHA (Netherlands Consortium Healthy Ageing) Leiden/ Rotterdam; Ministry of Economic Affairs, Agriculture and Innovation (project KB-15-004-003), the

Hague; Wageningen University, Wageningen; VU University Medical Center, Amsterdam; Erasmus MC, Rotterdam.

**Diabetes Genetics Initiative (DGI)**
The contribution of the Botnia and Skara research teams is gratefully acknowledged. DGI (Principal investigators Leif Groop and Tiinamaija Tuomi) and the Botnia Study is supported by the Sigrid Juselius Foundation, The Folkhalsan Research Foundation, Nordic Center of Excellence in Disease Genetics, EU (EXGENESIS), Finnish Diabetes Research Foundation, Foundation for Life and Health in Finland, Finnish Medical Society, Helsinki University Central Hospital Research Foundation, Perklén Foundation, Ollqvist Foundation, Narpes Health Care Foundation as well as the Municipal Heath Care Center and Hospital in Jakobstad and Health Care Centers in Vasa, Narpes and Korsholm. The work in Malmö, Sweden, was also funded by a Linné grant from the Swedish Research Council (349-2006-237). Additionally supported by NIH R01DK075787. The GIANT Consortium is supported by R01DK075787 to J.N.H.

**Prostate cancer Genome-wide Association Study to Uncover Susceptibility loci (PEGASUS)**
PEGASUS gratefully acknowledges contributions of the study participants. The content of this publication does not necessarily reflect the views or policies of the Department of Health and Human Services nor does mention of trade names, commercial products, or organization indicate endorsement by the U.S. Government. PEGASUS was supported by the Intramural Research Program of the Division of Cancer Epidemiology and Genetics, National Cancer Institute, NIH (ZIA CP010152-20).

**NESCOG**
NESCOG research was part of Science Live, the innovative research program of science center NEMO that enables scientists to carry out their research using NEMO visitors as volunteers. The Netherlands Organization for Scientific Research (NWO) Division for the Social Sciences (MaGW) provided funding for this research throughVIDI 016-065-318 to D.P.

**UK Biobank**
We are grateful to UK Biobank participants. This research has been conducted using the UK Biobank Resource under project 12505. LY was funded by the Australian Research Council (DE200100425). PMV was funded by the Australian Research Council (FL180100072) and the Australian National Health and Medical Research Council (Grant #1113400). JY was funded by the Australian National Health and Medical Research Council (Grant #1113400) and Westlake Education Foundation. R.E.M. was supported by US National Institutes of Health (NIH) grant K25 HL150334. PRL was funded by US NIH grant DP2 ES030554, a Burroughs Wellcome Fund Career Award at the Scientific Interfaces, the Next Generation Fund at the Broad Institute of MIT and Harvard, and a Sloan Research Fellowship.

**Erasmus Rucphen Family study (ERF)**
We are grateful to all study participants and their relatives, general practitioners and neurologists for their contributions and to P. Veraart for her help in genealogy, J. Vergeer for the supervision of the laboratory work, both S.J. van der Lee and A. van der Spek for collection of the follow-up data and P. Snijders M.D. for his help in data collection of both baseline and follow-up data. Erasmus Rucphen Family (ERF) was supported by the Consortium for Systems Biology (NCSB), both within the framework of the Netherlands Genomics Initiative (NGI)/Netherlands Organisation for Scientific Research (NWO). ERF study as a part of EUROSPAN (European Special Populations Research Network) was supported by European Commission FP6 STRP grant number 018947 (LSHG-CT-2006-01947) and also received funding from the European Community's Seventh Framework Programme (FP7/2007-2013)/grant agreement HEALTH-F4-2007-201413 by the European Commission under the programme "Quality of Life and Management of the Living Resources" of 5th Framework Programme (No. QLG2-CT-2002-01254) as well as FP7 project EUROHEADPAIN (nr 602633).

**Breast Cancer Association Consortium (BCAC)**

BCAC is funded by the European Union's Horizon 2020 Research and Innovation Programme (grant numbers 634935 and 633784 for BRIDGES and B-CAST respectively), and the PERSPECTIVE I&I project, funded by the Government of Canada through Genome Canada and the Canadian Institutes of Health Research, the Ministère de l'Économie et de l'Innovation du Québec through Genome Québec, the Quebec Breast Cancer Foundation. The EU Horizon 2020 Research and Innovation Programme funding source had no role in study design, data collection, data analysis, data interpretation or writing of the report. Additional funding for BCAC is provided via the Confluence project which is funded with intramural funds from the National Cancer Institute Intramural Research Program, National Institutes of Health. Genotyping of the OncoArray was funded by the NIH Grant U19 CA148065, and Cancer UK Grant C1287/A16563 and the PERSPECTIVE project supported by the Government of Canada through Genome Canada and the Canadian Institutes of Health Research (grant GPH-129344) and, the Ministère de l'Économie, Science et Innovation du Québec through Genome Québec and the PSRSIIRI-701 grant, and the Quebec Breast Cancer Foundation. Funding for iCOGS came from: the European Community's Seventh Framework Programme under grant agreement n° 223175 (HEALTH-F2-2009-223175) (COGS), Cancer Research UK (C1287/A10118, C1287/A10710, C12292/A11174, C1281/A12014, C5047/A8384, C5047/A15007, C5047/A10692, C8197/A16565), the National Institutes of Health (CA128978) and Post-Cancer GWAS initiative (1U19 CA148537, 1U19 CA148065 and 1U19 CA148112 - the GAME-ON initiative), the Department of Defence (W81XWH-10-1-0341), the Canadian Institutes of Health Research (CIHR) for the CIHR Team in Familial Risks of Breast Cancer, and Komen Foundation for the Cure, the Breast Cancer Research Foundation, and the Ovarian Cancer Research Fund.

**1958BC-WTCCC and 1958BC-T1DGC.**

This work made use of data and samples generated by the 1958 Birth Cohort (NCDS), which is managed by the Centre for Longitudinal Studies at the UCL Institute of Education, The authors are deeply grateful to the 1958 birth cohort participants for their longstanding commitment and support, and to all staff for cohort coordination and data collection. The management of the 1958 Birth Cohort is funded by the Economic and Social Research Council (grant number ES/M001660/1). Access to these resources was enabled via the 58READIE Project funded by Wellcome Trust and Medical Research Council (grant numbers WT095219MA and G1001799). DNA collection was funded by MRC grant G0000934 and cell-line creation by Wellcome Trust grant 068545/Z/02. This study makes use of data generated by the Wellcome Trust Case-Control Consortium. A full list of investigators who contributed to generation of the data is available from the Wellcome Trust Case-Control Consortium website. Funding for the project was provided by the Wellcome Trust under the award 076113. This research used resources provided by the Type 1 Diabetes Genetics Consortium, a collaborative clinical study sponsored by the National Institute of Diabetes and Digestive and Kidney Diseases (NIDDK), National Institute of Allergy and Infectious Diseases, National Human Genome Research Institute, National Institute of Child Health and Human Development, and Juvenile Diabetes Research Foundation International (JDRF) and supported by U01 DK062418. M.I.M. was a Wellcome Investigator and NIHR Senior Investigator. This work was supported by: NIDDK (U01-DK105535) and Wellcome (090532, 098381, 106130, 203141, 212259).

**Indian Diabetes Consortium (INDICO)**

The INDICO gratefully acknowledges the contributions of the participants and the study staff. INDICO was supported by Council of Scientific and Industrial Research (CSIR), Government of India through Centre for Cardiovascular and Metabolic Disease Research (CARDIOMED) project (Grant no. BSC0122); also partially funded by Department of Science and Technology, Government of India through PURSE II CDST/SR/PURSE PHASE II/11 provide to Jawaharlal Nehru University, New Delhi, INDIA.

**GoDARTS**

We are grateful to all the participants in this study, the general practitioners, the Scottish School of Primary Care for their help in recruiting the participants, and to the whole team, which includes interviewers, computer and laboratory technicians, clerical workers, research scientists, volunteers, managers, receptionists, and nurses. The study complies with the Declaration of Helsinki. We acknowledge the support of the Health Informatics Centre, University of Dundee for managing and supplying the anonymised data and NHS Tayside, the original data owner. M.I.M. was a Wellcome Investigator and NIHR Senior Investigator. This work was supported by: NIDDK (U01-DK105535) and Wellcome (090532, 098381, 106130, 203141, 212259). The Wellcome Trust United Kingdom Type 2 Diabetes Case Control Collection (GoDARTS) was funded by The Wellcome Trust (072960/Z/03/Z, 084726/Z/08/Z, 084727/Z/08/Z, 085475/Z/08/Z, 085475/B/08/Z). GoDARTS was funded by the Wellcome Trust (084727/Z/08/Z, 085475/Z/08/Z, 085475/B/08/Z) and as part of the EU IMI-SUMMIT program.

**Oxford Biobank**
This research was funded by the National Institute for Health Research Oxford Biomedical Research Centre and the British Heart Foundation [RG/17/1/32663].

**SORBS**
We thank all those who participated in the study. Sincere thanks are given to Dr. Knut Krohn (University of Leipzig) for the genotyping support. This work was supported by grants from the Deutsche Forschungsgemeinschaft (DFG, German Research Foundation – Projektnummer 209933838 – SFB 1052; B03, C01; SPP 1629 TO 718/2- 1).

**The Finnish Cardiovascular Study (FINCAVAS)**
The authors thank the staff of the Department of Clinical Physiology for collecting the exercise test data. The Finnish Cardiovascular Study (FINCAVAS) has been financially supported by the Competitive Research Funding of the Tampere University Hospital (Grant 9M048 and 9N035), the Finnish Cultural Foundation, the Finnish Foundation for Cardiovascular Research, the Emil Aaltonen Foundation, Finland, the Tampere Tuberculosis Foundation, EU Horizon 2020 (grant 755320 for TAXINOMISIS and grant 848146 for To Aition), and the Academy of Finland grant 322098.

**Special Turku Coronary Risk Factor Intervention Project (STRIP) parents**
We thank the study participants and their families as well as the research group who collected the data. STRIP was supported by Academy of Finland (grants 206374, 251360 and 276861); Juho Vainio Foundation; Finnish Cardiac Research Foundation; Finnish Cultural Foundation; Finnish Ministry of Education and Culture; Sigrid Juselius Foundation; Yrjö Jahnsson Foundation; C.G. Sundell Foundation; Special Governmental Grants for Health Sciences Research, Turku University Hospital; Foundation for Pediatric Research; and Turku University Foundation.

**Young Finns Study (YFS)**
We thank the teams that collected data at all measurement time points; the persons who participated as both children and adults in these longitudinal studies; and biostatisticians Irina Lisinen, Johanna Ikonen, Noora Kartiosuo, Ville Aalto, and Jarno Kankaanranta for data management and statistical advice. The Young Finns Study has been financially supported by the Academy of Finland: grants 322098, 286284, 134309 (Eye), 126925, 121584, 124282, 129378 (Salve), 117787 (Gendi), and 41071 (Skidi); the Social Insurance Institution of Finland; Competitive State Research Financing of the Expert Responsibility area of Kuopio, Tampere and Turku University Hospitals (grant X51001); Juho Vainio Foundation; Paavo Nurmi Foundation; Finnish Foundation for Cardiovascular Research ; Finnish Cultural Foundation; The Sigrid Juselius Foundation; Tampere Tuberculosis Foundation; Emil Aaltonen Foundation; Yrjö Jahnsson Foundation; Signe and Ane Gyllenberg Foundation; Diabetes Research Foundation of Finnish Diabetes Association; EU Horizon 2020 (grant 755320 for TAXINOMISIS and grant 848146 for To Aition); European Research Council (grant 742927 for MULTIEPIGEN project); Tampere University Hospital Supporting Foundation and Finnish Society of Clinical Chemistry.

**GerMiFS**
The GerMiFS gratefully acknowledges the contributions of the participants and of the study staff.

**SR - Silk Road**

The authors gratefully acknowledge the subjects from the SR cohort. This study is part of the scientific activities carried out within the scientific expedition Marcopolo 2010. We thank the Terramadre organization and the Terramadre communities who participated in the project. This study was funded by the Italian Ministry of Health—RC 01/21 to MPC and D70-RESRICGIROTTO to GG.

**INGI-FVG (INGI-Friuli Venezia Giulia)**
The authors gratefully acknowledge the subjects from the INGI-FVG cohort. This study was funded by 5 per mille 2015 senses, Genetics of senses and related diseases, CUP: C92F17003560001, to PG.

**GIANT Consortium banners**

**23andMe, Inc.**
The following members of the 23andMe Research Team contributed to this study: Stella Aslibekyan, Elizabeth Babalola, Robert K. Bell, Jessica Bielenberg, Katarzyna Bryc, Emily Bullis, Daniella Coker, Gabriel Cuellar Partida, Devika Dhamija, Sayantan Das, Sarah L. Elson, Teresa Filshtein, Kipper Fletez-Brant, Pierre Fontanillas, Will Freyman, Pooja M. Gandhi, Karl Heilbron, Barry Hicks, David A. Hinds, Ethan M. Jewett, Katelyn Kukar, Keng-Han Lin, Maya Lowe, Jey C. McCreight, Matthew H. McIntyre, Steven J. Micheletti, Meghan E. Moreno, Joanna L. Mountain, Priyanka Nandakumar, Elizabeth S. Noblin, Jared O'Connell, Aaron A. Petrakovitz, G. David Poznik, Morgan Schumacher, Anjali J. Shastri, Janie F. Shelton, Suyash Shringarpure, Vinh Tran, Joyce Y. Tung, Xin Wang, Wei Wang, Catherine H. Weldon, Peter Wilton, Alejandro Hernandez, Corinna Wong, Christophe Toukam Tchakouté.

**VA Million Veteran Program (MVP)**
The MVP gratefully acknowledges the contributions of the participants and of the study staff. This research is based on data from the Million Veteran Program, Office of Research and Development, Veterans Health Administration, and was supported by award # I01-BX004821 (MVP 001, PIs Peter Wilson and Kelly Cho). This publication does not represent the views of the Department of Veteran Affairs or the United States Government. YVS was partly supported by NR013520 and DK125187 from NIH. SR was supported by US Department of Veterans Affairs grant IK2-CX001907 and a Webb-Waring Biomedical Research Award from the Boettcher Foundation. MVP acknowledgements and banner authors include:
MVP Executive Committee
Co-Chair: J. Michael Gaziano, M.D., M.P.H., VA Boston Healthcare System, 150 S. Huntington Avenue, Boston, MA 02130
Co-Chair: Sumitra Muralidhar, Ph.D., US Department of Veterans Affairs, 810 Vermont Avenue NW, Washington, DC 20420
Rachel Ramoni, D.M.D., Sc.D., Chief VA Research and Development Officer, US Department of Veterans Affairs, 810 Vermont Avenue NW, Washington, DC 20420
Jean Beckham, Ph.D., Durham VA Medical Center, 508 Fulton Street, Durham, NC 27705
Kyong-Mi Chang, M.D., Philadelphia VA Medical Center, 3900 Woodland Avenue, Philadelphia, PA 19104
Philip S. Tsao, Ph.D., VA Palo Alto Health Care System, 3801 Miranda Avenue, Palo Alto, CA 94304
James Breeling, M.D., Ex-Officio, US Department of Veterans Affairs, 810 Vermont Avenue NW, Washington, DC 20420
Grant Huang, Ph.D., Ex-Officio, US Department of Veterans Affairs, 810 Vermont Avenue NW, Washington, DC 20420
Juan P. Casas, M.D., Ph.D., Ex-Officio, VA Boston Healthcare System, 150 S. Huntington Avenue, Boston, MA 02130

MVP Program Office
Sumitra Muralidhar, Ph.D., US Department of Veterans Affairs, 810 Vermont Avenue NW, Washington, DC 20420

Jennifer Moser, Ph.D., US Department of Veterans Affairs, 810 Vermont Avenue NW, Washington, DC 20420

MVP Recruitment/Enrollment

MVP Cohort Management Director/Recruitment/Enrollment Director, Boston – Stacey B. Whitbourne, Ph.D.; Jessica V. Brewer, M.P.H., VA Boston Healthcare System, 150 S. Huntington Avenue, Boston, MA 02130

VA Central Biorepository, Boston – Mary T. Brophy M.D., M.P.H.; Donald E. Humphries, Ph.D.; Luis E. Selva, Ph.D., VA Boston Healthcare System, 150 S. Huntington Avenue, Boston, MA 02130

MVP Informatics, Boston – Nhan Do, M.D.; Shahpoor (Alex) Shayan, M.S., VA Boston Healthcare System, 150 S. Huntington Avenue, Boston, MA 02130

MVP Data Operations/Analytics, Boston – Kelly Cho, M.P.H., Ph.D., VA Boston Healthcare System, 150 S. Huntington Avenue, Boston, MA 02130

Director of Regulatory Affairs – Lori Churby, B.S., VA Palo Alto Health Care System, 3801 Miranda Avenue, Palo Alto, CA 94304

MVP Coordinating Centers

- Cooperative Studies Program Clinical Research Pharmacy Coordinating Center, Albuquerque – Todd Connor, Pharm.D.; Dean P. Argyres, B.S., M.S., New Mexico VA Health Care System, 1501 San Pedro Drive SE, Albuquerque, NM 87108
- Genomics Coordinating Center, Palo Alto – Philip S. Tsao, Ph.D., VA Palo Alto Health Care System, 3801 Miranda Avenue, Palo Alto, CA 94304
- MVP Boston Coordinating Center, Boston - J. Michael Gaziano, M.D., M.P.H., VA Boston Healthcare System, 150 S. Huntington Avenue, Boston, MA 02130
- MVP Information Center, Canandaigua – Brady Stephens, M.S., Canandaigua VA Medical Center, 400 Fort Hill Avenue, Canandaigua, NY 14424

MVP Science

Saiju Pyarajan Ph.D., VA Boston Healthcare System, 150 S. Huntington Avenue, Boston, MA 02130

Philip S. Tsao, Ph.D., VA Palo Alto Health Care System, 3801 Miranda Avenue, Palo Alto, CA 94304

Data Core - Kelly Cho, M.P.H, Ph.D., VA Boston Healthcare System, 150 S. Huntington Avenue, Boston, MA 02130

VA Informatics and Computing Infrastructure (VINCI) – Scott L. DuVall, Ph.D., VA Salt Lake City Health Care System, 500 Foothill Drive, Salt Lake City, UT 84148

Data and Computational Sciences – Saiju Pyarajan, Ph.D., VA Boston Healthcare System, 150 S. Huntington Avenue, Boston, MA 02130

Statistical Genetics – Elizabeth Hauser, Ph.D., Durham VA Medical Center, 508 Fulton Street, Durham, NC 27705; Yan Sun, Ph.D., Atlanta VA Medical Center, 1670 Clairmont Road, Decatur, GA 30033; Hongyu Zhao, Ph.D., West Haven VA Medical Center, 950 Campbell Avenue, West Haven, CT 06516

Current MVP Local Site Investigators

Atlanta VA Medical Center (Peter Wilson, M.D.), 1670 Clairmont Road, Decatur, GA 30033

Bay Pines VA Healthcare System (Rachel McArdle, Ph.D.), 10,000 Bay Pines Blvd Bay Pines, FL 33744

Birmingham VA Medical Center (Louis Dellitalia, M.D.), 700 S. 19th Street, Birmingham AL 35233

Central Western Massachusetts Healthcare System (Kristin Mattocks, Ph.D., M.P.H.), 421 North Main Street, Leeds, MA 01053

Cincinnati VA Medical Center (John Harley, M.D., Ph.D.), 3200 Vine Street, Cincinnati, OH 45220

Clement J. Zablocki VA Medical Center (Jeffrey Whittle, M.D., M.P.H.), 5000 West National Avenue, Milwaukee, WI 53295

VA Northeast Ohio Healthcare System (Frank Jacono, M.D.), 10701 East Boulevard, Cleveland, OH 44106

Durham VA Medical Center (Jean Beckham, Ph.D.), 508 Fulton Street, Durham, NC 27705

Edith Nourse Rogers Memorial Veterans Hospital (John Wells., Ph.D.), 200 Springs Road, Bedford, MA 01730

Edward Hines, Jr. VA Medical Center (Salvador Gutierrez, M.D.), 5000 South 5th Avenue, Hines, IL 60141
Veterans Health Care System of the Ozarks (Kathrina Alexander, M.D.), 1100 North College Avenue, Fayetteville, AR 72703
Fargo VA Health Care System (Kimberly Hammer, Ph.D.), 2101 N. Elm, Fargo, ND 58102
VA Health Care Upstate New York (James Norton, Ph.D.), 113 Holland Avenue, Albany, NY 12208
New Mexico VA Health Care System (Gerardo Villareal, M.D.), 1501 San Pedro Drive, S.E. Albuquerque, NM 87108
VA Boston Healthcare System (Scott Kinlay, M.B.B.S., Ph.D.), 150 S. Huntington Avenue, Boston, MA 02130
VA Western New York Healthcare System (Junzhe Xu, M.D.), 3495 Bailey Avenue, Buffalo, NY 14215-1199
Ralph H. Johnson VA Medical Center (Mark Hamner, M.D.), 109 Bee Street, Mental Health Research, Charleston, SC 29401
Columbia VA Health Care System (Roy Mathew, M.D.), 6439 Garners Ferry Road, Columbia, SC 29209
VA North Texas Health Care System (Sujata Bhushan, M.D.), 4500 S. Lancaster Road, Dallas, TX 75216
Hampton VA Medical Center (Pran Iruvanti, D.O., Ph.D.), 100 Emancipation Drive, Hampton, VA 23667
Richmond VA Medical Center (Michael Godschalk, M.D.), 1201 Broad Rock Blvd., Richmond, VA 23249
Iowa City VA Health Care System (Zuhair Ballas, M.D.), 601 Highway 6 West, Iowa City, IA 52246-2208
Eastern Oklahoma VA Health Care System (River Smith, Ph.D.), 1011 Honor Heights Drive, Muskogee, OK 74401
James A. Haley Veterans' Hospital (Stephen Mastorides, M.D.), 13000 Bruce B. Downs Blvd, Tampa, FL 33612
James H. Quillen VA Medical Center (Jonathan Moorman, M.D., Ph.D.), Corner of Lamont & Veterans Way, Mountain Home, TN 37684
John D. Dingell VA Medical Center (Saib Gappy, M.D.), 4646 John R Street, Detroit, MI 48201
Louisville VA Medical Center (Jon Klein, M.D., Ph.D.), 800 Zorn Avenue, Louisville, KY 40206
Manchester VA Medical Center (Nora Ratcliffe, M.D.), 718 Smyth Road, Manchester, NH 03104
Miami VA Health Care System (Ana Palacio, M.D., M.P.H.), 1201 NW 16th Street, 11 GRC, Miami FL 33125
Michael E. DeBakey VA Medical Center (Olaoluwa Okusaga, M.D.), 2002 Holcombe Blvd, Houston, TX 77030
Minneapolis VA Health Care System (Maureen Murdoch, M.D., M.P.H.), One Veterans Drive, Minneapolis, MN 55417
N. FL/S. GA Veterans Health System (Peruvemba Sriram, M.D.), 1601 SW Archer Road, Gainesville, FL 32608
Northport VA Medical Center (Shing Shing Yeh, Ph.D., M.D.), 79 Middleville Road, Northport, NY 11768
Overton Brooks VA Medical Center (Neeraj Tandon, M.D.), 510 East Stoner Ave, Shreveport, LA 71101
Philadelphia VA Medical Center (Darshana Jhala, M.D.), 3900 Woodland Avenue, Philadelphia, PA 19104
Phoenix VA Health Care System (Samuel Aguayo, M.D.), 650 E. Indian School Road, Phoenix, AZ 85012
Portland VA Medical Center (David Cohen, M.D.), 3710 SW U.S. Veterans Hospital Road, Portland, OR 97239
Providence VA Medical Center (Satish Sharma, M.D.), 830 Chalkstone Avenue, Providence, RI 02908
Richard Roudebush VA Medical Center (Suthat Liangpunsakul, M.D., M.P.H.), 1481 West 10th Street, Indianapolis, IN 46202
Salem VA Medical Center (Kris Ann Oursler, M.D.), 1970 Roanoke Blvd, Salem, VA 24153
San Francisco VA Health Care System (Mary Whooley, M.D.), 4150 Clement Street, San Francisco, CA 94121
South Texas Veterans Health Care System (Sunil Ahuja, M.D.), 7400 Merton Minter Boulevard, San Antonio, TX 78229
Southeast Louisiana Veterans Health Care System (Joseph Constans, Ph.D.), 2400 Canal Street, New Orleans, LA 70119
Southern Arizona VA Health Care System (Paul Meyer, M.D., Ph.D.), 3601 S 6th Avenue, Tucson, AZ 85723
Sioux Falls VA Health Care System (Jennifer Greco, M.D.), 2501 W 22nd Street, Sioux Falls, SD 57105

St. Louis VA Health Care System (Michael Rauchman, M.D.), 915 North Grand Blvd, St. Louis, MO 63106
Syracuse VA Medical Center (Richard Servatius, Ph.D.), 800 Irving Avenue, Syracuse, NY 13210
VA Eastern Kansas Health Care System (Melinda Gaddy, Ph.D.), 4101 S 4th Street Trafficway, Leavenworth, KS 66048
VA Greater Los Angeles Health Care System (Agnes Wallbom, M.D., M.S.), 11301 Wilshire Blvd, Los Angeles, CA 90073
VA Long Beach Healthcare System (Timothy Morgan, M.D.), 5901 East 7th Street Long Beach, CA 90822
VA Maine Healthcare System (Todd Stapley, D.O.), 1 VA Center, Augusta, ME 04330
VA New York Harbor Healthcare System (Peter Liang, M.D., M.P.H.), 423 East 23rd Street, New York, NY 10010
VA Pacific Islands Health Care System (Daryl Fujii, Ph.D.), 459 Patterson Rd, Honolulu, HI 96819
VA Palo Alto Health Care System (Philip Tsao, Ph.D.), 3801 Miranda Avenue, Palo Alto, CA 94304-1290
VA Pittsburgh Health Care System (Patrick Strollo, Jr., M.D.), University Drive, Pittsburgh, PA 15240
VA Puget Sound Health Care System (Edward Boyko, M.D.), 1660 S. Columbian Way, Seattle, WA 98108-1597
VA Salt Lake City Health Care System (Jessica Walsh, M.D.), 500 Foothill Drive, Salt Lake City, UT 84148
VA San Diego Healthcare System (Samir Gupta, M.D., M.S.C.S.), 3350 La Jolla Village Drive, San Diego, CA 92161
VA Sierra Nevada Health Care System (Mostaqul Huq, Pharm.D., Ph.D.), 975 Kirman Avenue, Reno, NV 89502
VA Southern Nevada Healthcare System (Joseph Fayad, M.D.), 6900 North Pecos Road, North Las Vegas, NV 89086
VA Tennessee Valley Healthcare System (Adriana Hung, M.D., M.P.H.), 1310 24th Avenue, South Nashville, TN 37212
Washington DC VA Medical Center (Jack Lichy, M.D., Ph.D.), 50 Irving St, Washington, D. C. 20422
W.G. (Bill) Hefner VA Medical Center (Robin Hurley, M.D.), 1601 Brenner Ave, Salisbury, NC 28144
White River Junction VA Medical Center (Brooks Robey, M.D.), 163 Veterans Drive, White River Junction, VT 05009
William S. Middleton Memorial Veterans Hospital (Prakash Balasubramanian, M.D.), 2500 Overlook Terrace, Madison, WI 53705

**DiscovEHR (DiscovEHR and MyCode Community Health Initiative)**
**Regeneron Genetics Center Banner Author List and Contribution Statements**
All authors/contributors are listed in alphabetical order.

RGC Management and Leadership Team
Goncalo Abecasis, Aris Baras, Michael Cantor, Giovanni Coppola, Andrew Deubler, Aris Economides, Luca A. Lotta, John D. Overton, Jeffrey G. Reid, Alan Shuldiner, Katia Karalis and Katherine Siminovitch

Sequencing and Lab Operations
Christina Beechert, Caitlin Forsythe, Erin D. Fuller, Zhenhua Gu, Michael Lattari, Alexander Lopez, John D. Overton, Thomas D. Schleicher, Maria Sotiropoulos Padilla, Louis Widom, Sarah E. Wolf, Manasi Pradhan, Kia Manoochehri, Ricardo H. Ulloa.

Genome Informatics
Xiaodong Bai, , Suganthi Balasubramanian, Boris Boutkov, Gisu Eom, Lukas Habegger, Alicia Hawes, Shareef Khalid, Olga Krasheninina, Rouel Lanche, Adam J. Mansfield, Evan K. Maxwell, Mona Nafde, Sean O'Keeffe, Max Orelus, Razvan Panea, Tommy Polanco, Ayesha Rasool, Jeffrey G. Reid, William Salerno, Jeffrey C. Staples,

Clinical Informatics:
Michael Cantor, Dadong Li, Deepika Sharma

Research Program Management
Marcus B. Jones, Jason Mighty, and Lyndon J. Mitnaul

**eMERGE (Electronic Medical Records and Genomics Network) banner author list:**
Murray Brilliant, Wendy Chung, Paul Crane, Damien Croteau-Chonka, Josh Denny, Todd Edwards, Geoff Hayes, Scott Hebbring, George Hripsak, Krzysztok Kiryluk, Terrie Kitchner, Iftikhar Kullo, Bahram Namjou, Peggy Peissig, Ning Shang, Digna Velez Edwards, Chunhua Weng

**The Prostate Cancer Association Group to Investigate Cancer Associated Alterations in the Genome (PRACTICAL) Consortium**

Funding & Acknowledgements
Genotyping of the OncoArray was funded by the US National Institutes of Health (NIH) [U19 CA 148537 for ELucidating Loci Involved in Prostate cancer SuscEptibility (ELLIPSE) project and X01HG007492 to the Center for Inherited Disease Research (CIDR) under contract number HHSN268201200008I]. Additional analytic support was provided by NIH NCI U01 CA188392 (PI: Schumacher).

The PRACTICAL consortium was supported by Cancer Research UK Grants C5047/A7357, C1287/A10118, C1287/A16563, C5047/A3354, C5047/A10692, C16913/A6135, European Commission's Seventh Framework Programme grant agreement n° 223175 (HEALTH-F2-2009-223175), and The National Institute of Health (NIH) Cancer Post-Cancer GWAS initiative grant: No. 1 U19 CA 148537-01 (the GAME-ON initiative).

We would also like to thank the following for funding support: The Institute of Cancer Research and The Everyman Campaign, The Prostate Cancer Research Foundation, Prostate Research Campaign UK (now PCUK), The Orchid Cancer Appeal, Rosetrees Trust, The National Cancer Research Network UK, The National Cancer Research Institute (NCRI) UK. We are grateful for support of NIHR funding to the NIHR Biomedical Research Centre at The Institute of Cancer Research and The Royal Marsden NHS Foundation Trust.

This study would not have been possible without the contributions of the following:
Coordination team, bioinformatician and genotyping centres:
Genotyping at CCGE, Cambridge: Caroline Baines and Don Conroy

Additional funding and acknowledgments from studies in PRACTICAL:
Information of the PRACTICAL consortium can be found at
http://practical.icr.ac.uk/

Acknowledgments for studies contributing to PRACTICAL Consortium
AHS
This work was supported by the Intramural Research Program of the NIH, National Cancer Institute, Division of Cancer Epidemiology and Genetics (Z01CP010119).

ATBC
The ATBC Study is supported by the Intramural Research Program of the U.S. National Cancer Institute, National Institutes of Health, Department of Health and Human Services.

APCB
The Australian Prostate Cancer BioResource (APCB) was supported by The National Health and Medical Research Council, Enabling Grant [614296] and the Prostate Cancer Foundation of Australia.
The Australian Prostate Cancer BioResource (APCB) would like to acknowledge and sincerely thank the urologists, pathologists, coordinators, data managers, nurses and patient participants who have generously and altruistically supported the APCB.

CCI
This work was awarded by Prostate Cancer Canada and is proudly funded by the Movember Foundation - Grant # D2013-36.
The CCI group would like to thank David Murray, Razmik Mirzayans, and April Scott for their contribution to this work.

CeRePP

COH
SLN is partially supported by the Morris and Horowitz Families Endowed Professorship

COSM
COSM is funded by The Swedish Research Council (grant for the Swedish Infrastructure for Medical Population-based Life-course Environmental Research – SIMPLER), the Swedish Cancer Foundation.

EPICAP
The EPICAP study was supported by grants from Ligue Nationale Contre le Cancer; Institut National du Cancer (INCa); Fondation ARC; Fondation de France; Agence Nationale de sécurité sanitaire de l'alimentation, de l'environnement et du travail (ANSES); Ligue départementale du Val de Marne.
The EPICAP study group would like to thank all urologists, Antoinette Anger and Hasina Randrianasolo (study monitors), Anne-Laure Astolfi, Coline Bernard, Oriane Noyer, Marie-Hélène De Campo, Sandrine Margaroline, Louise N'Diaye, Sabine Perrier-Bonnet (Clinical Research nurses)

ESTHER
The ESTHER study was supported by a grant from the Baden Württemberg Ministry of Science, Research and Arts.
The ESTHER group would like to thank Hartwig Ziegler, Sonja Wolf, Volker Hermann, Heiko Müller, Karina Dieffenbach, Katja Butterbach for valuable contributions to the study.

FHCRC
The FHCRC studies were supported by grants R01-CA080122, R01-CA056678, R01-CA082664, and R01-CA092579, and K05-CA175147 from the US National Cancer Institute, National Institutes of Health, with additional support from the Fred Hutchinson Cancer Research Center (P30-CA015704).
We thank all the individuals who participated in these studies.

Hamburg-Zagreb

IMPACT
The IMPACT study was funded by The Ronald and Rita McAulay Foundation, CR-UK Project grant (C5047/A1232), Cancer Australia, AICR Netherlands A10-0227, Cancer Australia and Cancer Council Tasmania, NIHR, EU Framework 6, Cancer Councils of Victorial and South Australia, Philanthropic donation to Northshore University Health System.
We acknowledge support from the National Institute for Health Research (NIHR) to the Biomedical Research Centre at The Institute of Cancer Research and Royal Marsden Foundation NHS Trust.
We acknowledge the IMPACT study steering committee, collaborating centres and participants.

KULEUVEN
F.C. and S.J. are holders of grants from FWO Vlaanderen (G.0684.12N and G.0830.13N), the Belgian federal government (National Cancer Plan KPC_29_023), and a Concerted Research Action of the KU Leuven (GOA/15/017). TVDB is holder of a doctoral fellowship of the FWO.

MCC-Spain

"The study was partially funded by the ""Accion Transversal del Cancer"", approved on the Spanish Ministry Council on the 11th October 2007, by the Instituto de Salud Carlos III-FEDER (PI08/1770, PI09/00773-Cantabria, PI11/01889-FEDER, PI12/00265, PI12/01270, PI12/00715, PI15/00069), by the Fundación Marqués de Valdecilla (API 10/09), by the Spanish Association Against Cancer (AECC) Scientific Foundation and by the Catalan Government DURSI grant 2009SGR1489. Samples: Biological samples were stored at the Parc de Salut MAR Biobank (MARBiobanc; Barcelona) which is supported by Instituto de Salud Carlos III FEDER (RD09/0076/00036). Also sample collection was supported by the Xarxa de Bancs de Tumors de Catalunya sponsored by Pla Director d'Oncologia de Catalunya (XBTC). ISGlobal acknowledges support from the Spanish Ministry of Science and Innovation through the "Centro de Excelencia Severo Ochoa 2019-2023" Program (CEX2018-000806-S), and support from the Generalitat de Catalunya through the CERCA Program."
We acknowledge the contribution from Esther Gracia-Lavedan in preparing the data. We thank all the subjects who participated in the study and all MCC-Spain collaborators.

MCCS
Melbourne Collaborative Cohort Study (MCCS) cohort recruitment was funded by VicHealth and Cancer Council Victoria. The MCCS was further augmented by Australian National Health and Medical Research Council grants 209057, 396414 and 1074383 and by infrastructure provided by Cancer Council Victoria. Cases and their vital status were ascertained through the Victorian Cancer Registry and the Australian Institute of Health and Welfare, including the National Death Index and the Australian Cancer Database.

MEC
The MEC was supported by NIH grants CA063464, CA054281, CA098758, and CA164973.

MOFFITT
The Moffitt group was supported by the US National Cancer Institute (R01CA128813, PI: J.Y. Park).

PLCO
This PLCO study was supported by the Intramural Research Program of the Division of Cancer Epidemiology and Genetics, National Cancer Institute, NIH
The authors thank Drs. Christine Berg and Philip Prorok, Division of Cancer Prevention at the National Cancer Institute, the screening center investigators and staff of the PLCO Cancer Screening Trial for their contributions to the PLCO Cancer Screening Trial. We thank Mr. Thomas Riley, Mr. Craig Williams, Mr. Matthew Moore, and Ms. Shannon Merkle at Information Management Services, Inc., for their management of the data and Ms. Barbara O'Brien and staff at Westat, Inc. for their contributions to the PLCO Cancer Screening Trial. We also thank the PLCO study participants for their contributions to making this study possible.

PRAGGA
PRAGGA was supported by Programa Grupos Emergentes, Cancer Genetics Unit, CHUVI Vigo Hospital, Instituto de Salud Carlos III, Spain.
PRAGGA wishes to thank Victor Muñoz Garzón, Manuel Enguix Castelo, Sara Miranda Ponte, Carmen M Redondo, Manuel Calaza, Francisco Gude Sampedro, Joaquín González-Carreró and the staff of the Department of Pathology and Biobank of University Hospital Complex of Vigo, Instituto de Investigación Sanitaria Galicia Sur (IISGS), SERGAS, Vigo, Spain; Máximo Fraga, José Antúnez and the Biobank of University Hospital Complex of Santiago, Santiago de Compostela, Spain; and Maria Torres, Angel Carracedo and the Galician Foundation of Genomic Medicine.

PROFILE
We would like to acknowledge the support of the Ronald and Rita McAulay Foundation and Cancer Research UK. We also acknowledge support from the National Institute for Health Research (NIHR) to

the Biomedical Research Centre at The Institute of Cancer Research and Royal Marsden Foundation NHS Trust
We acknowledge the Profile study steering committee and participants.

QLD
The QLD research is supported by The National Health and Medical Research Council (NHMRC) Australia Project Grants [390130, 1009458] and NHMRC Career Development Fellowship, Cancer Australia PdCCRS and Cancer Council Queensland funding to J Batra
The QLD team would like to acknowledge and sincerely thank the urologists, pathologists, data managers and patient participants who have generously and altruistically supported the QLD cohort.

SEARCH
SEARCH is funded by a programme grant from Cancer Research UK [C490/A10124] and supported by the UK National Institute for Health Research Biomedical Research Centre at the University of Cambridge. The University of Cambridge has received salary support in respect of PP from the NHS in the East of England through the Clinical Academic Reserve.

SFPCS
SFPCS was funded by California Cancer Research Fund grant 99-00527V-10182

SWOG-PCPT / SWOG-SELECT
SELECT and PCPT are funded by Public Health Service grants U10CA37429 and 5UM1CA182883 from the National Cancer Institute.
The authors thank the site investigators and staff and, most importantly, the participants from PCPT who donated their time to this trial.

Toronto
Prostate Cancer Canada Movember Discovery Grant (D2013-17) to RJH; Canadian Cancer Society Research Institute Career Development Award in Cancer Prevention (2013-702108) to RJH

UKGPCS
UKGPCS would also like to thank the following for funding support: The Institute of Cancer Research and The Everyman Campaign, The Prostate Cancer Research Foundation, Prostate Research Campaign UK (now Prostate Action), The Orchid Cancer Appeal, The National Cancer Research Network UK, The National Cancer Research Institute (NCRI) UK. We are grateful for support of NIHR funding to the NIHR Biomedical Research Centre at The Institute of Cancer Research and The Royal Marsden NHS Foundation Trust. UKGPCS should also like to acknowledge the NCRN nurses, data managers and Consultants for their work in the UKGPCS.
UKGPCS would like to thank all urologists and other persons involved in the planning, coordination, and data collection of the study. KM and AL were in part supported from the NIHR Manchester Biomedical Research Centre

ULM
The Ulm group received funds from the German Cancer Aid (Deutsche Krebshilfe).

WUGS
WUGS would like to thank the following for funding support: The Anthony DeNovi Fund, the Donald C. McGraw Foundation, and the St. Louis Men's Group Against Cancer.

Information about the PRACTICAL consortium can be found at http://practical.icr.ac.uk/. The PRACTICAL consortium investigators are: Rosalind A. Eeles, Zsofia Kote-Jarai, Kenneth R. Muir, UKGPCS collaborators, Artitaya Lophatananon, Catherine M. Tangen, Phyllis J. Goodman, Ian M. Thompson Jr., Alicja Wolk, Niclas Håkansson, Demetrius Albanes, Stephanie Weinstein, Nora Pashayan, Alison M.

Dunning, Maya Ghoussaini, Jyotsna Batra, APCB BioResource (Australian Prostate Cancer BioResource), Suzanne Chambers, Judith A. Clements, Lisa Horvath, Leire Moya, Gail P. Risbridger, Wayne Tilley, Sonja I. Berndt, Stephen Chanock, Gerald L. Andriole, Robert N. Hoover, Mitchell J. Machiela, Stella Koutros, Laura E. Beane Freeman, Florence Menegaux, Thérèse Truong, Christopher A. Haiman, Fredrick R. Schumacher, Loic Le Marchand, Xin Sheng, Robert J. MacInnis, Melissa C. Southey, Graham G. Giles, Roger L. Milne, Robert J. Hamilton, Neil E. Fleshner, Antonio Finelli, Manolis Kogevinas, Javier Llorca, Gemma Castaño-Vinyals, Antonio Alcaraz, Olivier Cussenot, Géraldine Cancel-Tassin, The IMPACT Study Steering Committee and Collaborators, Janet L. Stanford, Elaine A. Ostrander, Adam S. Kibel, Bettina F. Drake, Hermann Brenner, Xin Gao, Bernd Holleczek, Ben Schöttker, Esther M. John, Jong Y. Park, Hui-Yi Lin, Nawaid Usmani, Matthew Parliament, Aswin Abraham, Susan L. Neuhausen, Yuan Chun Ding, Linda Steele, Jose Esteban Castelao, Manuela Gago-Dominguez, Maria Elena Martinez, Frank Claessens, Steven Joniau, Thomas Van den Broeck, Christiane Maier, Manuel Luedeke, Thomas Schnoeller, Marija Gamulin, Davor Lessel, Tomislav Kulis, The Profile Study Steering Committee.

**UKHLS (Understanding Society: The UK Household Longitudinal Study)**
**"Understanding Society Scientific Group"**

Michaela Benzeval[1], Jonathan Burton[1], Nicholas Buck[1], Annette Jäckle[1], Meena Kumari[1], Heather Laurie[1], Peter Lynn[1], Stephen Pudney[1], Birgitta Rabe[1], Dieter Wolke[2]
[1]Institute for Social and Economic Research
[2]University of Warwick

### SUPPLEMENTARY METHODS

A summary of the methods, together with a full description of genome-wide association analyses and follow-up analyses is described below. Written informed consent was obtained from every participant in each study, and the study was approved by relevant ethics committees.

#### Quality control checks of individual studies

All study files were checked for quality using the software EasyQC[1] that was adapted to the format from RVTESTS[2]. The checks performed included allele frequency differences with ancestry specific reference panels, total number of markers, total number of markers not present in the reference panels, imputation quality, genomic inflation factor and trait transformation. We excluded 2 studies that did not pass our quality checks in the data.

#### Genome-wide association study meta-analysis

We first performed ancestry groups specific GWAS meta-analyses of 173 studies of EUR, 56 studies of EAS, 29 studies of AFR, 11 studies of HIS and 12 studies of SAS. Meta-analyses within ancestry groups were performed as described before[3,4] using a modified version of RAREMETAL[5] (version 4.15.1), which accounts for multi-allelic variants in the data. Study-specific GWAS are described in Suppl. Tables 1 - 3. We kept in our analyses SNPs with an imputation accuracy ($r^2_{\mathrm{INFO}}$) >0.3, Hardy-Weinberg Equilibrium (HWE) p-value ($P_{\mathrm{HWE}}$) >$10^{-8}$ and a minor allele count (MAC) >5 in each study. Next, we performed a fixed-effect inverse variance weighted meta-analysis of summary statistics from all five ancestry groups GWAS meta-analysis using a custom R script.

#### Hold-out sample from the UK Biobank

We excluded 56,477 UK Biobank (UKB) participants from our discovery GWAS for following analyses including quantification of population stratification. More precisely, our hold-out EUR sample consists of 17,942 siblings pairs and 981 trios (i.e. two parents and one offspring) plus all UKB participants with an estimated genetic relationship, with our set of siblings pairs and trios, larger than 0.05. We identified 14,587 individuals among these 56,477 UKB participants who were unrelated (i.e. HM3 SNP-based estimated related <0.05) with each other and used their data to quantify the variance explained by SNPs within GWS loci (described below) and prediction accuracy of PGS.

#### Conditional and joint association analyses

We performed conditional and joint-analyses (COJO) of each of the five ancestry groups GWAS meta-analyses using the software GCTA.[6,7] The GCTA-COJO method implements a stepwise model selection that aims at retaining a set of SNPs, which joint effects reach genome-wide significance, defined in this study as a p-value $P<5\times10^{-8}$. In addition to GWAS summary statistics, COJO analyses also require genotypes from an ancestry-matched sample that is used a LD reference. For all sets of genotypes used as LD reference panels, we selected HM3 SNPs with $r^2_{\mathrm{INFO}}$>0.3 and $P_{\mathrm{HWE}}$ >$10^{-6}$. For EUR, we used genotypes at 1,318,293 HM3 SNPs (MAC>5) from 348,501 unrelated EUR participants of the UKB as our LD reference. For EAS, we used genotypes at 1,034,263 quality-controlled (MAF>1%, SNP missingness <5%) HM3 SNPs from a merged panel of N=5,875 unrelated participants from UKB (N=2,257) and Genetic Epidemiology Research on Aging (GERA; N=3,618). Data from the GERA study were obtained from the database of Genotypes and Phenotypes (dbGaP; accession number: phs000788.v2.p3.c1) under project 15096. For SAS, we used genotypes at 1,222,935 HM3 SNPs (MAC>5; SNP missingness <5%) from 9,448 unrelated individuals. For AFR, we used genotypes at 1,007,949 quality-controlled (MAF>1%, SNP missingness <5%) HM3 SNPs from a merged panel of 15,847 participants from the Women's Health Initiative (WHI; N=7,480), and the National Heart, Lung, and Blood Institute's Candidate Gene Association Resource (CARe[8]; N=8,367). Both WHI and CARe datasets were obtained from dbGaP (accession numbers: phs000386 for WHI; CARe including phs000557.v4.p1, phs000286.v5.p1, phs000613.v1.p2, phs000284.v2.p1, phs000283.v7.p3 for ARIC, JHS, CARDIA, CFS and MESA cohorts) and processed following the protocol provided by the dbGaP data submitters. After excluding samples with >10% missing values and retained only unrelated individuals, our final LD reference included data from N=10,636 unrelated AFR individuals. For HIS, we used genotypes at 1,246,763 sequenced HM3 SNPs (MAF>1%) from N=4,883 unrelated samples from the Hispanic Community Health Study / Study of Latinos (HCHS/SOL; dbGaP accession number: phs001395.v2.p1) cohorts. Finally, we performed a COJO analysis of the combined meta-analysis of all ancestries (referred to as $META_{FE}$ in the main text) using 348,501 unrelated EUR participants of the UKB.

To assess if SNPs detected in non-EUR were independent of signals detected in EUR, we performed another COJO analysis of ancestry groups GWAS by fitting jointly SNPs detected in EUR with those detected in each of the non-EUR GWAS meta-analyses. For each non-EUR GWAS, we performed a single-step COJO analysis only including SNPs identified in that non-EUR GWAS and which LD squared correlation ($r_{\text{LD}}^2$) with any of the EUR signals (marginally or conditionally GWS) is lower than 0.8 in both EUR and corresponding non-EUR data. Single-step COJO analyses were performed using the *--cojo-joint* option of GCTA, which does not involve model selection and simply approximates a multivariate regression model where all selected SNPs on a chromosome are fitted jointly. LD correlations used in these filters were estimated in ancestry-matched samples of the 1,000 Genomes Project (1KGP; release 3). More specifically, LD was estimated in 661 AFR, 347 HIS (referred to with the AMR label in the 1000 Genomes Project), 504 EAS, 503 EUR and 489 SAS 1KGP participants. We used the same LD reference samples in these analyses as for our main discovery analysis described at the beginning of the section.

**$F_{ST}$ calculation and (stratified) LD score regression**

We used two statistics to evaluate whether an EUR LD reference could approximate well enough the LD structure in our trans-ancestry GWAS meta-analysis. The first statistic that we used is the Wright fixation index,[9] which measures allele frequency divergence between two populations. We used the Hudson's estimator of $F_{ST}$[10] as previously recommended[11] to compare allele frequencies from our $META_{FE}$ with that from our EUR GWAS meta-analysis and an independent replication sample from the Estonian Biobank (EBB). The other statistics that we used is the attenuation ratio statistic from the LD score regression methodology. These LD score regression analyses were performed using the version 1.0 of the LDSC software and using LD scores calculated from EUR participants of the 1KGP (URLs). Moreover, we performed a stratified LD score regression analysis to quantify the enrichment of height heritability in 97 genomic annotations curated and described in Gazal et al.[12] as the baseline-LD model. Annotation-weighted LD scores used for those analyses were also calculated using data from 1KGP (URLs).

**Density of GWS signal and enrichment near OMIM genes**

We defined the density of independent signals around each GWS SNP as the number of other independent associations identified with COJO within a 100 kb window on both sides. Therefore, a SNP with no other associations within 100 kb has a density of 0, while a SNP colocalising with 20 other GWS associations within 100 kb will have a density of 20. We quantified the standard error of the mean signal density across the genome using a leave-one-chromosome-out jackknife procedure. We then quantified the enrichment of 462 curated OMIM[13] genes near GWS SNPs with large signal density, by counting the number of OMIM genes within 100 kb of a GWS SNP, then comparing that number for SNPs with a density of 0 and those with a density of at least 1. The strength of the enrichment was measured using an odds ratio calculated from a 2×2 contingency table: "presence/absence of an OMIM gene" versus "density of 0 or larger than 0". To assess the significance of the enrichment, we simulated the distribution of enrichment statistics for a random set of 462 length-matched genes. We used 22 length classes (<10kb; between $i \times 10$ kb and $(i+1) \times 10$ kb, with i=1,…,9; between $i \times 100$ kb and $(i+1) \times 100$ kb, with i=1,…,10; and between 1 Mb and 1.5 Mb; between 1.5 Mb and 2 Mb; and >2Mb) to match OMIM genes with random genes. OMIM genes within a given length class were matched with the same number of non-OMIM genes present in the class. We sampled 1,000 random sets of genes and calculated for each them an enrichment statistic. Enrichment p-value was calculated as number of times enrichment statistics of random genes exceeded that of OMIM genes. The list of OMIM genes is provided in Suppl. Table 11.

### Replication analyses

To assess the replicability of our results, we tested if the correlation $\rho_b$ of estimated SNP effects between our discovery GWAS and our replication sample of 49,160 participants of the Estonian Biobank (EBB) was statistically different from 1. We used the estimator of $\rho_b$ from Qi et al.[14], which accounts for sampling errors in both discovery and replication samples. Standard errors were calculated using a leave-one-SNP-out jackknife procedure. We quantified the correlation of marginal and also that of joint SNP effects. Joint SNP effects in our replication sample were obtained by performing a single-step COJO analysis of GWAS summary statistics from our EBB sample, using the same LD reference as in the discovery GWAS. Correlation of SNP effects were calculated after correcting SNP effects for winner's curse using the method from Zhong and Prentice.[15] We provide the R scripts used to apply these corrections and estimate correlation of SNP effects (URLs). The expected proportion, $E[P]$, of sign-consistent SNP effects between discovery and replication was calculated using the quadrant probability of a standard bivariate Gaussian distribution with correlation $E[\rho_b]$, denoting the expected correlation between estimated SNP effects in the discovery and replication sample:

$$(1) \qquad \mathrm{E[P]} = \frac{1}{2} + \frac{\sin^{-1}(\mathrm{E}[\rho_b])}{\pi},$$

where $\sin^{-1}$ denotes the inverse of the sine function and $\mathrm{E}[\rho_b]$ the expectation of the $\rho_b$ statistic under the assumption that the true SNP effects are the same across discovery and replications cohorts.

E[P] was calculated as

$$(2) \quad \mathrm{E}[\rho_b] = \frac{\sigma_b^2}{\sqrt{\left(\sigma_b^2 + [1-\sigma_b^2 h_d]/(N_d h_d)\right)\left(\sigma_b^2 + [1-\sigma_b^2 h_r]/(N_r h_r)\right)}}$$

where $N_d$ and $N_r$ denote the sizes of the discovery and replication samples, respectively; $h_d$ and $h_r$ the average heterozygosity (i.e. $2 \times \mathrm{MAF} \times (1 - \mathrm{MAF})$) across GWS SNPs in the discovery and replication samples, respectively; and $\sigma_b^2$ the mean per-SNP variance explained by GWS SNPs, which we calculated (as in Qi et al.[14]) as the sample variance of estimated SNP effects in the discovery sample minus the median squared standard error.

### Variance explained by GWS SNPs and loci

We estimated the variance explained by GWS SNPs using the Genetic relationship based REstricted Maximum Likelihood (GREML) approach implemented in GCTA.[6,16] This approach involves two main steps: (i) calculation of genetic relationships matrices (GRM) and (ii) estimation of variance components corresponding to each of these matrices using a REML algorithm. We partitioned the genome in two sets containing GWS loci on the one hand and all other HM3 SNPs on the other hand. GWS loci were defined as non-overlapping genomic segments containing at least 1 GWS SNP and such that GWS SNPs in adjacent loci are >2×35kb away of each other (i.e. 35kb window on each side). We then calculated a GRM based on each set of SNPs and estimated jointly a variance explained by GWS alone and that explained by the rest of the genome. We performed these analyses in multiple samples independent of our discovery GWAS, which include participants of diverse ancestry. Details about the samples used for these analyses are provided below. We extended our analyses to also quantify the variance explained by GWS loci using alternative definitions based on a window size of 0 kb and 10 kb around GWS SNPs (Suppl. Figs. 16 - 17).

We also repeated our analyses using a random set of 12,111 SNPs matched with GWS SNPs on MAF and LD. Loci for these 12,111 random SNPs were defined similarly as for GWS loci. To match random SNPs with GWS SNPs on MAF and LD, we first created 28 MAF-LD classes of HM3 SNPs (7 MAF classes × LD score classes). MAF classes were defined as <1%; between 1% and 5%; between 5% and 10%; between

10% and 20%; between 20% and 30%; between 30% and 40%; and between 40% and 50%. LD score classes were defined using quartiles of the HM3 LD score distribution. We next matched GWS SNPs in each of the 28 MAF-LD classes, with the same number of SNPs randomly sampled from that MAF-LD class.

**Prediction analyses**

We quantified the accuracy of various predictors of adult height as the squared Pearson correlation coefficient between a given predictor and height of study participants. For these analyses, height was corrected for mean and variance sex differences and 20 genotypic principal components calculated form subsets of LD pruned HM3 SNPs ($r_{\mathrm{LD}}^2$<0.1). We used height of siblings or parents as predictor of height as well as polygenic scores (PGS) calculated as a weighted sum of height-increasing alleles. The direction and magnitude of these weights was determined by estimated SNP effects from our discovery GWAS meta-analyses. No calibration of tuning parameters in a validation was performed. The prediction accuracy of sibling's height was assessed in 17,942 unrelated sibling pairs from the UKB. Those pairs were determined by intersecting the list of UKB siblings pairs previously determined by Bycroft et al.[17] with a list of genetically determined European ancestry participants from the UKB also described previously.[18] We then filtered the resulting list for SNP-based genetic relationship between members of different families to be smaller than 0.05. The prediction accuracy of parental height (each parent and their average) was assessed in 981 unrelated trios obtained as described above by crossing information from Bycroft et al.[17] with that from Yengo et al.[18]. We quantified the within-family variance explained by PGS as the squared correlation of height difference between siblings with PGS difference between siblings. We describe in **Supplementary Note 2** how familial information and PGS were combined to generate a single predictor.

**Samples used for prediction analyses and estimation of variance explained by SNPs in GWS loci**

We quantified the accuracy of a PGS based on GWS SNPs as well as the variance explained by SNPs within GWS loci, in 8 different datasets independent of our discovery GWAS meta-analyses. These datasets include 2 samples of EUR from UKB (N=14,587) and the Lifelines study (N=14,058), 2 samples of AFR from UKB (N=6,911) and the PAGE study (N=8,238), 2 samples of EAS (N=2,246) from UKB and the China Kadoorie Biobank (CKB; N=47,693), 1 sample of SAS from UKB (N=9,257) and 1 sample of HIS from the PAGE study (N=4,939). Analyses were adjusted for age, sex, 20 genotypic principal components and study specific covariates (e.g., recruitment centres). Genotypes of EUR UKB participants were imputed to the Haplotype Reference Consortium (HRC) and to a combined reference panel including haplotypes from the 1KG Project and the UK10K Project. To improve variants coverage in non-EUR participants of UKB, we re-imputed their genotypes to the 1KG reference panel, as described previously.[19] Lifelines samples were imputed to the HRC panel. PAGE and CKB were imputed to the 1KG reference panel. Standard quality control ($r_{\mathrm{INFO}}^2$>0.3, $P_{\mathrm{HWE}}$ >$10^{-6}$ and MAC >5) were applied to imputed genotypes in each dataset.

**Contribution of LD and MAF to the loss of prediction accuracy in African ancestry participants**

We defined the EUR-to-AFR relative accuracy as the ratio of prediction accuracies from an AFR sample over that from a EUR sample. We used the method of Wang et al.[19] to quantify the expectation of that relative accuracy under the assumption that causal variants and their effects are shared between EUR and AFR, while MAF and LD structures can differ. In brief, this methods contrasts LD and MAF patterns within 100 kb windows around each GWS SNPs and use them to predict the expected loss of accuracy. As in Wang et al.[19], we used genotypes from 503 EUR and 661 AFR participants of the 1KGP as a reference sample to estimate ancestry-specific MAF and LD correlations between GWS SNPs and SNPs in their close vicinity and defined candidate causal variants as any sequenced SNP with an $r_{\mathrm{LD}}^2$>0.45 with a GWS SNP within that 100 kb window. Standard errors were calculated using a delta-method approximation as described in Wang et al.[19]

**Down-sampled GWAS analyses**

In addition to our EUR GWAS meta-analysis and our trans-ancestry meta-analysis ($META_{FE}$), we re-analysed 5 down-sampled GWAS as shown in Table 2. These down-sampled GWAS include various iterations of previous efforts of the GIANT consortium and have a sample size varying between ~130,000 and 2.5M (EUR participants from 23andMe). To ensure sufficient genomic coverage of HM3 SNPs we imputed GWAS summary statistics from Lango-Allen et al.[3], Wood et al.[4] and Yengo et al.[18] with SSImp[20] using haplotypes from 1KGP as a LD reference. GWAS summary statistics from Kang-Allen et al. only contain p-values (P), height-increasing alleles and per-SNP sample sizes (N). We therefore derived SNP effects and their corresponding standard errors using linear regression theory (R script; URLs). Imputed GWAS summary statistics from these 3 studies are made publicly available on the GIANT consortium website (URLs). We next performed a COJO analysis of all down-sampled GWAS using genotypes of 348,501 unrelated EUR participants of the UKB as a LD reference panel, as for our $META_{FE}$ and EUR GWAS meta-analysis.

**Gene prioritisation using Summary data-based Mendelian Randomization (SMR)**

We performed a summary-data-based Mendelian randomization (SMR) to identify genes which expressions could mediate SNP effects on height. We used publicly available gene expression quantitative trait loci (eQTL) identified from two large eQTL studies, namely the GTEx[21] version 8 and the eQTLgen studies (URLS). To ensure that our SMR results robustly reflect causality or pleiotropic effects of height-associated SNPs on gene expression, we only report here significant SMR results (i.e. $P<5\times10^{-8}$), which do not pass the HEterogeneity In Dependent Instrument (HEIDI) test (i.e. $P>0.01$; METHODS). The significant threshold for the HEIDI test was chosen based on recommendations from Wu et al.[22]

**Selection of OMIM genes**

To generate a list of genes known to underlie syndromes of abnormal skeletal growth, we queried the Online Mendelian Inheritance in Man database (OMIM, www.omim.org). From July, 2019 to August, 2020, we performed queries using search terms of "short stature," "tall stature," "overgrowth," "skeletal dysplasia," and "brachydactyly." We then used the free text descriptions in OMIM to manually curate the resulting combined list of genes, as well as genes in our earlier list from Wood et al.[4] and all genes listed as causing skeletal disease in an online endocrine textbook (endotext.org, accessed in September, 2020). For short stature, we only included genes underlying syndromes where short stature was either consistent (<-2 SD in the great majority of patients with data recorded), or present in multiple families/sibships and accompanied by either (a) more severe short stature (-3 SD), or (b) skeletal dysplasia was present (beyond poor bone quality/fractures) or (c) brachydactyly/shortened digits/disproportionate short stature/limb shortening was present (not simply absence of specific bones). We removed genes underlying syndromes where short stature was likely attributable to failure to thrive, specific metabolic disturbances, intestinal failure/enteropathy and/or very severe disease (e.g. early lethality or severe neurologic disease). For tall stature/overgrowth, we only included genes underlying syndromes where tall stature was consistent (> +2 SD in the great majority of patients with data recorded) or present in multiple families/sibships and accompanied by either (a) more severe tall stature (> +3 SD) or (b) arachnodactyly. For brachydactyly, we required more than only fifth finger involvement, and that brachydactyly be either consistent (present in the great majority of patients) or accompanied by consistent short stature or other skeletal dysplasias. For skeletal dysplasias, we only considered genes underlying syndromes where the skeletal dysplasia involved long bones or the spine and was accompanied by short stature, brachydactly, or limb/digit shortening. We also included all genes in a list we generated in Lango-Allen et al.[3], which was curated using similar criteria. The resulting list contained 536 genes, of which 462 (Suppl. Table 11) are autosomal based on annotation from PLINK (URL: https://www.cog-genomics.org/static/bin/plink/glist-hg19).

### SUPPLEMENTARY FIGURES

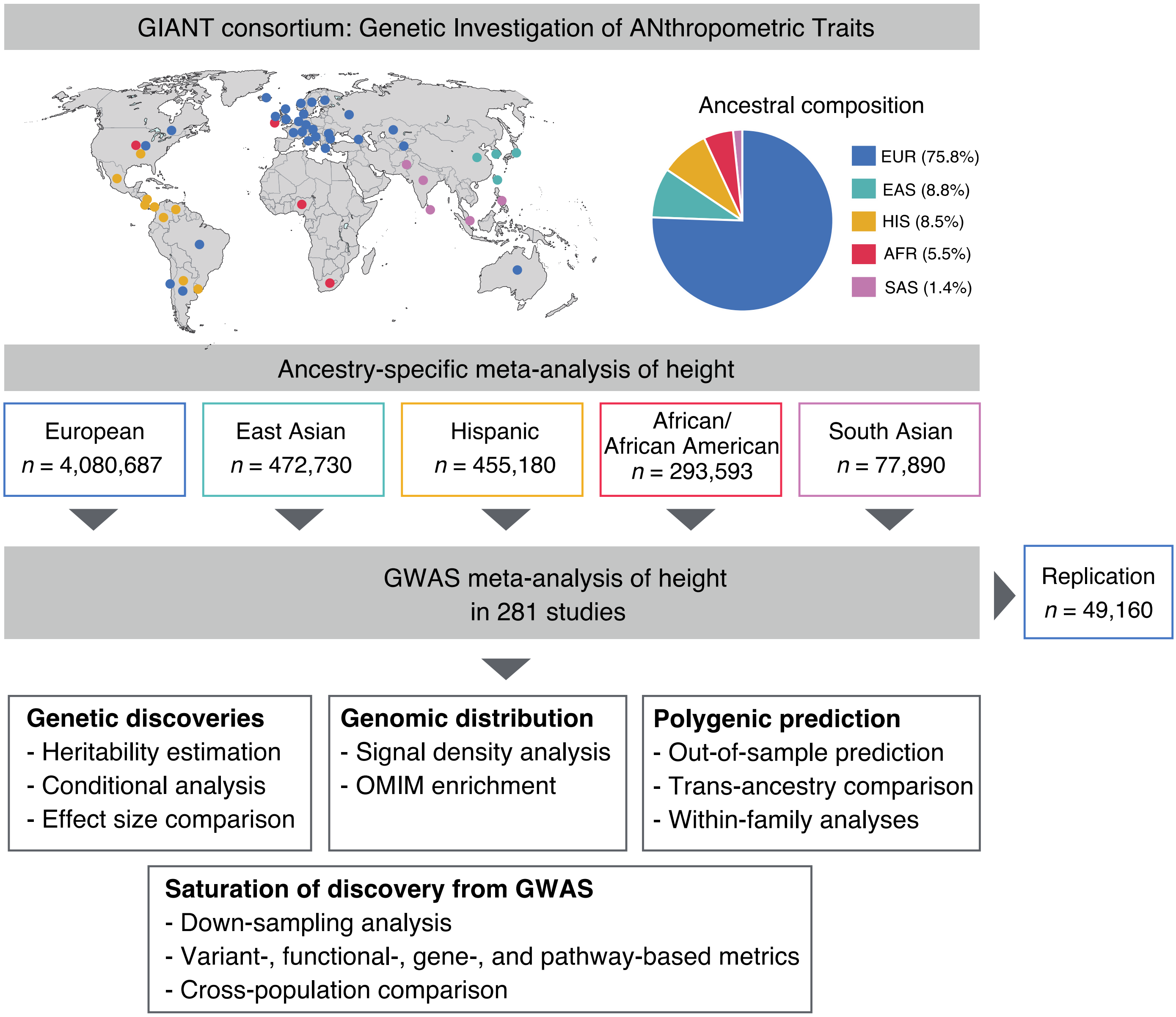


**Suppl. Fig. 1.** Overview of study design and analytical strategy.

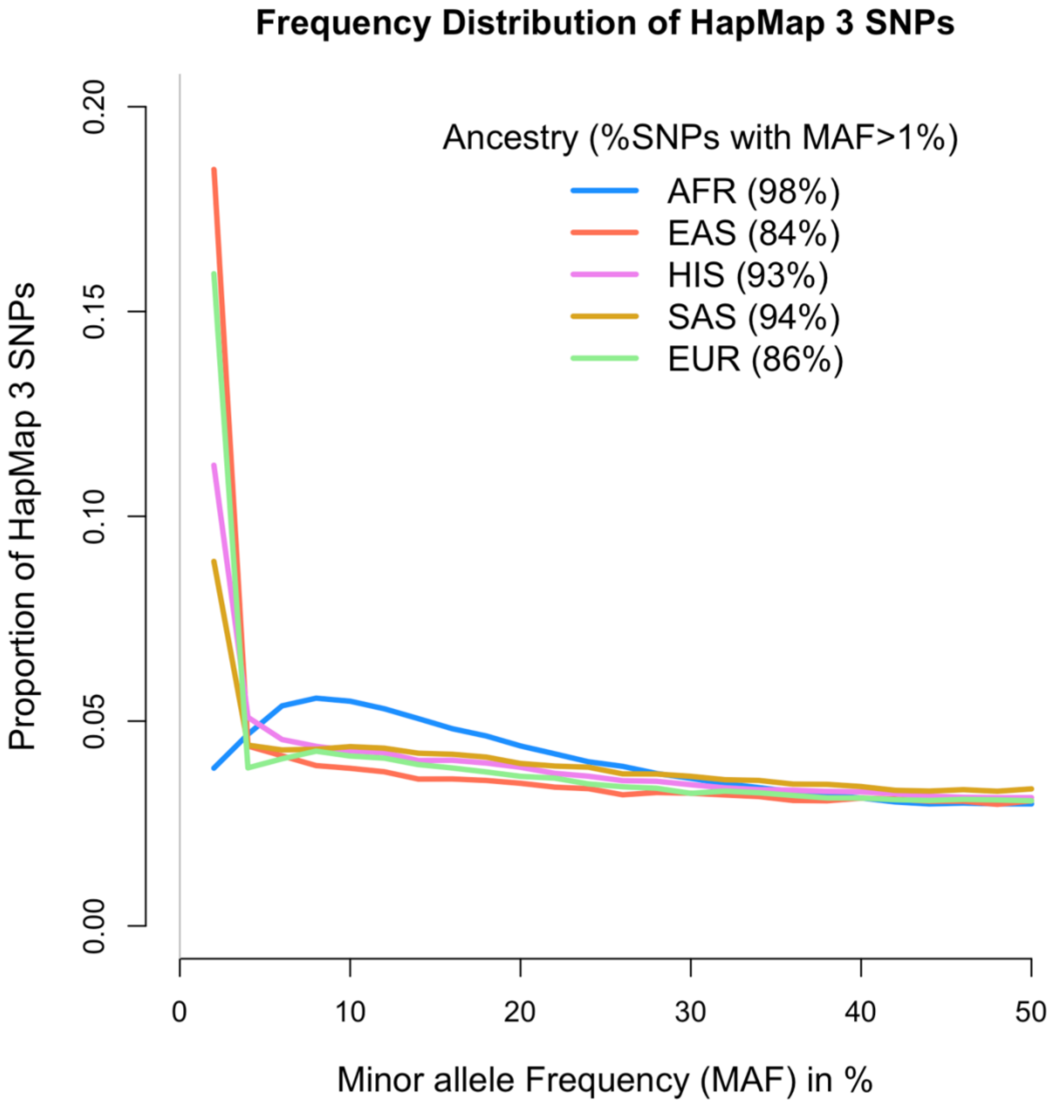


**Suppl. Fig. 2.** Minor allele frequency (MAF) distribution of HapMap 3 SNPs across 5 ancestries: European (EUR), Hispanic (HIS), African (AFR), East-Asian (EAS) and South-Asian (SAS).

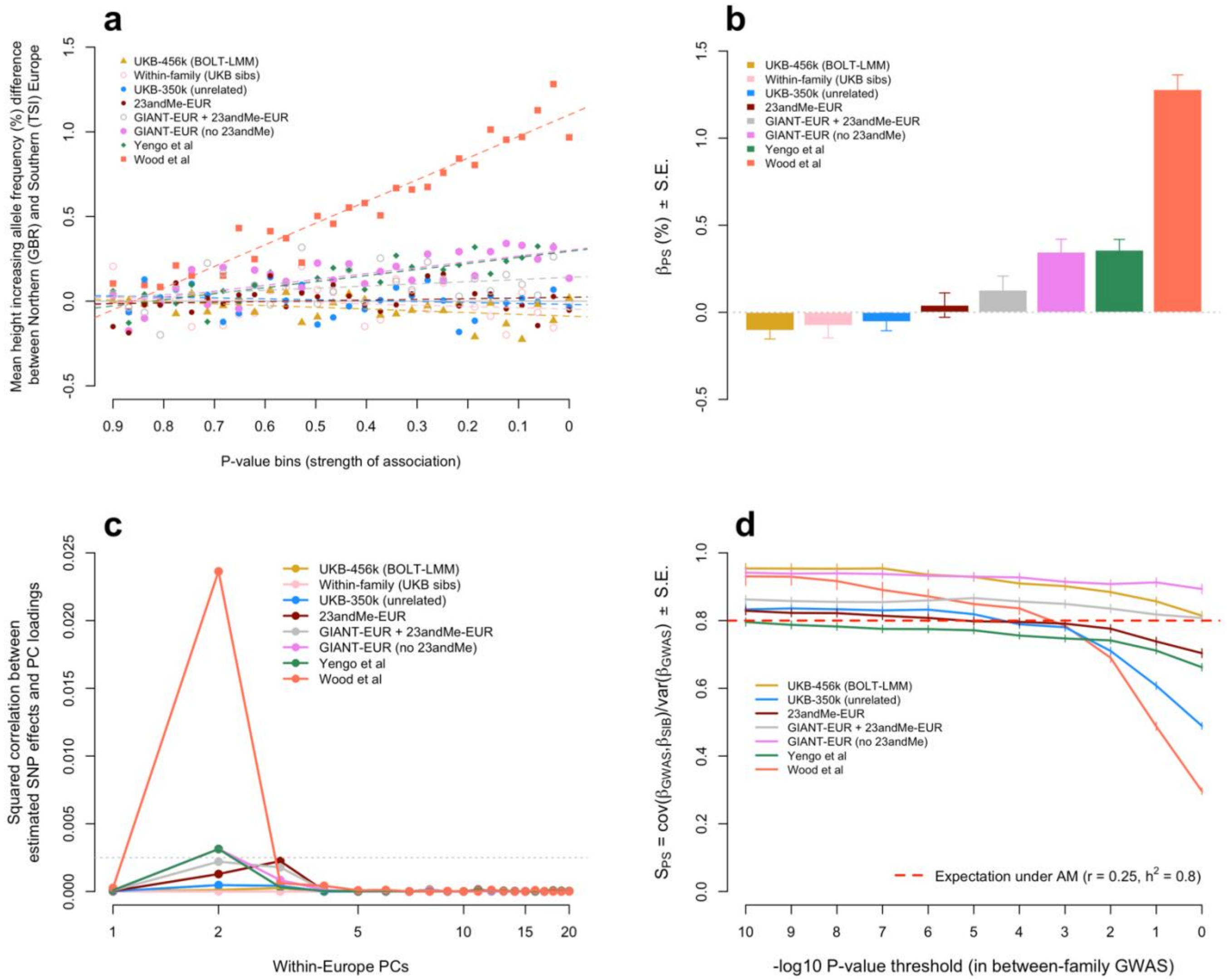


**Suppl. Fig. 3. Quantification of confounding due to population stratification (PS) in various European ancestry (EUR) GWAS of height.** UKB-456k (BOLT-LMM): GWAS in 456,414 EUR participants of the UKB. UKB-350k (unrelated): GWAS in 348,501 unrelated (i.e. estimated genetic relatedness <0.05) EUR participants of UKB. 23andMe-EUR: GWAS in EUR participants of 23andMe. GIANT-EUR: GWAS meta-analysis of 173 EUR cohorts from the GIAN consortium. Within-family (UKB): Family-based GWAS performed in 17,942 independent EUR sibling pairs from the UKB. **Panel a** represents the relationship between strength of association between 101,360 independent (LD $r^2<0.1$) HM3 SNPs and height (x-axis: 30 SNP bins) and height-increasing allele frequency differences between the British (GBR) and Tuscan (TSI) samples from the 1,000 Genomes Project (1KGP). A significant positive slope ($\beta_{PS}$) indicates uncorrected population stratification. **Panel b** shows estimates of $\beta_{PS}$ for different GWAS and their associated leave-one-SNP-bin-out jackknife standard errors (S.E.). **Panel c** shows the variance in estimated SNP effects from various height GWAS that is explained by SNP loading on genotypic principal components (PC) calculated among 503 EUR samples from the 1KGP. The x-axis in **Panel c** indicates number of vectors of PC loadings including in the regression model (SNP effect regressed on PC loading). **Panel d** shows on the y-axis the slope ($S_{PS} = cov(\beta_{GWAS}, \beta_{SIB})/var(\beta_{GWAS})$) from regressing SNP effects estimated in a family-based GWAS ($\beta_{SIB}$) onto SNP effects estimated in a standard population-based GWAS ($\beta_{GWAS}$). The x-axis in **Panel d** is the -log10 of the p-value threshold used to ascertain SNPs for estimating the regression slope. For each p-value threshold, the effects of ascertained SNPs were corrected for winner's curse (**Suppl. Methods**). The horizontal red dotted line represented the expected slope under assortative mating (AM) assuming an equilibrium heritability $h^2 = 0.8$ and a spousal correlation $r = 0.25$ (**Suppl. Note 1**).

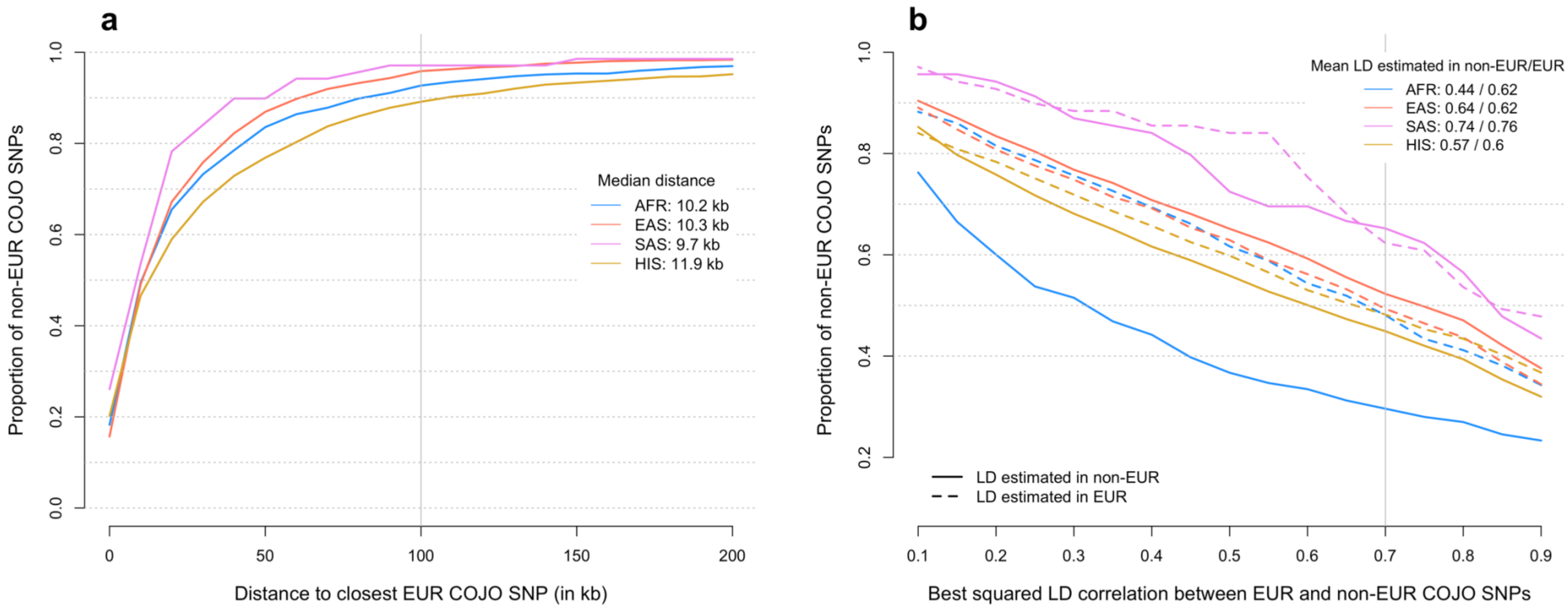


Suppl. Fig. 4. **Colocalization of height-associated signals across ancestries. Panel a:** Proportion (y-axis) of genome-wide significant (GWS) SNPs identified in our GWAS meta-analyses of non-European (non-EUR: African – AFR; East-Asian – EAS; South-Asian – SAS; Hispanic – HIS) ancestry/ethnicity participants thar are located within a certain distance (x-axis) of GWS SNPs identified in our GWAS meta-analysis of EUR participants only. **Panel b**: Proportion (y-axis) of GWS SNPs from non-EUR GWAS meta-analyses being in linkage disequilibrium (LD; x-axis) with a SNP reaching GWS in our EUR GWAS meta-analysis. LD was calculated either in a EUR (dotted line) or in a corresponding non-EUR sample of the 1,000 Genomes reference.

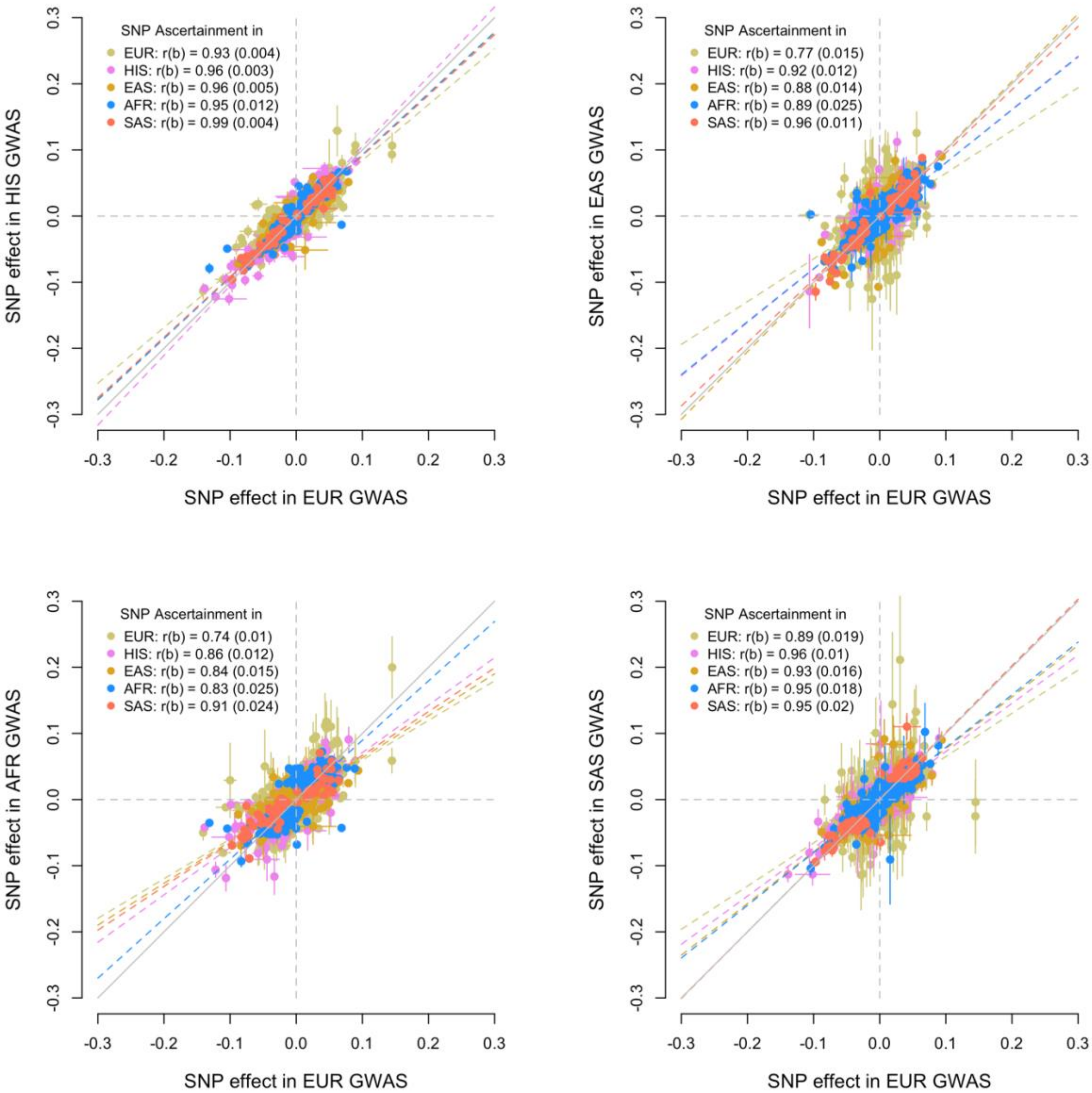


Suppl. Fig. 5. **Correlation of marginal SNP effects between European ancestry (EUR, on the x-axis) and non-EUR GWAS** (i.e. Hispanic (HIS), African (AFR), East-Asian (EAS) and South-Asian (SAS) on the y-axis). Correlations *r(b)* were for corrected for estimation errors as described in the **Suppl. Methods** section. Each estimate of the correlation of SNP effect is based on SNPs reaching marginal genome-wide significance in any of the 5 ancestries analysed. Standard errors for *r(b)* were obtained using jackknife. Error bars denote standard errors of SNP effects.

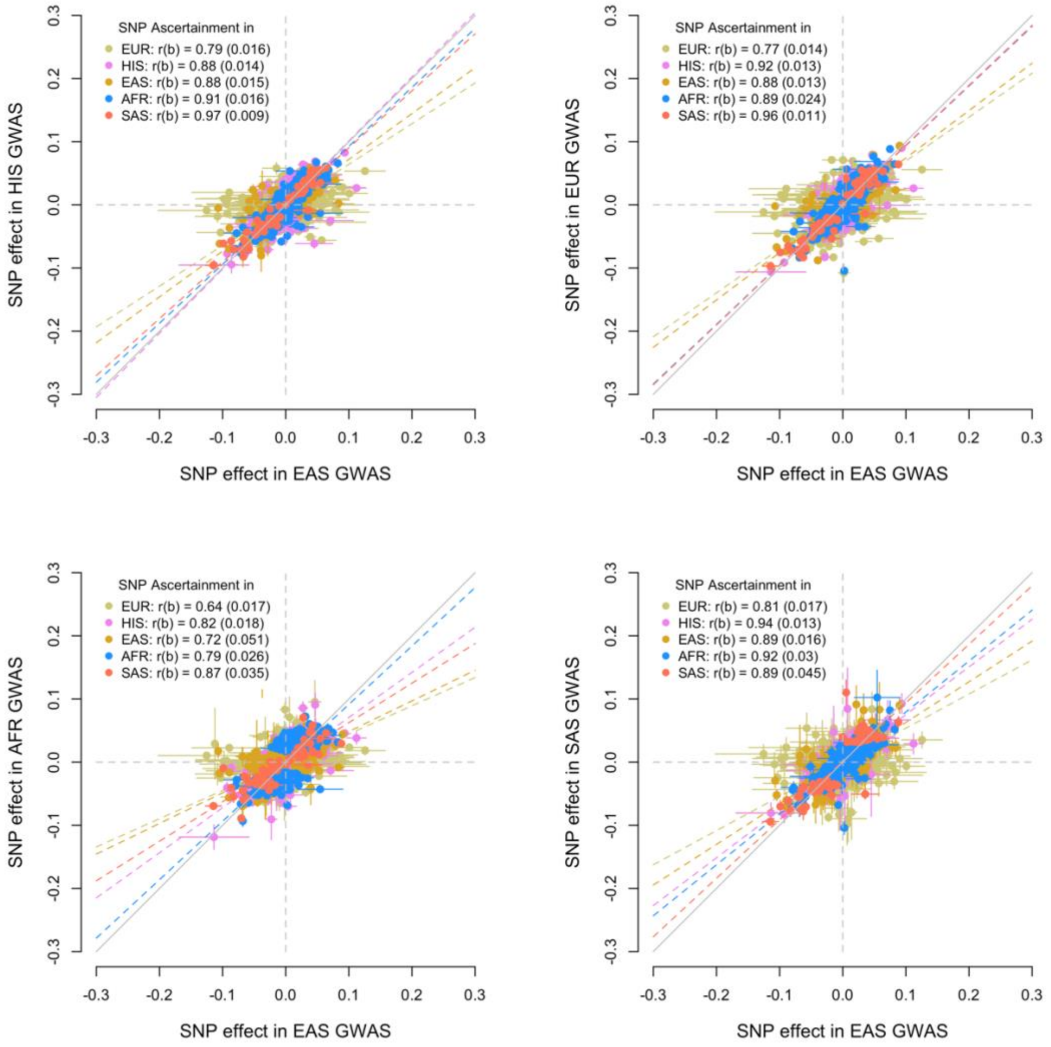


Suppl. Fig. 6. **Correlation of marginal SNP effects between East-Asian ancestry (EAS, on the x-axis) and non-EAS GWAS** (i.e. Hispanic (HIS), African (AFR), European (EUR) and South-Asian (SAS) on the y-axis). Correlations *r(b)* were for corrected for estimation errors as described in the **Suppl. Methods** section. Each estimate of the correlation of SNP effect is based on SNPs reaching marginal genome-wide significance in any of the 5 ancestries analysed. Standard errors for *r(b)* were obtained using jackknife. Error bars denote standard errors of SNP effects.

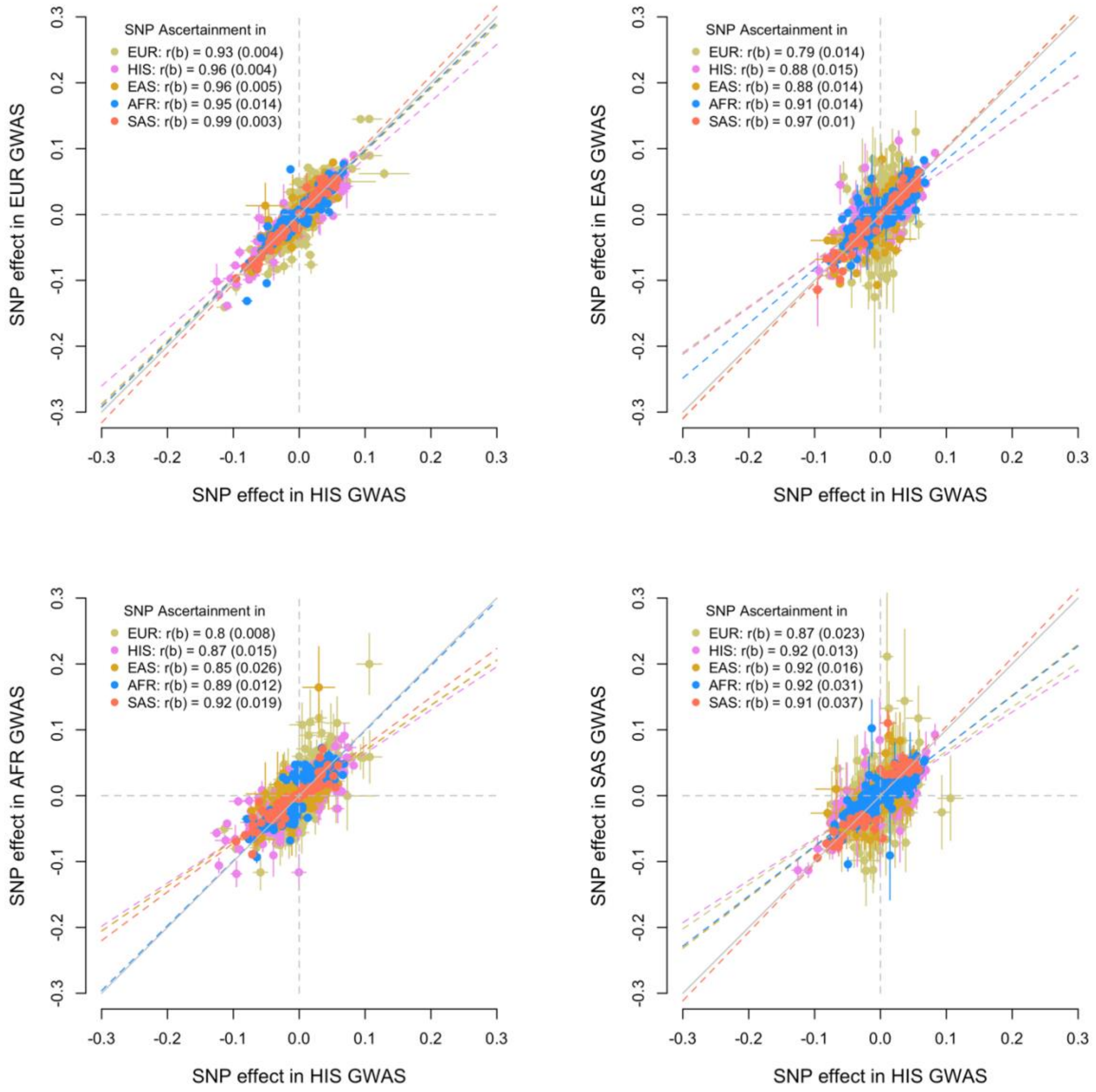


**Suppl. Fig. 7. Correlation of marginal SNP effects between Hispanic ethnicity (HIS, on the x-axis) and non-HIS GWAS** (i.e. European (EUR), African (AFR), East-Asian (EAS) and South-Asian (SAS) on the y-axis). Correlations *r(b)* were for corrected for estimation errors as described in the **Suppl. Methods** section. Each estimate of the correlation of SNP effect is based on SNPs reaching marginal genome-wide significance in any of the 5 ancestries analysed. Standard errors for *r(b)* were obtained using jackknife. Error bars denote standard errors of SNP effects.

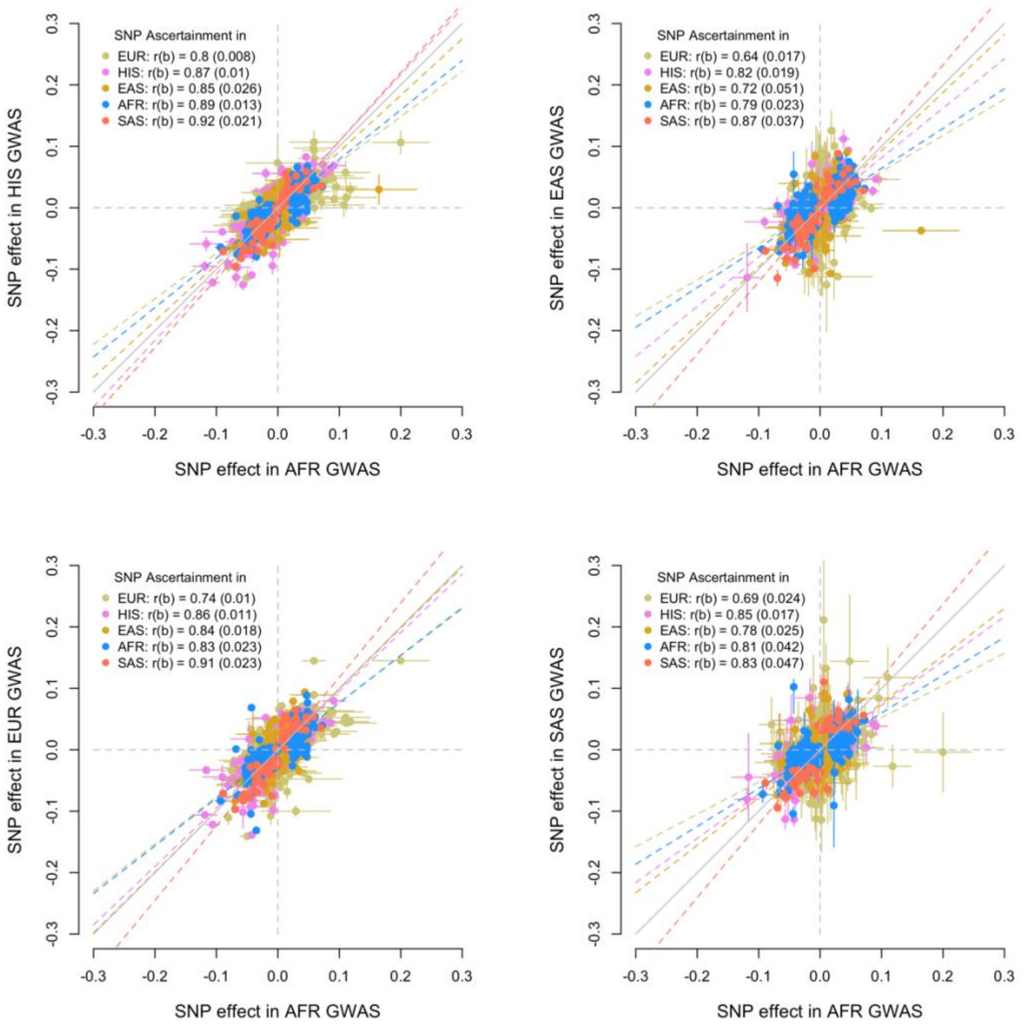


Suppl. Fig. 8. **Correlation of marginal SNP effects between African ancestry (AFR, on the x-axis) and non-AFR GWAS** (i.e. Hispanic (HIS), European (EUR), East-Asian (EAS) and South-Asian (SAS) on the y-axis). Correlations *r(b)* were for corrected for estimation errors as described in the **Suppl. Methods** section. Each estimate of the correlation of SNP effect is based on SNPs reaching marginal genome-wide significance in any of the 5 ancestries analysed. Standard errors for *r(b)* were obtained using jackknife. Error bars denote standard errors of SNP effects. Error bars denote standard errors of SNP effects.

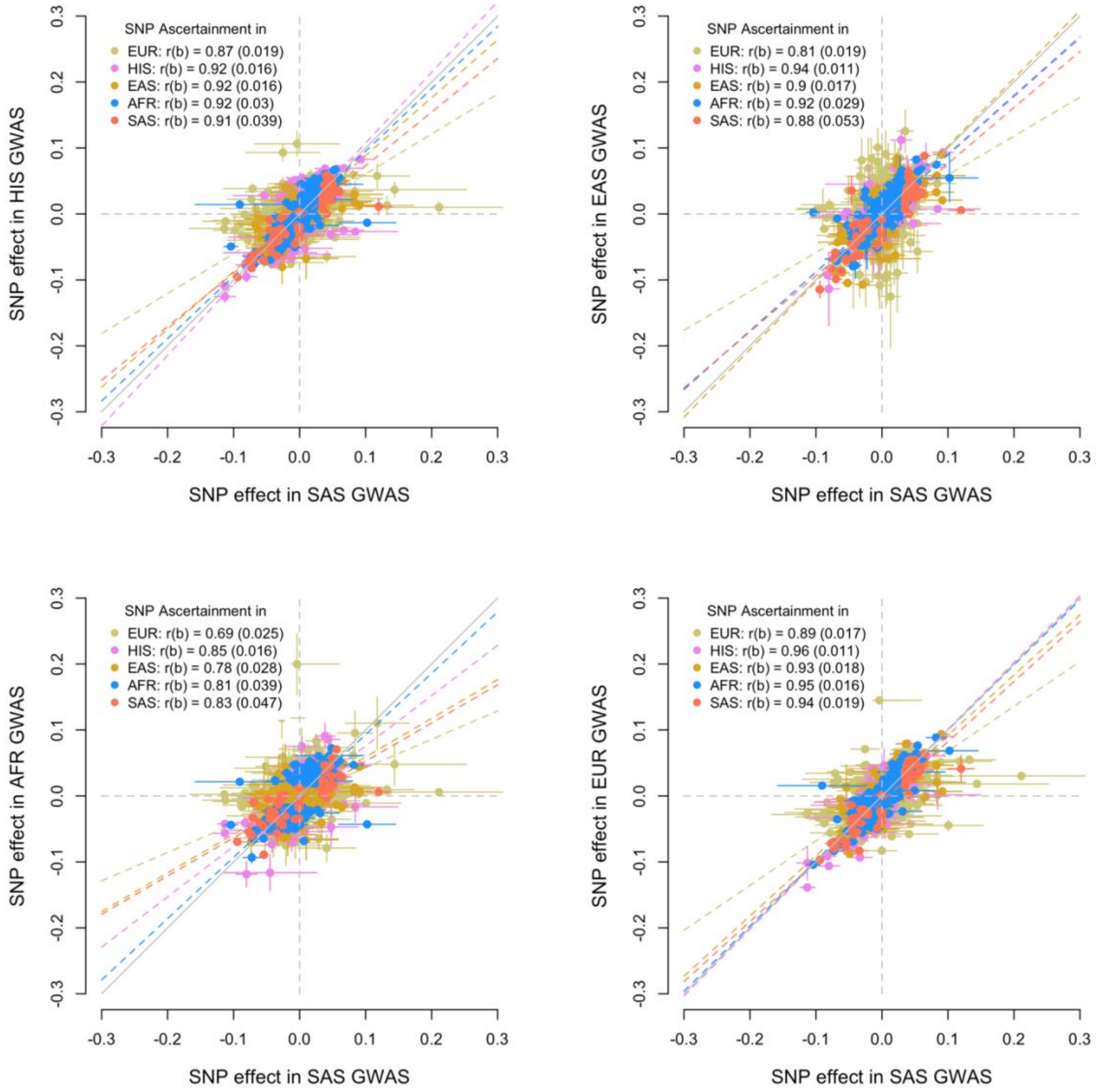


Suppl. Fig. 9. **Correlation of marginal SNP effects between South-Asian ancestry (SAS, on the x-axis) and non-SAS GWAS** (i.e. Hispanic (HIS), African (AFR), East-Asian (EAS) and European (EUR) on the y-axis). Correlations *r(b)* were for corrected for estimation errors as described in the **Suppl. Methods** section. Each estimate of the correlation of SNP effect is based on SNPs reaching marginal genome-wide significance in any of the 5 ancestries analysed. Standard errors for *r(b)* were obtained using jackknife. Error bars denote standard errors of SNP effects.

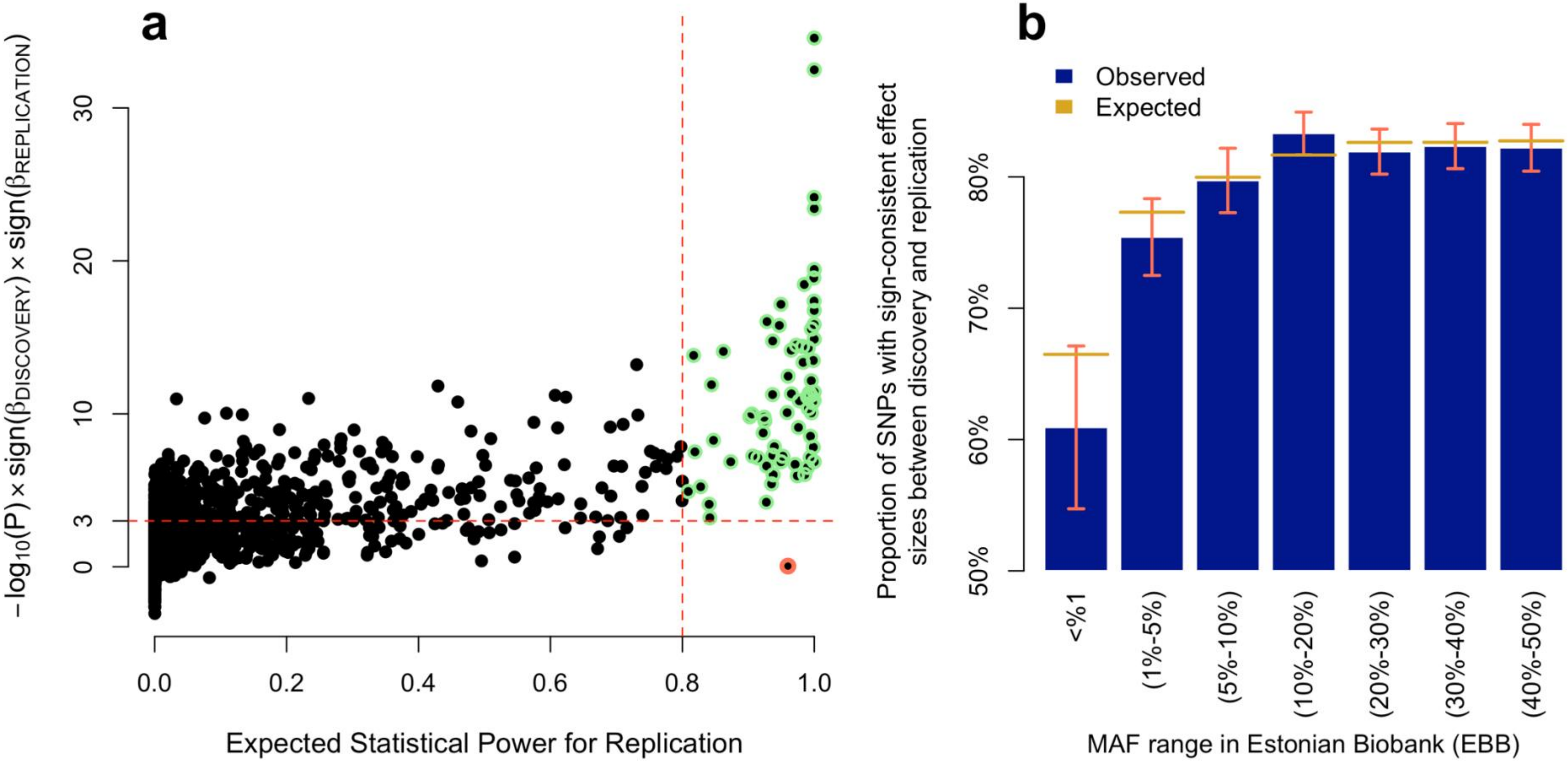


**Suppl. Fig. 10. Replication of marginal associations in the Estonian Biobank (EBB)**. **Panel a**: each dot represents one the 12,111 SNPs detected in our trans-ancestry meta-analysis. The x-axis represents the expected statistical power to replicate each association ($P$<0.05/9,473=5.3×10$^{-6}$; where 9,473 is the number of associations reaching marginal genome-wide significance in our discovery trans-ancestry GWAS and with a minor allele frequency>1% in the EBB sample). The y-axis represents the -log$_{10}$ of the association p-value in the EBB multiplied by the product of signs of estimated SNP effects in the discovery and in the EBB. Horizontal dotted line represents replication at $P$<0.001 and the vertical dotted line indicates 80% of statistical power. One outlier (rs11100870), highlighted in red, does not replicate in the EBB sample. **Panel b**: proportion ($P$) of SNPs with a sign-consistent estimated effect between discovery GWAS (N~5.3M) and EBB. Expected proportions ($E[P]$) are calculated using Equation (2) in the **Suppl. Methods** section. Error bars are defined as $1.96\times\sqrt{P(1-P)/m}$, where $m$ is the number of SNPs in the corresponding MAF interval.

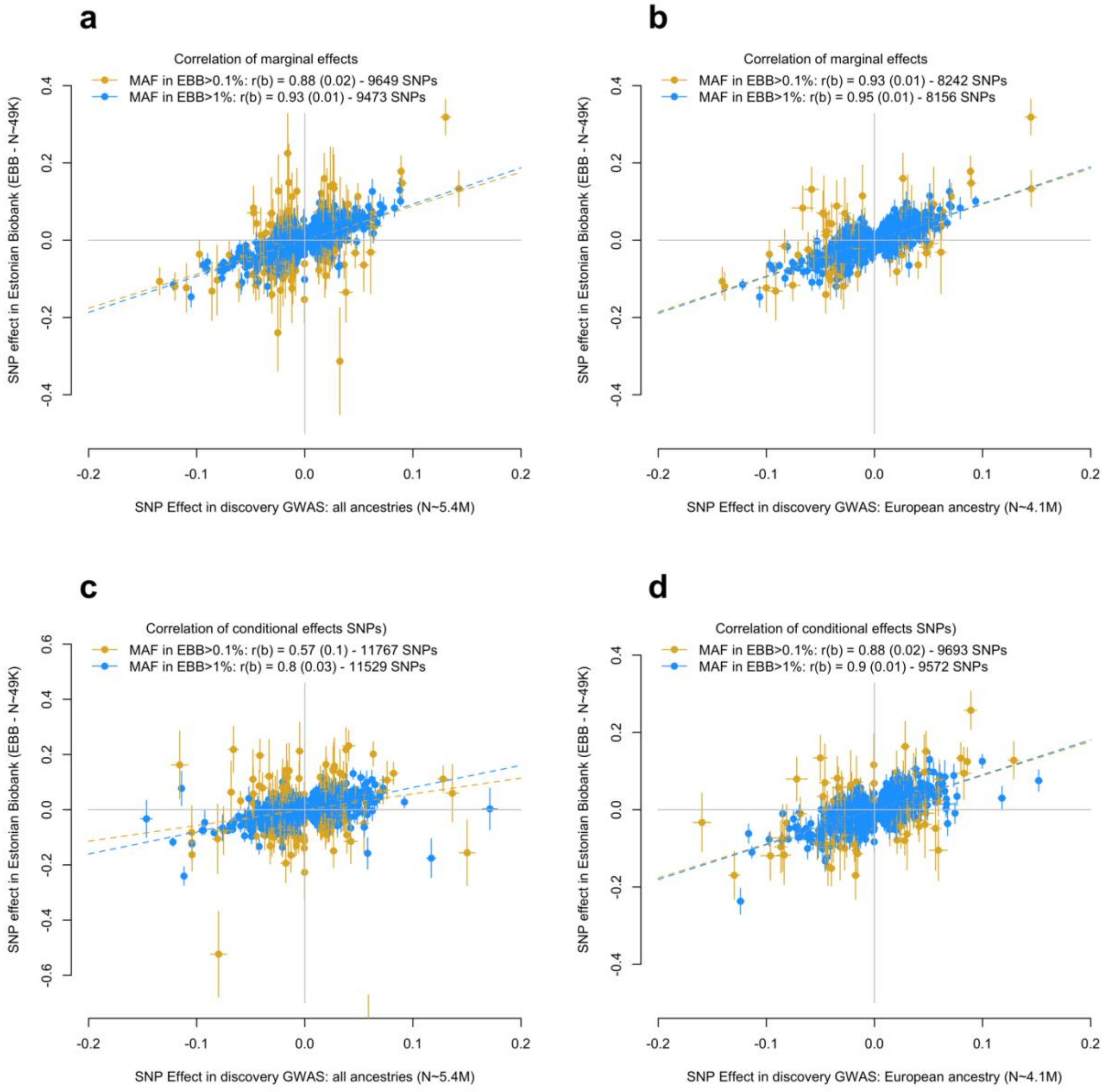


**Suppl. Fig. 11. Correlation of marginal and conditional SNPs effects between discovery and replication (Estonian Biobank – EBB) GWAS. Panels a** and **b** show correlations of marginal SNP effects. **Panels c** and **d** represent joint effect re-estimated using approximate conditional analyses (implemented in the GCTA software). Genotypes of ~350,000 unrelated participants of the UK Biobank were used as linkage disequilibrium (LD) reference.

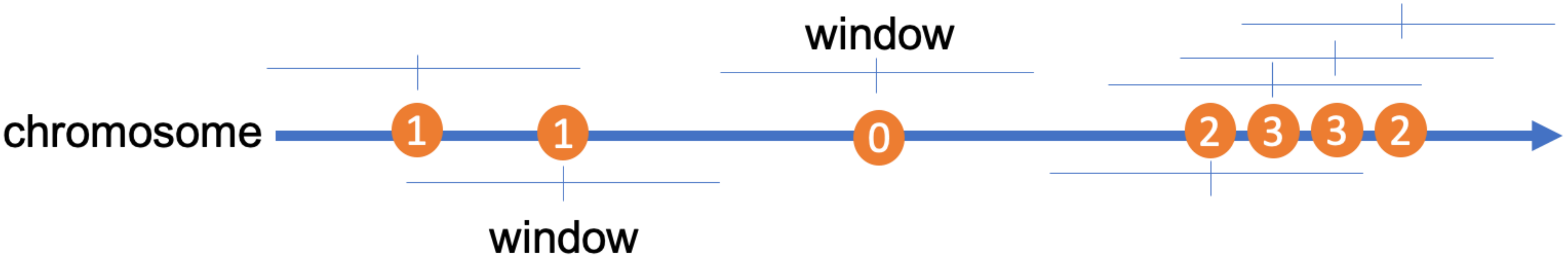


**Suppl. Fig. 12. Schematic representation of the measure of signal density.** The horizontal arrow represents a chromosomes and each circle a specific association. For each association, the density is defined as the number of other independent associations within a certain window. In the example above, the window around the first SNP contains 1 SNP, so its density is 1. Similarly, the density at the third SNP (from the left) is 0 because the window around it does not contain any other association.

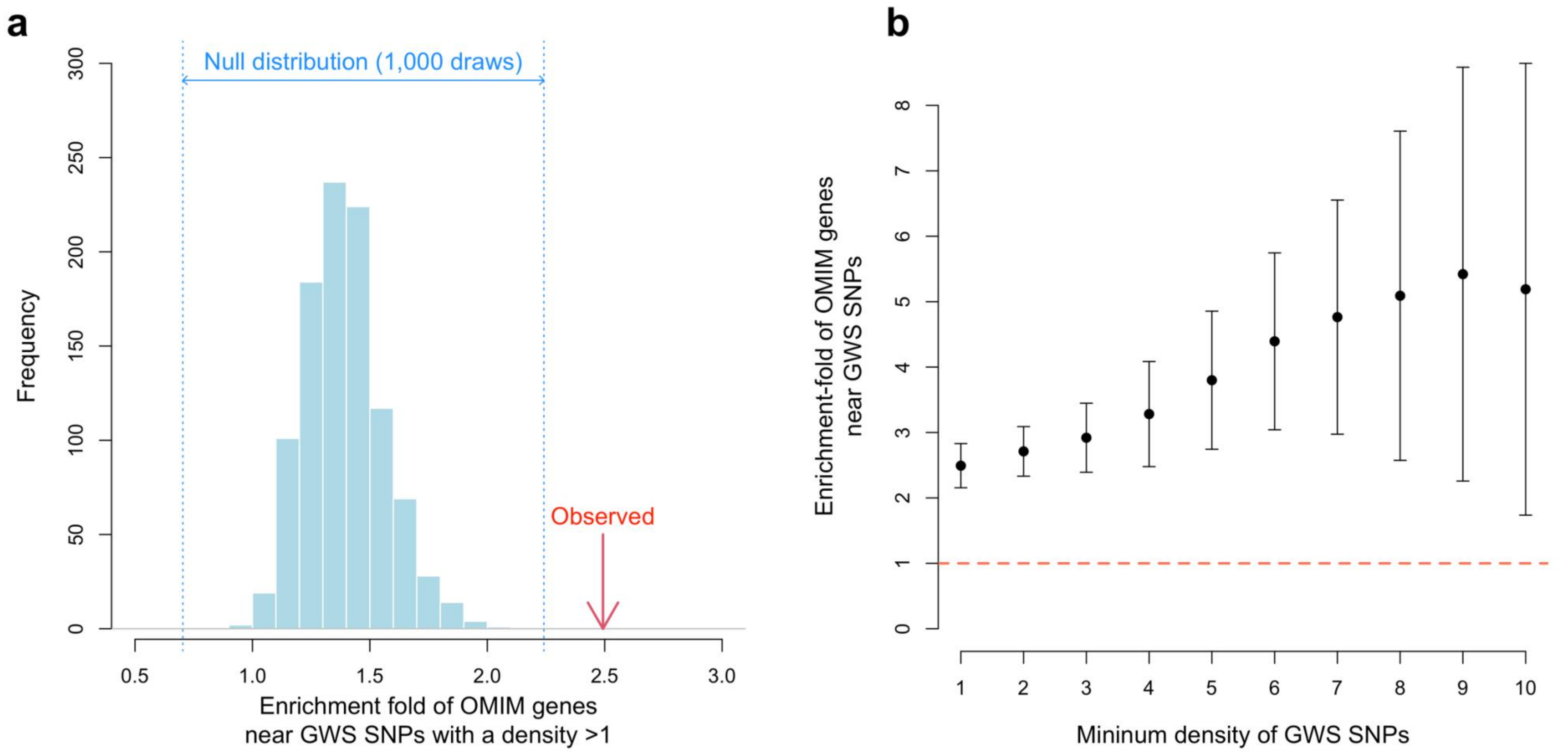


**Suppl. Fig. 13. Enrichment of genes harbouring pathogenic mutations causing extreme height or abnormal skeletal growth syndromes near hotspots of genome-wide significant (GWS) SNPs**. Four hundred and sixty-two (462) autosomal genes were curated from the Online Mendelian Inheritance in Man (OMIM) database. Panel **a**. Red arrow indicates the observed enrichment statistic (OR=2.5-fold) measuring the odds ratio of the presence of an OMIM gene within 100 kb of a GWS SNPs with a density > 1. The blue histogram represents the distribution of enrichment statistics from 1,000 random genes matched, which length distribution matches that of the OMIM genes. Panel **b**: Enrichment of OMIM genes near high density GWS SNPs. High density is defined by on the x-axis by the minimum number of other independent GWS SNPs detected within 100 kb.

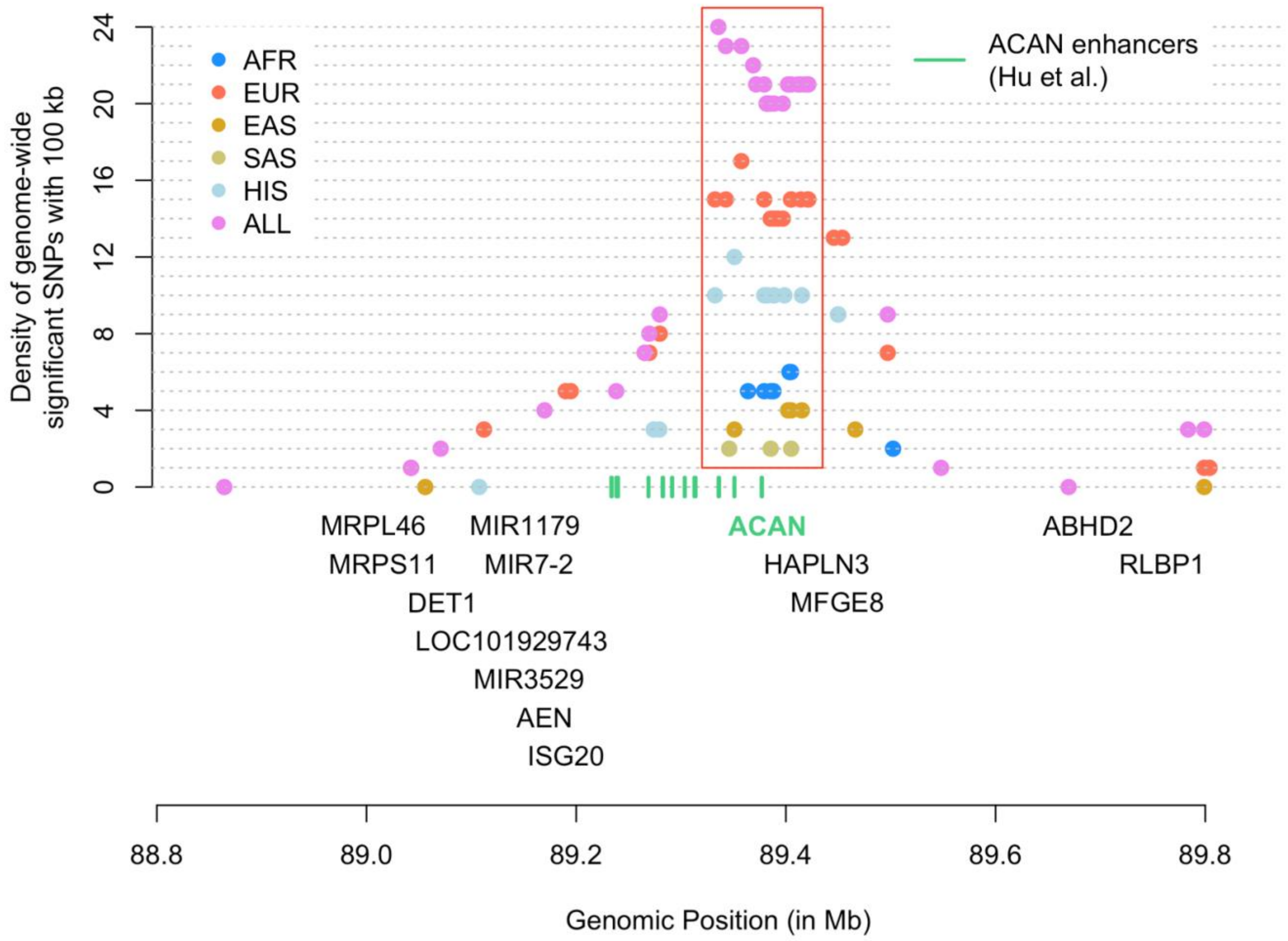


**Suppl. Fig. 14. Independent signal density at the *ACAN* gene locus across ancestries**. Independent associations were identified from GWAS performed in 5 ancestries (African: AFR; European: EUR; East-Asian: EAS; South-Asian: SAS and Hispanic: HIS) as well as from the meta-analysis of all ancestries (ALL). Genomic segments with a signal density >1 are found in each ancestry group.

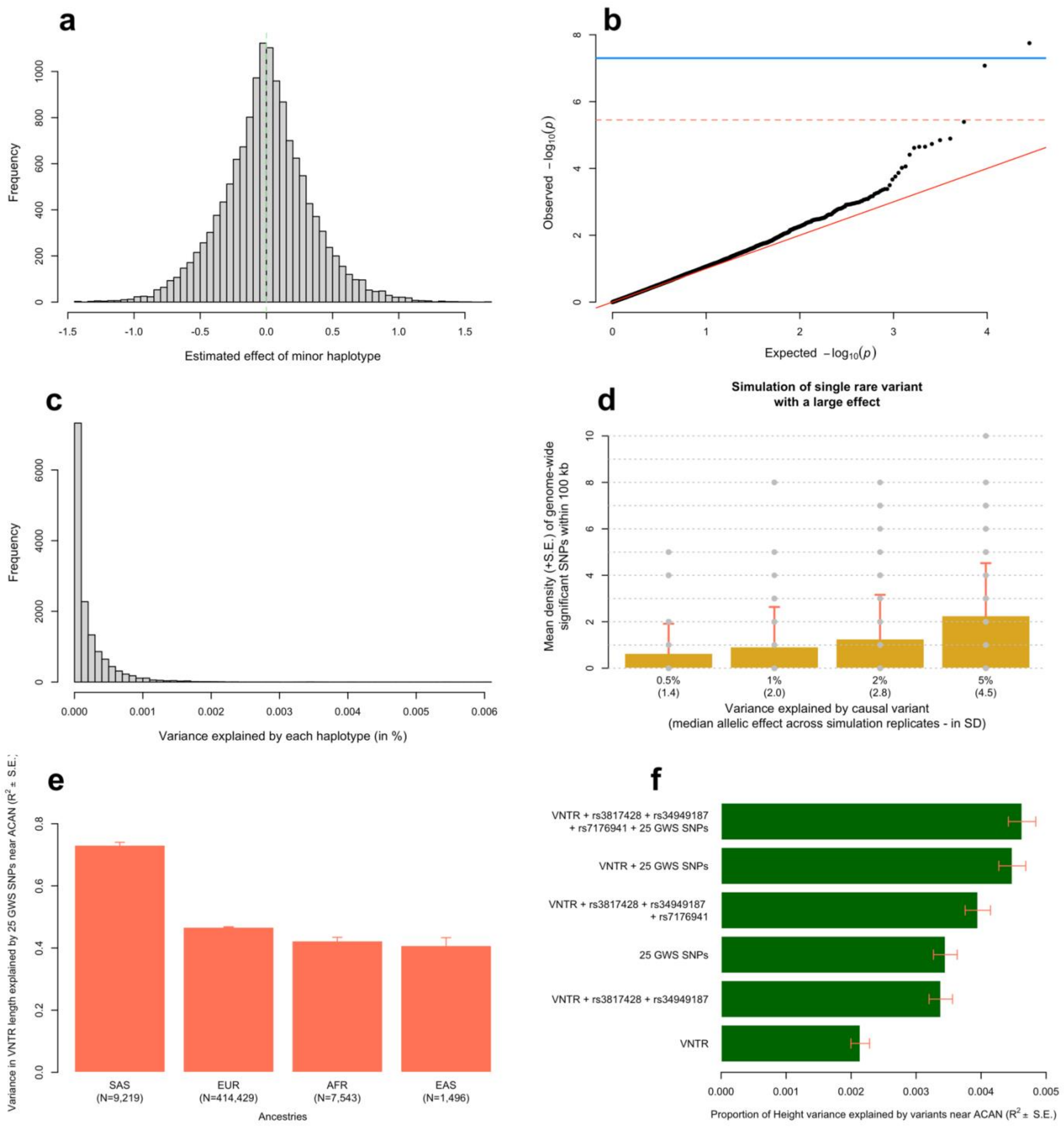


**Suppl. Fig. 15. Haplotypic analysis at the *ACAN* locus. Panel a** shows the distribution of estimated haplotype effects from 14,117 haplotypes covering a 100 kb long genomic region near the *ACAN* gene (hg19 genomic coordinates: chr15:89,307,521-89,407,521). **Panel b** show the quantile-quantile plot of associations between these 14,117 haplotypes and height. **Panel c** shows the distribution of the variance explained by each of the 14,117 haplotypes. **Panel d** shows the mean signals density (y-axis) across simulated data where 1 causal SNP within the locus explains between 0.5% and 5% (x-axis) of trait variance. Causal variants were sampled from a pool of 13 SNPs with a $1.4\times10^{-5}<MAF<1\%$ genotyped in 291,683 unrelated EUR participants of the UKB, with no missing values at these 13 SNPs. S.E. were calculated as the standard deviation (SD) of signal density across 100 simulation replicates. GCTA-COJO analyses to identify independent signals were performed using a subset of 10,000 unrelated EUR participants of the UKB to mimic the large discrepancy between the size of the discovery GWAS and that of the LD reference used in our real data analyses. **Panel e** shows the proportion of variable-number-tandem-repeat (VNTR) length explained by 25 genome-wide significant (GWS) SNPs identified near ACAN in 4 ancestries (European: EUR; South-Asian: SAS; East-Asian: EAS; African: AFR). **Panel f** shows the proportion of height variance explained in a sample of EUR UK Biobank participants by various sets of polymorphisms at the ACAN locus. rs3817428 and rs34949187 are two missense variants and rs7176941 is an intronic variant with high posterior causal probability identified in ref.[28] In panels **e** and **f**, error bars represent standard errors (S.E.).

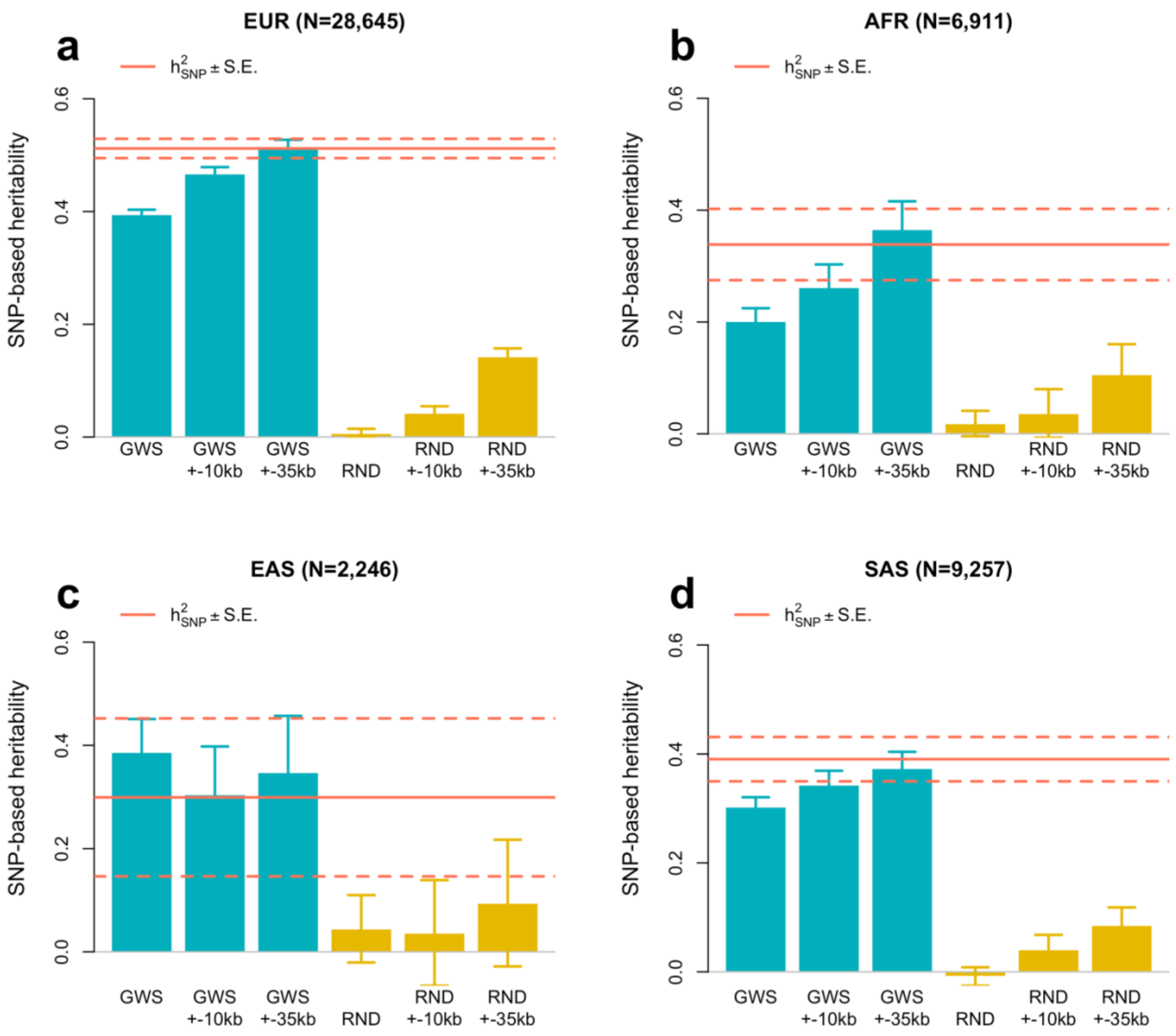


**Suppl. Fig. 16. Variance of height explained by SNPs in genome-wide significant (GWS) loci defined with various window sizes.** Stratified SNP-based heritability ($h^2_{\mathrm{SNP}}$) estimates were obtained for three partitions of the genome: (1) GWS SNPs alone vs. all other HapMap 3 (HM3) SNPs; (2) GWS SNPs +/- all HM3 SNPs within 10 kb vs. all other HM3 SNPs and (3) GWS SNPs +/- all HM3 SNPs within 35 kb vs. all other HM3 SNPs. Analyses were performed in samples of four different ancestries: European (EUR: meta-analysis of UK Biobank (UKB); N=14,587 + Lifelines data; N=14,058), African (AFR: UKB), East-Asian (EAS: UKB) and South-Asian (SAS: UKB). Estimates from stratified analyses were compared with SNP-based heritability estimates obtained from analysing all SNPs jointly (horizontal red bar; dotted lines represented standard errors). Analyses were repeated using a random set of 12,111 SNPs (and redefining loci relative to those), which minor allele frequency and linkage disequilibrium distribution matched that of GWS SNPs (RND: gold bars).

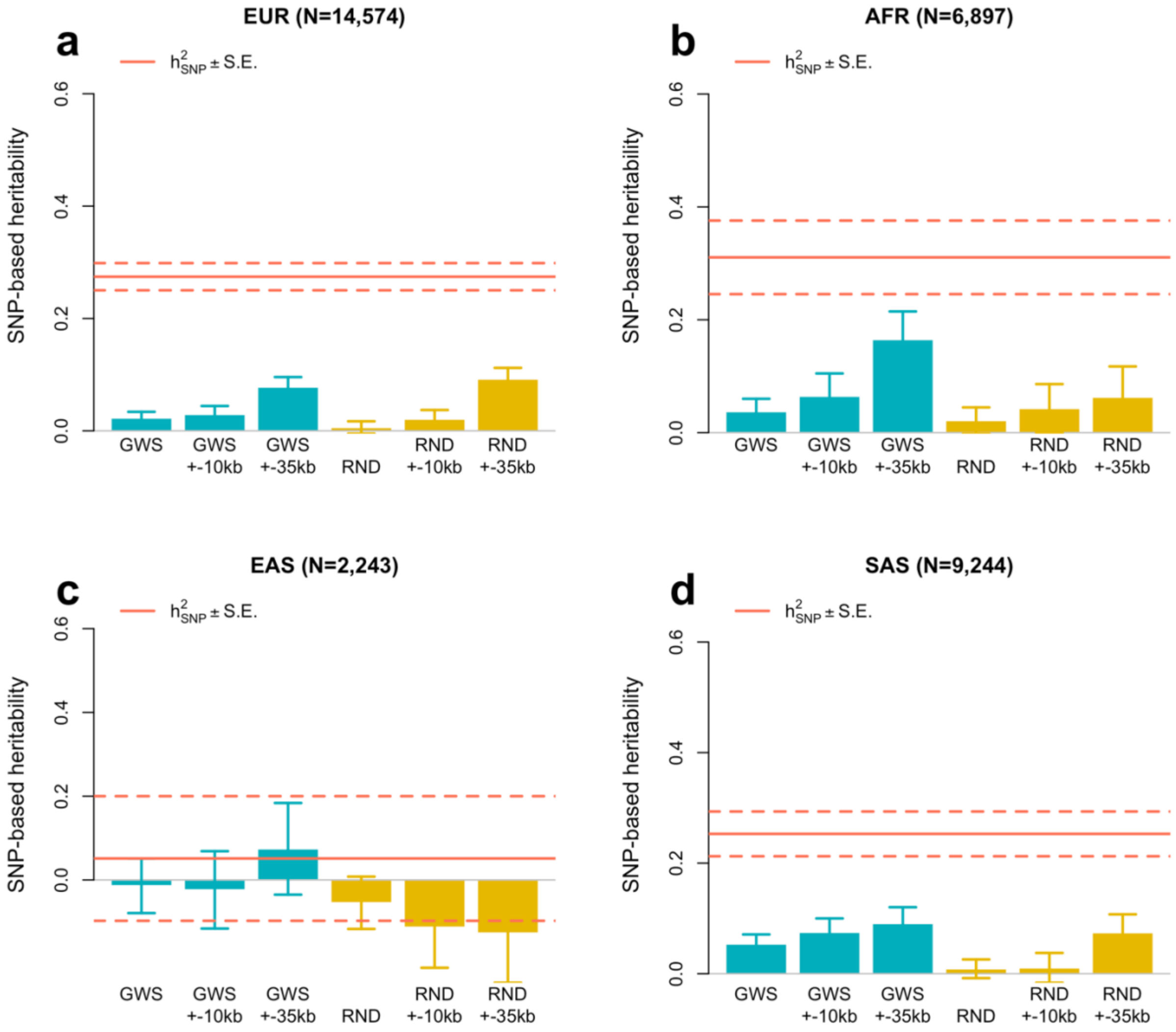


**Suppl. Fig. 17. Variance of body mass index (BMI) explained by height-associated genome-wide significant (GWS) loci defined with various window sizes.** Stratified SNP-based heritability ($h^2_{\mathrm{SNP}}$) estimates were obtained for three partitions of the genome: (1) GWS SNPs alone vs. all other HapMap 3 (HM3) SNPs; (2) GWS SNPs +/- all HM3 SNPs within 10 kb vs. all other HM3 SNPs and (3) GWS SNPs +/- all HM3 SNPs within 35 kb vs. all other HM3 SNPs. Analyses were performed in UK Biobank samples of four different ancestries: European (EUR), African (AFR), East-Asian (EAS) and South-Asian (SAS). Estimates from stratified analyses were compared with SNP-based heritability estimates obtained from analysing all SNPs jointly (horizontal red bar; dotted lines represented standard errors). Analyses were repeated using a random set of 12,111 SNPs (and redefining loci relative to those), which minor allele frequency and linkage disequilibrium distribution matched that of GWS SNPs (RND: gold bars).

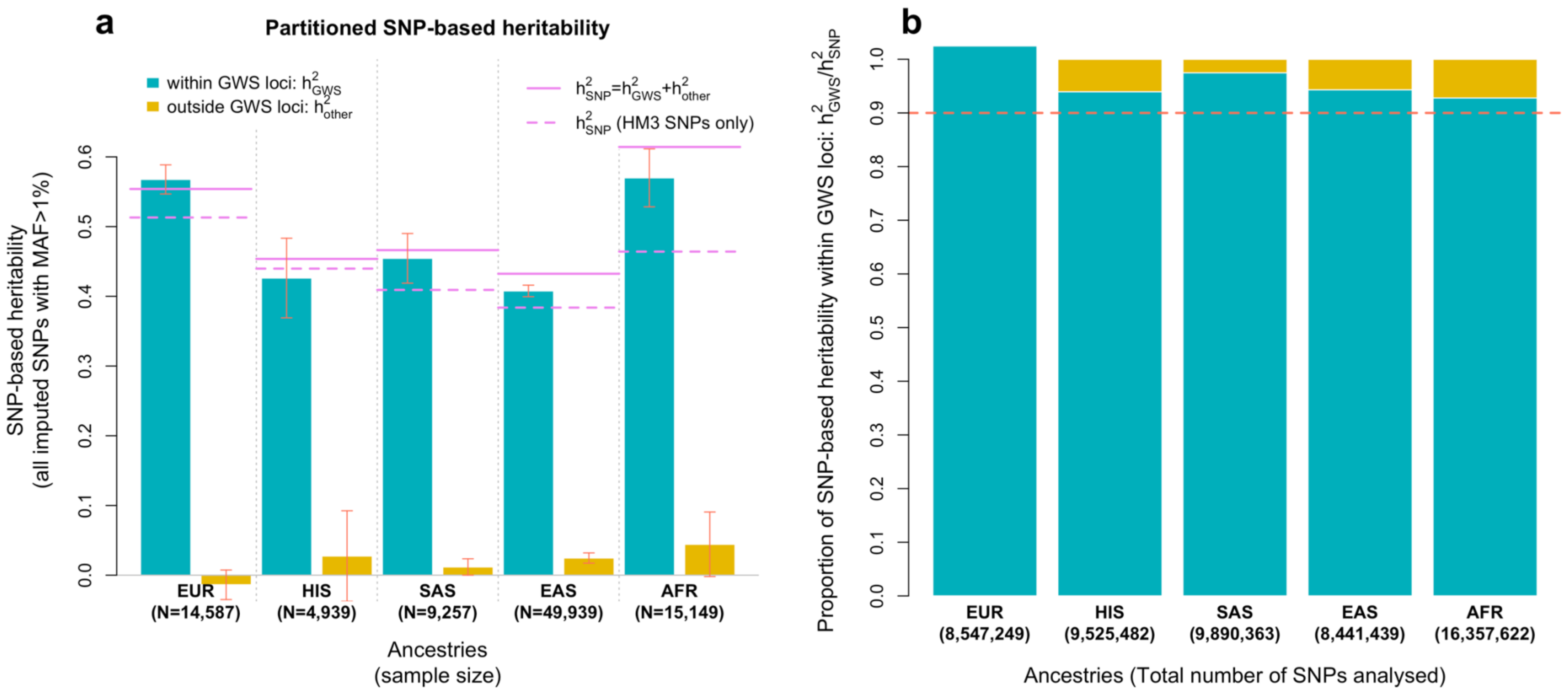


**Suppl. Fig. 18. Variance of height explained by common SNPs within 35 kb of genome-wide significant (GWS) SNPs.** Stratified SNP-based heritability ($h^2_{SNP}$) estimates were obtained from a partition of the genome into two sets of 1,000 Genomes imputed SNPs with a minor allel e frequency (MAF) >1%: (1) SNPs within +/- 35 kb of GWS (GWS loci) vs. all other SNPs. Analyses were performed in samples of five different ancestry groups: European (EUR; UK Biobank only), African (AFR), East-Asian (EAS) and South-Asian (SAS) as described in the legend of Fig. 2. Estimates from stratified analyses were compared with SNP-based heritability estimates obtained from analysing HapMap 3 (HM3) SNPs only (dotted horizontal violet bar).

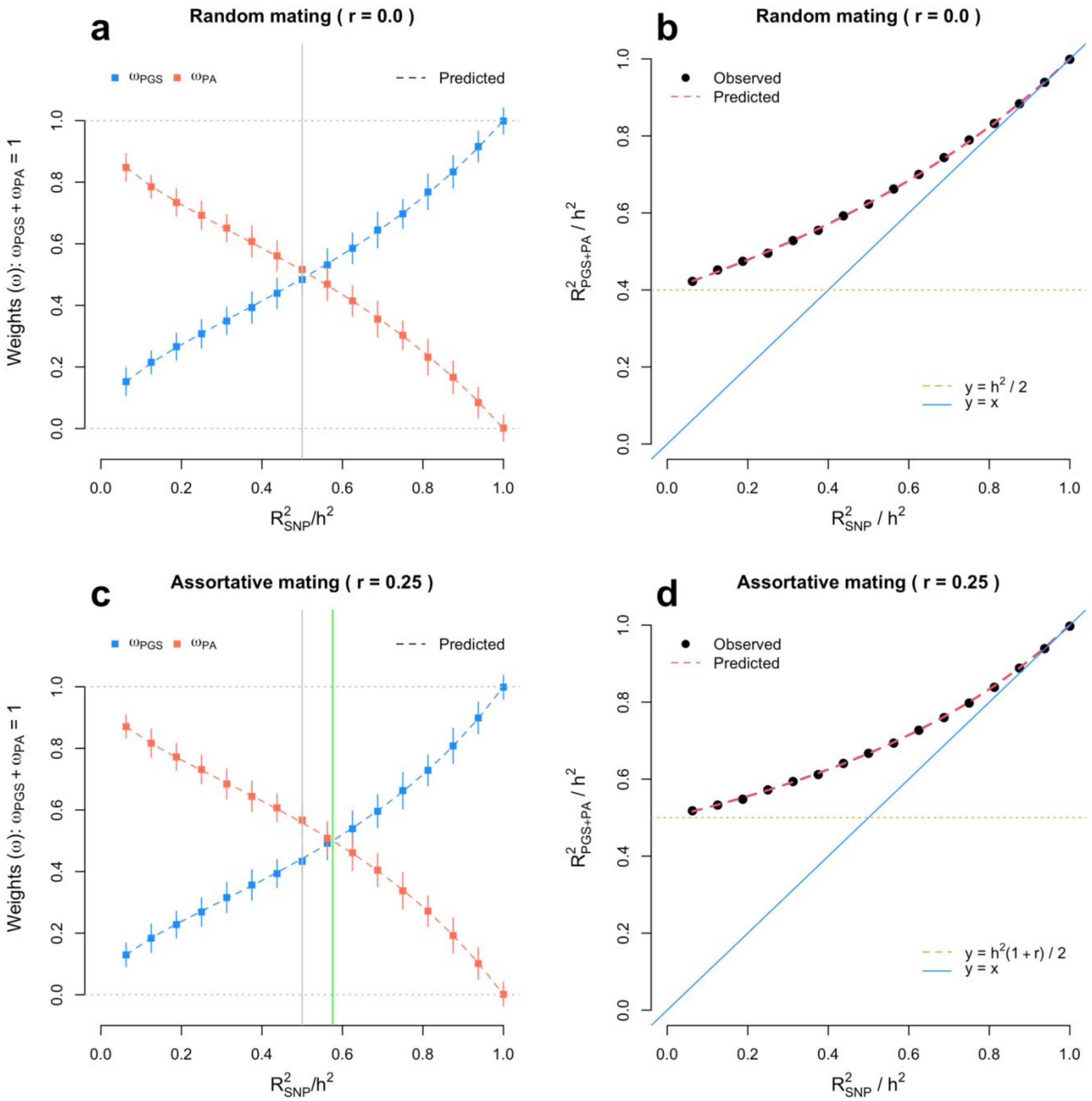


**Suppl. Fig. 19. Optimal weighting of PGS and parental information in simulated data.** We simulated a population of N=2,000 individuals and a trait controlled by M=1,000 causal variants. For simplicity, we assumed that a variable proportion of causal SNPs is used to calculate the PGS, and that SNP effects are estimated with negligible errors. We show below in Equation (3.11) how this proportion is chosen to achieve the desired prediction accuracy. We considered two scenarios: (i) random mating, i.e. r=0 (**Panels a** and **b**) and assortative mating (for 20 generations) based on a spousal phenotypic correlation r=0.25 (**Panels c** and **d**). In all simulations, we assumed a heritability $h^2 = 0.8$ and varied the expected prediction accuracy ($R^2_{SNP}$) between 0.05 and 0.8. For each simulated population, we compared our predictions from Equation (S3.1) and (S3.2) with estimated regression coefficients obtained from regressing y on $\hat{y}$ and $\bar{y}_p$. The vertical green bar in panel **c**, denotes the threshold above which PGS information outweighs parental information. This threshold is predicted using Equation (S3.4). We also compared our variance explained by fitting both predictors with our predicted expectation from Equation (S3.3). Each dot is generated using 100 replicates. Overall, we found a perfect consistency between our theoretical and simulation results, which provides an empirical validation of these predictions.

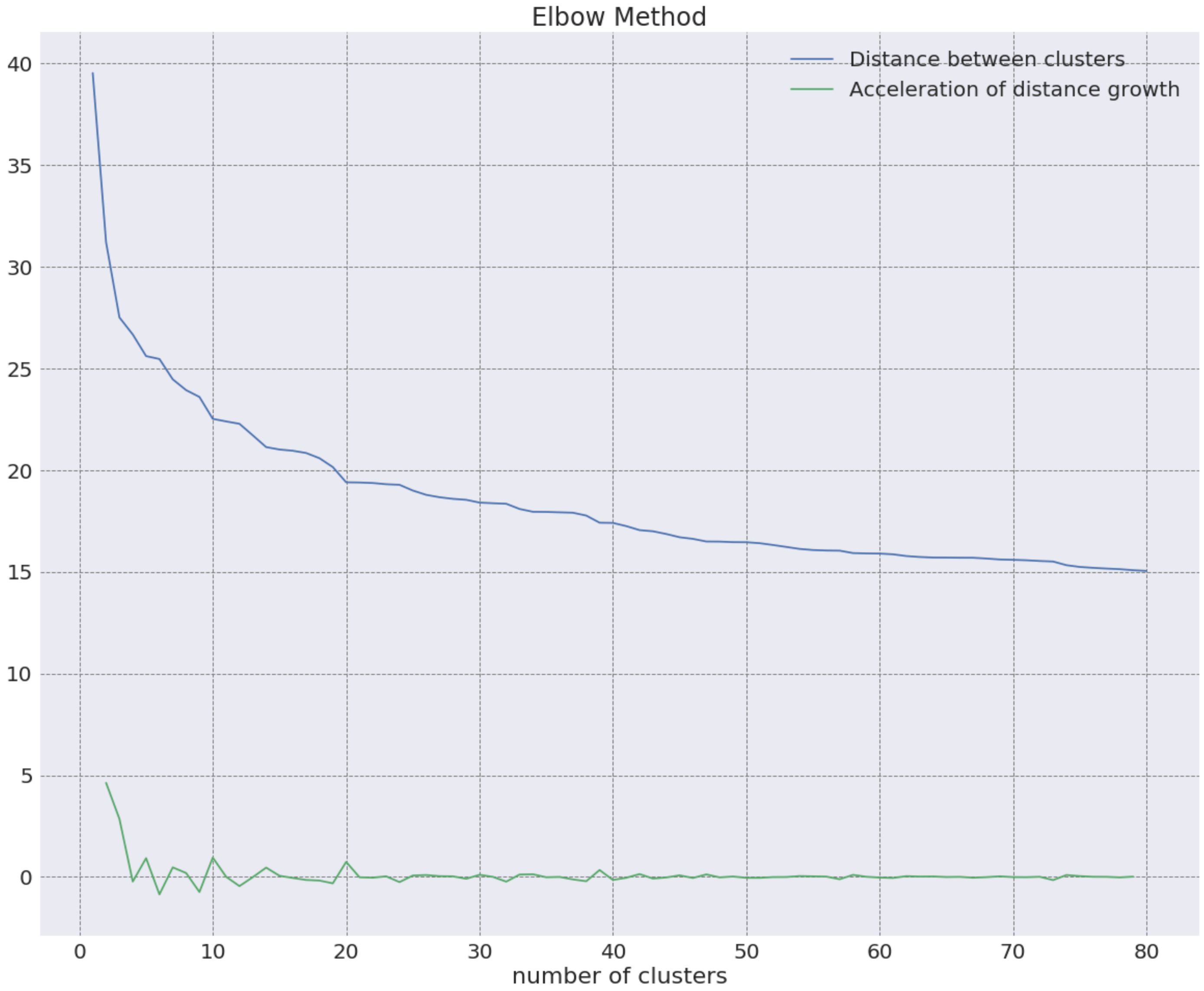


**Suppl. Fig. 20. Selection of the number of gene set clusters using the "Elbow method".** Gene sets were hierarchically clustered at different sizes and the distance between each cluster evaluated. 20 clusters was chosen as an appropriate number of gene set clusters to evaluate for enrichment.

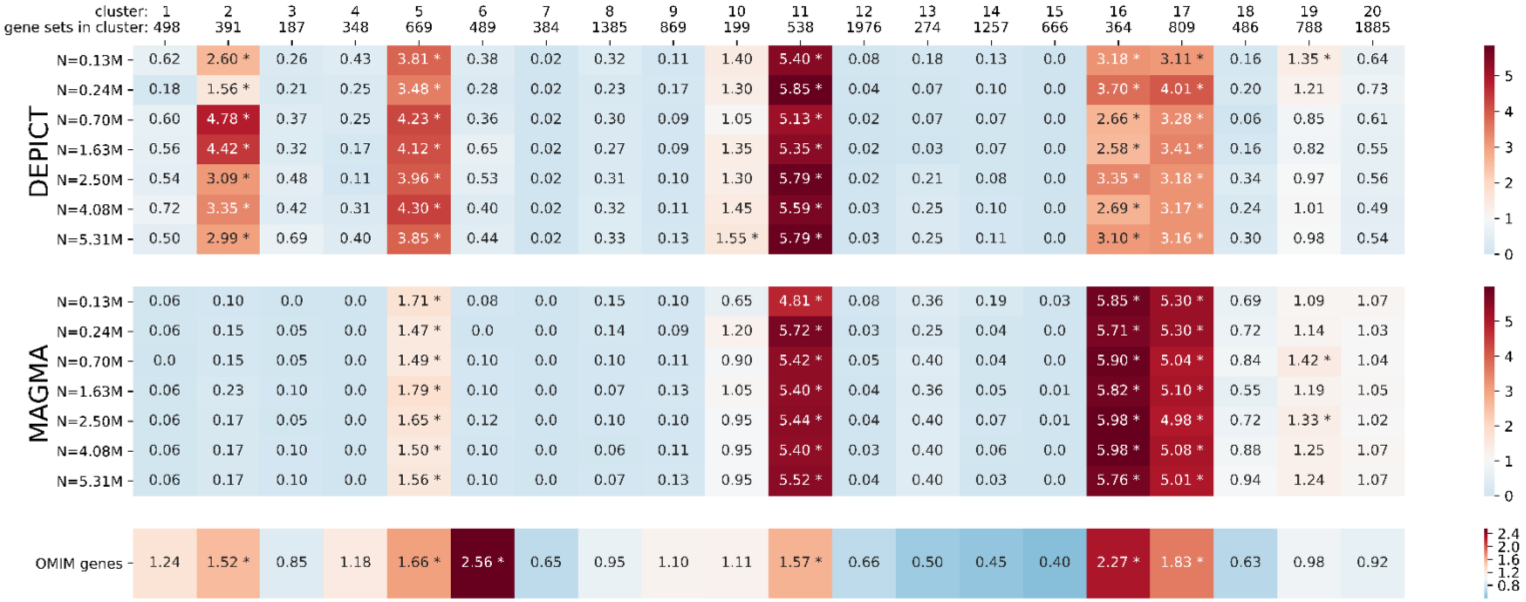


**Suppl. Fig. 21. Enrichment of height-associated genes identified at various GWAS sample size within 20 clusters of gene sets representing broad categories of biological pathways.** Gene set enrichment was performed with MAGMA and DEPICT across seven GWAS with increasing sample sizes. Samples used (Lango-Allen et al., Wood et al., Yengo et al., GIANT-EUR (no 23andMe), 23andMe-EUR, European, and Trans-ancestry meta-analysis) are described in **Tables 1-2**. The degree of enrichment of gene sets (MAGMA, DEPICT) of known skeletal growth disorder genes catalogued in the Online Mendelian Inheritance in Man (OMIM) database among 20 clusters of gene sets (see Methods section in **Suppl. Note 4**) is indicated by the blue-red colour scale. Enrichment for MAGMA and DEPICT was defined to be the number of prioritized gene sets (top 10% of gene sets) in each cluster divided by the 10% of the number of gene sets in the cluster. Enrichment for OMIM was defined to be the number of OMIM genes in a gene set (Z > 1.96) divided by the size of the gene set divided by the proportion of all genes in OMIM, then averaged across the cluster. Significant enrichment (compared to shuffled prioritization of gene sets or genes) is marked with *.

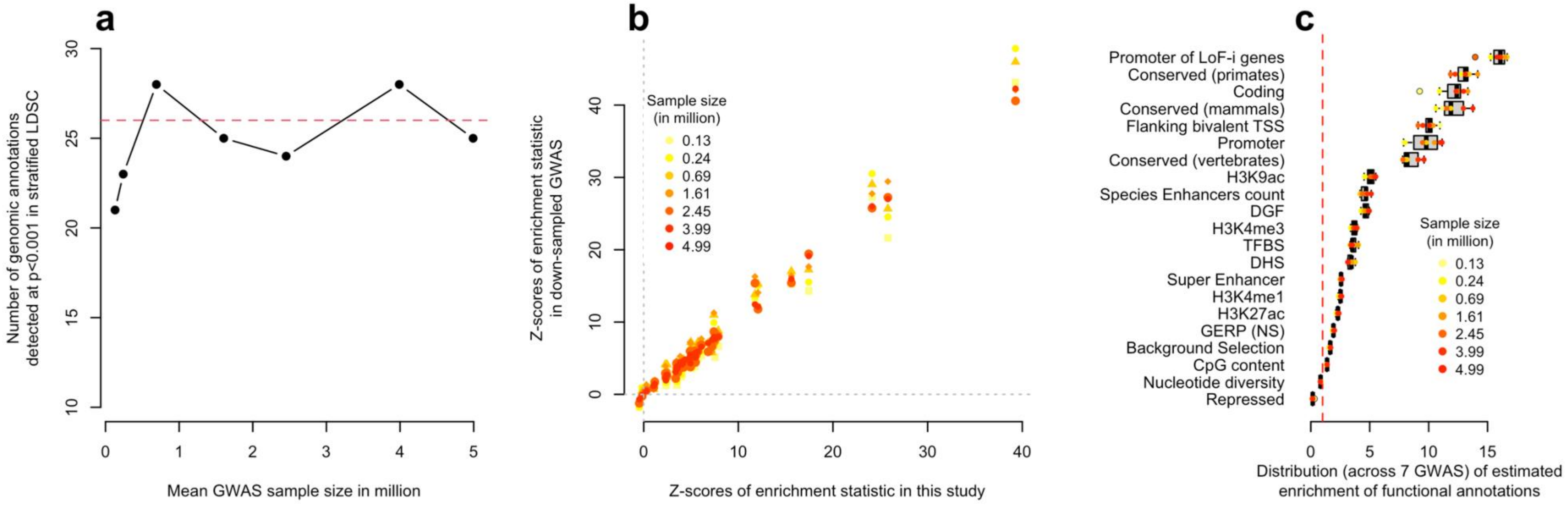


**Suppl. Fig. 22. Annotation-level saturation of GWAS discoveries as a function of sample size.** Increase in sample size from ~4 million to ~5 million is achieved by including ~1 million participants of non-European ancestry. **Panel a** shows the number annotations showing a significant heritability enrichment as function the function of the sample size of the GWAS used to estimate these enrichment. Heritability enrichment was detected using a stratified LD score regression (LDSC) analysis of 97 genomic annotations included in the "baseline + LD" model from Gazal et al. **Panel b** shows the correlation between Z-scores measuring the statistical significance of heritability enrichments of 97 annotations (each dot is an annotation) in our largest GWAS (x-axis) as compared to down-sampled GWAS (*y*-axis). Sample size is denoted by the colour-code. **Panel c** shows the distribution of estimated enrichment statistics for 21 annotations found significantly enriched ($P$<0.05/97) in at least 6 of the 7 GWAS analysed here. LoF-i genes: Loss of function intolerant genes; TSS: Transcription Start Sites; DGF: Digital genomic footprint; TFBS: Transcription Factor Binding Sites; DHS: DNAse I hypersensitive sites; GERP (NS): GERP++ score (number of substitutions).

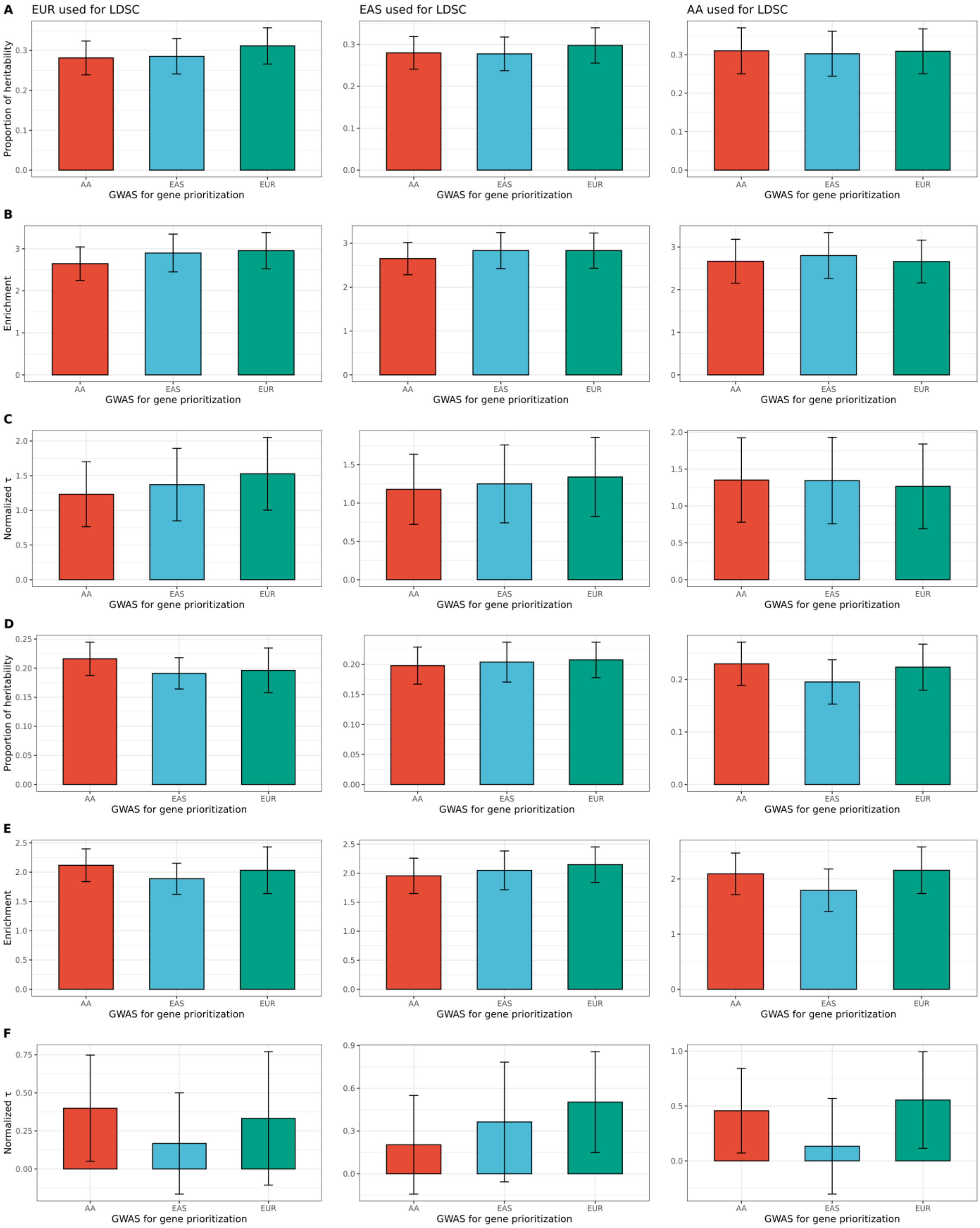


**Suppl. Fig. 23. Proportional heritability ($h^2$) explained, enrichment, and normalized $\tau$ estimates for MAGMA (panels A-C) and DEPICT (panels D-F).** The error bars represent the 95% confidence interval, calculated as estimate +/- 1.96*standard error. The labels for each subpanel indicate the ancestry represented in the GWAS used for LDSC, and the x-axis labels indicate the ancestry represented in the "discovery" GWAS used to prioritize the genes. EUR: European ancestry; AA: African-Americans of admixed European and African ancestries; EAS: East Asian ancestry.

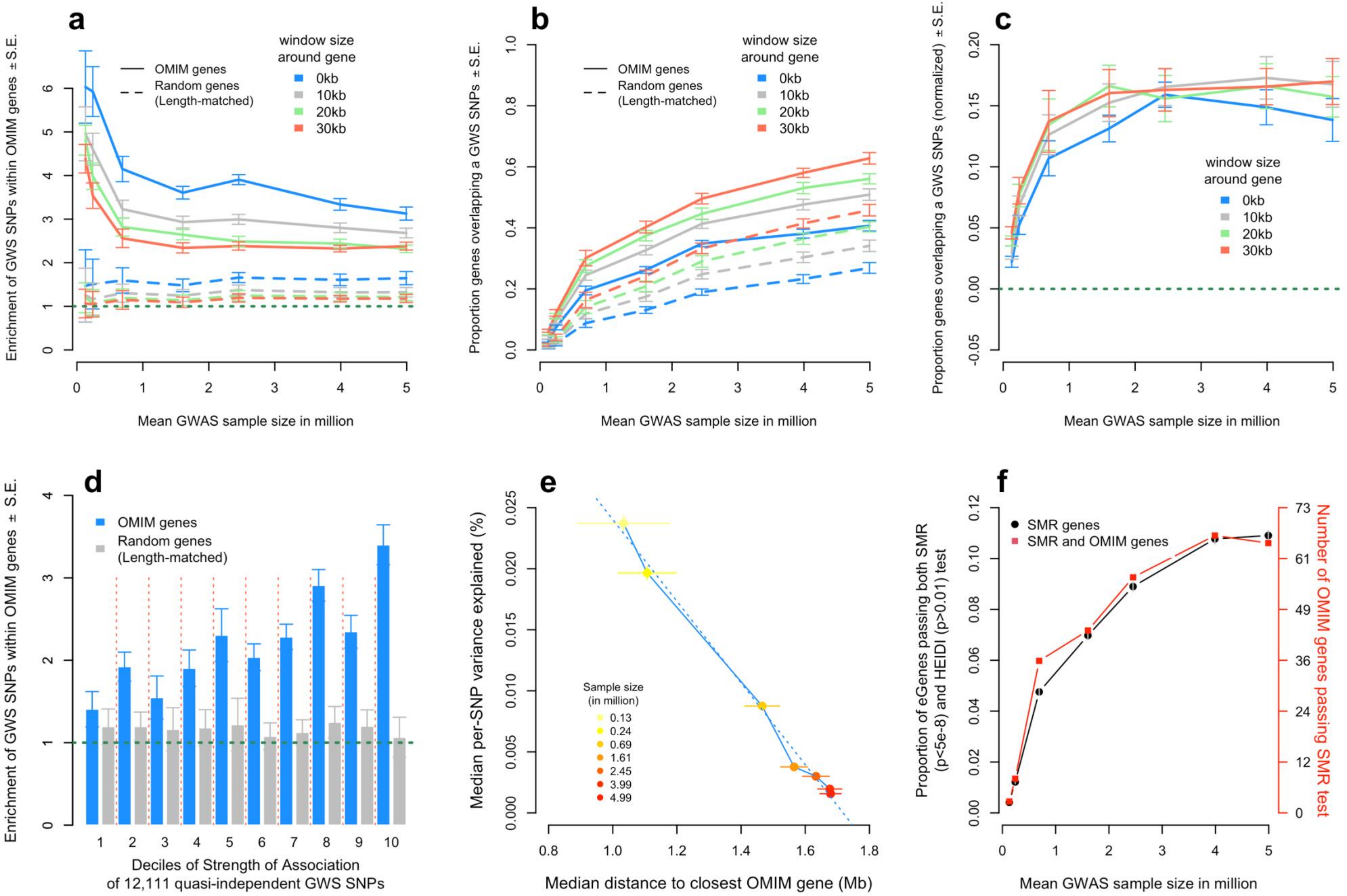
a
Enrichment of GWS SNPs within OMIM genes ± S.E.
OMIM genes
Random genes (Length-matched)
window size around gene
0kb
10kb
20kb
30kb
Mean GWAS sample size in million
b
Proportion genes overlapping a GWS SNPs ± S.E.
c
Proportion genes overlapping a GWS SNPs (normalized) ± S.E.
d
Deciles of Strength of Association of 12,111 quasi-independent GWS SNPs
e
Median per-SNP variance explained (%)
Median distance to closest OMIM gene (Mb)
Sample size (in million)
0.13
0.24
0.69
1.61
2.45
3.99
4.99
f
SMR genes
SMR and OMIM genes
Proportion of eGenes passing both SMR (p<5e-8) and HEIDI (p>0.01) test
Number of OMIM genes passing SMR test

**Suppl. Fig. 24. Gene-level saturation of GWAS discoveries as a function of sample size.** Increase in sample size from ~4 million to ~5 million is achieved by including ~1 million participants of non-European ancestry. **Panel a** shows the enrichment of genome-wide significant (GWS) SNPs identified from an approximate conditional and joint (COJO) analysis within 462 genes associated with skeletal growth disorders from the Online Mendelian Inheritance in Man (OMIM) database (y-axis) as a function of GWAS sample size (x-axis). Standard Error (S.E.) were calculated as the standard deviation of enrichment statistics (odds ratio in the 2x2 contingency table contrasting for each gene: "is the gene an OMIM gene" vs. "does the gene contain a GWS SNP") across 1,000 randomly sampled sets of 462 non-OMIM genes length-matched with OMIM genes. The average enrichment calculated across the 1,000 random gene sets is represented with dotted lines. Presence of a GWS SNP within a gene was assessed relative to gene start and stop position, considering flanking regions within 0 kb, 10 kb, 20 kb and 30 kb. **Panel b** shows the proportion of OMIM overlapping genes with at least one GWS SNPs (y-axis). As in **Panel a**, dotted lines represents the null distribution from 1,000 random sets of genes length-matched with OMIM genes. Standard errors (S.E.) were calculated as the standard deviation of the proportion observed across the 1,000 draws from the null distribution. **Panel c** represents the proportion of OMIM genes near GWS SNPs after subtracting the mean of the null distribution at each sample size. **Panel d** represents the enrichment of OMIM genes as a function of the strength of association of 12,111 independent GWS SNPs identified in our largest GWAS (N~5.4M). GWS SNPs were grouped into 10 decile groups of ~1,211 SNPs. Enrichment near OMIM genes is stronger of SNPs explaining a larger proportion of height variance (top decile). **Panel e** shows the median per-SNP variance explained (y-axis) as a function of the median distance to the closest OMIM gene. Large GWAS tend to identify variants with smaller effect sizes and further away from OMIM genes. **Panels f** shows the number of genes prioritised using Summary-data based Mendelian Randomization (SMR; $P<5\times10^{-8}$), which expression may act a mediator of the effects of SNP on height. SMR analyses were based on expression quantitative trait loci (eQTL) identified in the GTEx and eQTLgen studies (**Suppl. Methods**). The z-axis (in red) shows the number of OMIM genes overlapping with SMR genes identified from analysing GWAS with various sample sizes (x-axis).

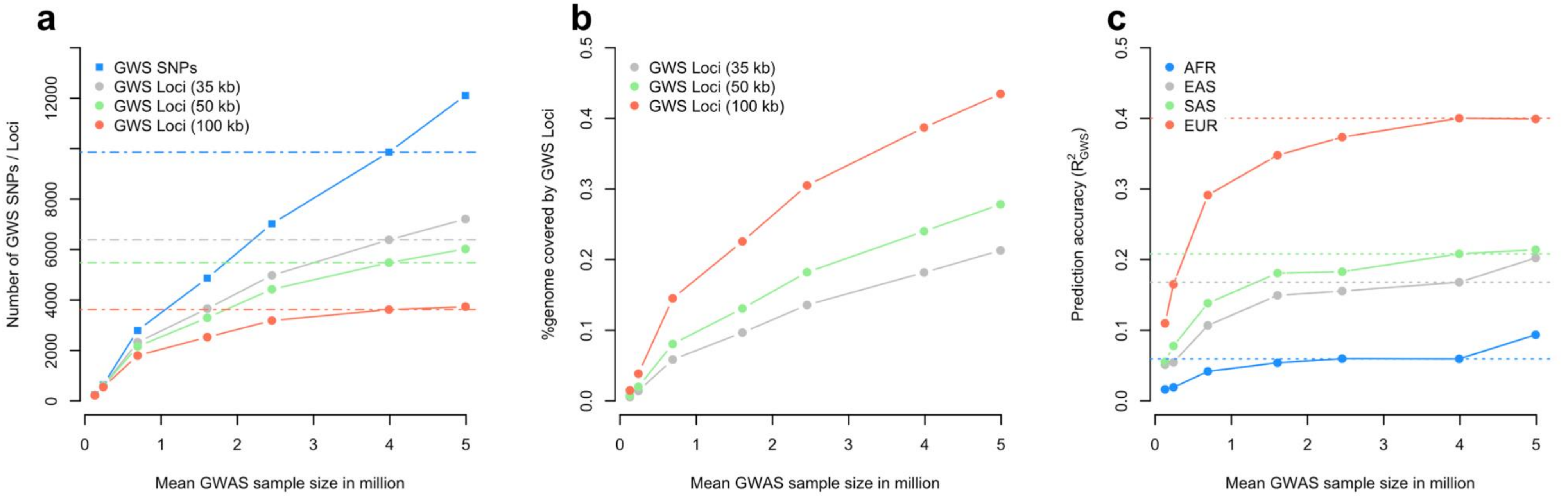


**Suppl. Fig. 25. Variant-level saturation of GWAS discoveries as a function of sample size.** Increase in sample size from ~4 million to ~5 million is achieved by including ~1 million participants of non-European ancestry. **Panel a** shows number of independent genome-wide significant (GWS) SNPs and loci identified at various GWAS sample sizes (Details about down-sampled GWAS are given in Table 2). GWS loci were defined using various window sizes including 35kb, 50kb and 100 kb. **Panel b** shows the percentage of the genome covered by GWS loci. Coverage was calculated as the cumulative length of GWS loci in Mb divided 3,039 Mb, the estimated length of the human genome. **Panel c** shows the prediction accuracy ($R^2_{\mathrm{GWS}}$) of various polygenic scores on GWS SNPs identified at various sample sizes. In **Panels a** and **c**, dotted lines represent y-axis values for our largest European ancestry GWAS (N~4 million).

### SUPPLEMENTARY NOTES

#### Supplementary Note 1: Investigation of population stratification in a large European ancestry GWAS of height

Recent studies[23,24] have shown evidence of significant confounding in estimated SNP effects induced by uncorrected population stratification (PS) in summary statistics from the Wood et al.[4] study. Importantly, Wood et al. also reported residual PS mostly affecting estimated effects of SNPs weakly associated with height (Figure 2a-c in ref.[4]). To ensure increased robustness and reliability of our findings, we perform here a series of analyses to quantify confounding effects of residual PS in our European ancestry (EUR) GWAS.

##### 1.1 LD score regression analysis

First, we performed a LD score regression (LDSC) analysis[25] of our GWAS meta-analysis of EUR participants (N=4,080,687). We assessed the degree of PS using the attenuation ratio statistic ($R_{LDSC}$), which provides a quantification of PS that is independent of sample size. The estimated $R_{LDSC}$ is ~3.8% (S.E. 0.8%), suggesting that most of the inflation of association test statistic is explained by polygenicity and not PS. In comparison, GWAS of height without any adjustment for population stratification produce values of $R_{LDSC}$ ~10-13%.[26] Our estimated $R_{LDSC}$ is slightly smaller than that from LDSC analyses of previously published GWAS of height (Wood et al: 4.3% (S.E. 1.3%); Yengo et al.[18] 4.0% (S.E. 1.2%)), which suggests a reduced effect of residual PS on our results. Note that the estimated $R_{LDSC}$ shown here are obtained from analysing imputed GWAS summary statistics from Wood et al. and Yengo et al. (**Supplementary Methods**), which have a better coverage of HM3 SNPs; and thus explaining the difference with the ~6.0% (S.E. 1.0%) reported in Yengo et al. However, as a measure of PS, $R_{LDSC}$ is not strictly comparable across studies. In fact, the expectation of $R_{LDSC}$ (across repeated GWAS) not only depends on how much trait variance is explained by PS, but also on the degree of genetic differentiation between cohorts ($F_{ST}$) and the trait heritability within each cohort.[25] The last two factors, can vary from one GWAS meta-analysis to another as a function of cohort composition. In summary, these LDSC analyses suggest that uncorrected PS only marginally affects SNP effects from our large GWAS of height in EUR participants.

##### 1.2 Assessment of allele frequencies for height increasing alleles across the North-South axis of Europe reveals attenuated correlation relative to previous studies.

As an alternative to $R_{LDSC}$, we next quantified PS in our GWAS using the correlation between strength of association (p-value) and height-increasing allele frequency differences between Great Britain (GBR sample in the 1,000 Genomes Project – 1KGP) and the Italian Tuscan population (TSI sample in 1KGP). This statistic was previously introduced by Sohail and colleagues[24] to reveal biases in SNP effect estimates from the Wood et al. study, that were induced by uncorrected PS along the North-South gradient of Europe. More precisely, the strategy implemented by Sohail et al. consists in grouping SNPs based on strength of association, then regress the mean height-increasing allele frequency differences between GBR and TSI for each SNP bin onto the mean p-value of the corresponding bin. The slope of that linear regression ($\beta_{PS}$) measures the degree of PS. We estimated $\beta_{PS}$ using 101,360 near independent HM3 SNPs with MAF>1% and calculated standard errors using a bootstrap strategy based on 1,000 independent draws.

As previously reported, we found a significant $\beta_{PS}$ of ~1.28% (S.E. 0.08%; $P = 2.3 \times 10^{-61}$) using summary statistics of the Wood et al. study but no significant $\beta_{PS}$ from within-family GWAS in 17,942 independent UKB siblings pairs ($\beta_{PS}$=-0.08%, S.E. = 0.07%; $P$=0.27). We show in Suppl. Fig. 3a-b estimates of $\beta_{PS}$ from GWAS summary statistics of Yengo et al., EUR participants of 23andMe (23andME-EUR), all EUR participants of the UKB (UKB-456k; GWAS using BOLT-LMM), unrelated EUR participants of the UKB (UKB-350k; GWAS using PLINK), the meta-analysis EUR participants from multiple cohorts of the GIANT consortium (GIANT-EUR; N~1.6M); and the meta-analysis of GIANT-EUR and 23andMe-EUR. Overall, we find that $\beta_{PS}$ decreases with sample size, consistent with an increased signal-to-noise ratio. In particular, $\beta_{PS}$ is ~0.13% (P=0.1) in our largest GWAS meta-analysis of N~4.1M EUR participants, which demonstrates a better correction of PS than previously published EUR GWAS of height.

Furthermore, we assessed the correlation between estimated effects at these 101,360 independent HM3 SNPs and SNP loadings from 20 principal components (PCs) calculated in 503 EUR samples from 1KGP. Across SNPs, we found that SNP loadings on PC2 explain most of the variance in estimated SNP effects (Suppl. Fig. 3c). This observation is not surprising given that PC2 is the PC that correlates the most with the North-South axis of Europe, and therefore explains the consistency with our results based on $\beta_{PS}$. However, <0.3% of the variance of SNP effects estimated in our largest GWAS is explained by SNP loadings, which is much lower than ~2.3% obtained when analysing SNP effects from Wood et al.

**1.3 Comparison with estimated SNP effects from family-based GWAS**

Finally, we directly compared SNP effects from our GWAS ($\beta_{GWAS}$) with that of a within-family GWAS in 17,942 independent UKB siblings pairs ($\beta_{SIB}$). We used $\mathrm{S_{PS}} = \mathrm{cov}(\beta_{\mathrm{GWAS}}, \beta_{\mathrm{SIB}})/\mathrm{var}(\beta_{\mathrm{GWAS}})$ as our metric of interest in this comparison, where both $\mathrm{cov}(\beta_{\mathrm{GWAS}}, \beta_{\mathrm{SIB}})$ and $\mathrm{var}(\beta_{\mathrm{GWAS}})$ are calculated across SNPs. When SNP effects are estimated using ordinary least-squares (OLS) regression, the expectation of $\mathrm{S_{PS}}$ in the absence of PS is $E[\mathrm{S_{PS}}] = 1$. Therefore, a significant deviation of $S_{PS}$ below 1, may indicate confounding due to residual PS. However, the statistical properties of $\mathrm{S_{PS}}$ based on SNP effects estimated using linear mixed models (LMM) (or meta-analyses of OLS and LMM estimates) are not well characterised, which may affect our interpretation below.

We found that estimates of $S_{PS}$ based on SNPs strongly associated with height are much closer to 1 than when all SNPs are used, i.e. regardless of strength of association ($P$<1; Suppl. Fig. 3d). We observed the lowest value of $S_{PS}$~0.29 (S.E. 0.01) when using effects of all 101,360 SNPs estimated in the Wood et al. study. In comparison, estimated SNP effects from our largest EUR GWAS yields an $S_{PS}$>0.8 regardless of strength of association, yet still significantly lower than 1 ($P < 7 \times 10^{-30}$).

Lee et al.[27] previously showed that assortative mating (AM) on height can produce values of $S_{PS} < 1$. Under the assumption that the population has reached an equilibrium after many generations of AM with a constant spousal correlation ($r$), they showed that

(1.1) $$E[S_{PS}] = 1 - rh^2,$$

where $h^2$ is the full narrow-sense heritability in the current generation (at equilibrium). We note here an error in the Supplementary Notes of Lee et al., who used in their derivations the heritability in the base population undergoing random mating ($h_0^2$) instead of the equilibrium heritability, $h^2$. In practice, differences between $h_0^2 = (h^2 - rh^4)/(1 - rh^4)$ and $h^2$ are small. Therefore, using one or the other heritability has a limited impact on the expected value of $S_{PS}$. Using Equation (1.1) and assuming an equilibrium heritability, $h^2 = 0.8$ and a spousal correlation, $r = 0.25$, we expect $S_{PS}$ to be $\sim 1 - 0.8 \times 0.25 = 0.8$, which is consistent with our observations (Suppl. Fig. 3d).

Altogether, these analyses show that estimated SNP effects from our EUR GWAS are inflated by ~10-20% relative to that from a family-based GWAS and that this inflation is not larger than expected because of phenotypic AM on height.

**Supplementary Note 2: Distinguishing loss of tagging from multiplicity of causal variants at the *ACAN* locus**

We observed the largest density of independent associations around rs4932198, where 24 other GWS SNPs were detected within less than 100 kb on each side (Fig 1.; Suppl. Fig. 14). We hereafter refer to the set of these 25 GWS SNPs as *ACAN* GWAS signals. A large density of signals may reflect the presence of multiple causal variants or the presence of poorly tagged causal variants with large effects at this locus. To disentangle these two possible explanations, we first used statistically phased haplotypes from 346,959 unrelated UKB participants of EUR and tested their association with height. We analysed 14,117 haplotypes covering a 100 kb long genomic region at this locus (hg19 genomic coordinates: chr15:89,307,521-89,407,521) and that were present in at least 5 UKB participants. We tested the association between each haplotype and height but could not identify a single haplotype with a large enough effect that can explain the majority of signals at this locus (Suppl. Fig. 15). In fact, *ACAN* GWAS signals cumulatively explain ~0.3% of height variance, while the two most associated haplotypes ($P<10^{-7}$) jointly only explain 0.01% of height variance.

Next, we used genotypes at this locus from 291,683 unrelated EUR participants of the UKB to simulate a trait controlled by a single rare variant (MAF<1%) explaining between $\eta^2$=0.5% and 5% of variance, then performed a GCTA-COJO analysis to estimate the density of independent signals. Previous studies have shown that large discrepancies in sample size between discovery GWAS and LD reference may contribute to inflate the number of independent associations detected with GCTA-COJO.[28] Therefore, to mimic that effect we used a random subset of 10,000 unrelated EUR participants of the UKB as LD reference, i.e. ~1/30$^{th}$ of the discovery sample. On average over 100 simulation replicates for each value of $\eta^2$, we observed a signal density lower than 2.3 associations per 100kb (Suppl. Fig. 15). The largest density of 10 associations per 100 kb was observed only when the simulated causal variant explains 5% of trait variance (i.e. $\beta$ between ~2 and ~43 trait SD/allele), which is an extreme and unrealistic scenario. In contrast, even when the simulated causal variant explains 0.5% of trait variance, which in this case corresponds to a median allelic effect of ~1.4 SD/allele (~10cm, so very large) across simulations, we found that signal density never exceeded 7 associations per 100 kb. Altogether, the results of our haplotype- and simulation-based analyses suggest that a multiplicity of independent causal variants is the most likely explanation of our observations, although signal density is not an common estimator of the number of causal variants.

Finally, we sought to quantify how much the presence of a recently identified[29,30] height-associated variable-number-of-tandem-repeat (VNTR) polymorphism in *ACAN* may contribute to the observed density of GWS SNPs. Therefore, we regressed VNTR length imputed in UKB participants[28] onto allele counts at *ACAN* GWAS signals and found that these 25 SNPs explain ~73%, ~47%, ~42% and ~40% of VNTR length variation in SAS (N=9,219), EUR (N=414,429), AFR (N=7,543) and EAS (N=1,496) respectively (Suppl. Fig. 15e). Consistent with partial tagging of VNTR length variation, *ACAN* GWAS signals explain ~0.24% ($R^2_{VNTR}$ = 0.21% vs. $R^2_{VNTR+25\,SNPs}$ = 0.45%; P=$8.7 \times 10^{-55}$) additional height variance in EUR over what is explained by VNTR length variation alone (Suppl. Fig. 15f). In summary, these complementary analyses suggest that a large density of independent GWS SNPs near *ACAN* is partially explained by the presence of a VNTR at this locus and also by additional causal variants not yet identified.

**Supplementary Note 3: Optimal weighting of PGS and parental information to maximize prediction accuracy in the presence of assortative mating**

**Overview of theory, simulations and application to real data from the UK Biobank**

For a given individual, we denote y their phenotype, $y_m$ and $y_f$ the phenotypes of their mother and father respectively, $\bar{y}_p = (y_m + y_f)/2$ the average of their parents' phenotypes and $\hat{y}$ their own PGS. We consider combined predictor that is a linear combination of $\hat{y}$ and $\bar{y}_p$. Under the assumption that the resemblance between relatives is solely due to genetic factors, our main result is that the optimal weighting $\alpha_{PGS}\hat{y} + \alpha_{PA}\bar{y}_p$ is given by

$$\text{(S3.1)}\quad \alpha_{PGS} = \frac{R_{\hat{y},y}[1 - h^2(1 + r)/2]}{1 - R^2_{\hat{y},y}(1 + r)/2}$$

and

$$\text{(S3.2)}\quad \alpha_{PA} = \frac{h^2 - R^2_{\hat{y},y}}{1 - R^2_{\hat{y},y}(1 + r)/2}$$

where $h^2$ denotes the heritability of the trait in the current population, $r$ the correlation between spouses phenotypes in the population, and $R^2_{\hat{y},y} = \text{corr}(\hat{y}, y)^2$, the prediction accuracy of the PGS. The expected accuracy ($R^2_{\hat{y}+\bar{y}_p}$) of the combined predictor using these optimal weights, is given by

$$\text{(S3.3)}\quad R^2_{\hat{y}+\bar{y}_p} = \frac{R^2_{\hat{y},y} + \left(\frac{h^2}{2}\right)(1 + r)\left[h^2 - 2R^2_{\hat{y},y}\right]}{1 - R^2_{\hat{y},y}(1 + r)/2}$$

Suppl. Fig. 19 shows the results of simulations performed to verify the results from Equations (S3.1-S3.3). We define the regression weights as $\omega_{PGS} = \alpha_{PGS}/(\alpha_{PGS} + \alpha_{PA})$ and $\omega_{PA} = \alpha_{PA}/(\alpha_{PGS} + \alpha_{PA})$. Therefore, values of $\omega_{PGS}$ such that $\omega_{PGS} > 0.5$ imply that the PGS has a stronger weight that the parental average.

Under assortative mating, Suppl. Fig. 19 shows that $\omega_{PA}$ remains above 0.5 even when the PGS explains 50% of $h^2$. We show below that $\omega_{PGS} = \omega_{PGS}$ if

$$\text{(S3.4)}\quad R_{\hat{y},y} = \frac{-[1 - h^2(1 + r)/2] + \sqrt{[1 - h^2(1 + r)/2]^2 + 4h^2}}{2}$$

For example, with $r = 0.25$ and $h^2 = 0.8$, Equation (S3.4) predicts an equal contribution of the PGS and parental information is $R^2_{\hat{y},y} \approx 0.46$.

Next, we estimated $\alpha_{PGS}$, $\alpha_{PA}$ and $R^2_{\hat{y}+\bar{y}_p}$ in 981 trios from the UK Biobank (**Supplementary Methods**). For this analysis, we used a PGS based on 12,111 GWS SNPs identified in our largest GWAS meta-analysis. We found $\hat{\alpha}_{PGS}$~0.375 (S.E. = 0.025) and $\hat{\alpha}_{PA}$~0.634 (S.E. = 0.034). The variance explained by fitting both predictors is $\hat{R}^2_{\hat{y}+\bar{y}_p}$ = 0.542 (S.E. = 0.032), which is larger than the accuracy of each single predictor ($R^2_{\hat{y}} = 0.38$, S.E. 0.031; and $R^2_{\bar{y}_p} = 0.439$, S.E. 0.032). Next, we estimated the spousal correlation $\hat{r} = 0.233$ (S.E. = 0.031) and the heritability $\hat{h}^2 = 0.894$ (S.E. = 0.032) using mid-parent regression. Besides, the prediction accuracy of the PGS is $\hat{R}^2_{\hat{y},y}$~0.4. Therefore, from these estimates of r, $h^2$ and $R^2_{\hat{y},y}$ we predict using Equations (S3.1-S3.3) that $\alpha_{PGS} = 0.377$, $\alpha_{PA} = 0.656$ and $R^2_{\hat{y}+\bar{y}_p} =$ 0.599. These three predictions are not statistically distinct from estimated values, which further validates our model.

**Proof of theoretical results**

The linear combination of $\hat{y}$ and $\bar{y}_p$ that maximises prediction of y is derived from multivariate linear regression theory:

$$(3.1)\quad \alpha_{PGS}\hat{y} + \alpha_{PA}\bar{y}_p$$

where

$$(3.2)\quad \alpha_{PGS} = \frac{\mathrm{var}(\bar{y}_p)\mathrm{cov}(\hat{y},y) - \mathrm{cov}(\hat{y},\bar{y}_p)\mathrm{cov}(\bar{y}_p,y)}{\mathrm{var}(\bar{y}_p)\mathrm{var}(\hat{y}) - \mathrm{cov}(\hat{y},\bar{y}_p)^2} = \frac{\mathrm{var}(\bar{y}_p)\mathrm{var}(\hat{y})}{\mathrm{var}(\bar{y}_p)\mathrm{var}(\hat{y})} \times \frac{\mathrm{cov}(\hat{y},y)/\mathrm{var}(\hat{y}) - \dfrac{\mathrm{cov}(\hat{y},\bar{y}_p)\mathrm{cov}(\bar{y}_p,y)}{\mathrm{var}(\bar{y}_p)\mathrm{var}(\hat{y})}}{1 - [\mathrm{cov}(\hat{y},\bar{y}_p)^2]/[\mathrm{var}(\bar{y}_p)\mathrm{var}(\hat{y})]}$$

and

$$(3.3)\quad \alpha_{PA} = \frac{\mathrm{var}(\hat{y})\mathrm{cov}(\bar{y}_p,y) - \mathrm{cov}(\hat{y},\bar{y}_p)\mathrm{cov}(\hat{y},y)}{\mathrm{var}(\bar{y}_p)\mathrm{var}(\hat{y}) - \mathrm{cov}(\hat{y},\bar{y}_p)^2} = \frac{\mathrm{var}(\bar{y}_p)\mathrm{var}(\hat{y})}{\mathrm{var}(\bar{y}_p)\mathrm{var}(\hat{y})} \times \frac{\mathrm{cov}(\bar{y}_p,y)/\mathrm{var}(\bar{y}_p) - \dfrac{\mathrm{cov}(\hat{y},\bar{y}_p)\mathrm{cov}(\hat{y},y)}{\mathrm{var}(\bar{y}_p)\mathrm{var}(\hat{y})}}{1 - [\mathrm{cov}(\hat{y},\bar{y}_p)^2]/[\mathrm{var}(\bar{y}_p)\mathrm{var}(\hat{y})]}$$

Without loss of generality, we assume that y and $\hat{y}$ are both centred (i.e. $E[y] = E[\hat{y}] = 0$) and scaled (i.e. $\mathrm{var}[y] = \mathrm{var}[\hat{y}] = 0$). Assuming that $\mathrm{var}[y] = 1$, implies that $\mathrm{var}[\bar{y}_p] = (1 + r)/2$, where r denotes the phenotypic correlation between mates in the population.

We also denote $R_{\hat{y},y} = \mathrm{corr}(\hat{y},y) = \mathrm{cov}(\hat{y},y)$, $R_{\bar{y}_p,y} = \mathrm{corr}(\bar{y}_p,y)$ and $R_{\hat{y},\bar{y}_p} = \mathrm{corr}(\bar{y}_p,y)$. Therefore, Equations (3.2) and (3.3) simplify as

$$(3.2')\quad \alpha_{PGS} = \frac{\mathrm{cov}(\hat{y},y)/\mathrm{var}(\hat{y}) - \dfrac{\mathrm{cov}(\hat{y},\bar{y}_p)}{\sqrt{\mathrm{var}(\bar{y}_p)\mathrm{var}(\hat{y})}} \times \dfrac{\mathrm{cov}(\bar{y}_p,y)}{\sqrt{\mathrm{var}(\bar{y}_p)\mathrm{var}(y)}} \times \sqrt{\dfrac{\mathrm{var}(y)}{\mathrm{var}(\hat{y})}}}{1 - R^2_{\hat{y},\bar{y}_p}} = \frac{R_{\hat{y},y} - R_{\hat{y},\bar{y}_p}R_{\bar{y}_p,y}}{1 - R^2_{\hat{y},\bar{y}_p}}$$

and

$$(3.3')\quad \alpha_{PA} = \frac{\dfrac{\mathrm{cov}(\bar{y}_p,y)}{\sqrt{\mathrm{var}(\bar{y}_p)\mathrm{var}(y)}} \times \sqrt{\dfrac{\mathrm{var}(y)}{\mathrm{var}(\bar{y}_p)}} - \dfrac{\mathrm{cov}(\hat{y},\bar{y}_p)}{\sqrt{\mathrm{var}(\bar{y}_p)\mathrm{var}(\hat{y})}} \times \dfrac{\mathrm{cov}(\hat{y},y)}{\sqrt{\mathrm{var}(\hat{y})\mathrm{var}(y)}} \times \sqrt{\dfrac{\mathrm{var}(y)}{\mathrm{var}(\bar{y}_p)}}}{1 - R^2_{\hat{y},\bar{y}_p}}$$

$$= \sqrt{\frac{2}{1+r}}\left(\frac{R_{\bar{y}_p,y} - R_{\hat{y},\bar{y}_p}R_{\hat{y},y}}{1 - R^2_{\hat{y},\bar{y}_p}}\right)$$

and further as

$$(3.2'')\quad \alpha_{PGS} = \frac{R_{\hat{y},y} - R_{\hat{y},\bar{y}_p}R_{\bar{y}_p,y}}{1 - R^2_{\hat{y},\bar{y}_p}}$$

and

$$(3.3'')\quad \alpha_{PA} = \sqrt{\frac{2}{1+r}}\left(\frac{R_{\bar{y}_p,y} - R_{\hat{y},\bar{y}_p}R_{\hat{y},y}}{1 - R^2_{\hat{y},\bar{y}_p}}\right)$$

Equations (3.2'') and (3.3'') are expressed in function of $R_{\hat{y},y}$, $R_{\bar{y}_p,y}$, $R_{\hat{y},\bar{y}_p}$ and r.

We assume that $R_{\hat{y},y}$ is known, e.g., from quantifying the accuracy of the PGS in some validation sample. If we denote $h^2$ as heritability in the current population (which could be undergoing assortative mating, i.e. $r \neq 0$) and assume no shared environmental effects between parents and offspring then

$$(3.4) \quad h^2 = cov(\bar{y}_p, y)/var(\bar{y}_p) \Rightarrow R_{\bar{y}_p,y} = h^2 \sqrt{\frac{1+r}{2}}$$

Finally, denote $\hat{y}_p$ as the average PGS of parents. We can express $\bar{y}_p$ as a function of $\hat{y}_p$ as follows

$$(3.5) \quad \bar{y}_p = \frac{cov(\bar{y}_p, \hat{y}_p)}{var(\hat{y}_p)} \hat{y}_p + \varepsilon_p$$

where $\varepsilon_p$ is a residual term with mean 0 and such that $cov(\varepsilon_p, \hat{y}_p) = 0$. Given that both phenotypes and PGS are centred, $cov(\bar{y}_p, \hat{y}_p)$ can be expressed as

$$cov(\bar{y}_p, \hat{y}_p) = \frac{1}{4} E[y_m \hat{y}_m + y_f \hat{y}_f + y_f \hat{y}_m + y_m \hat{y}_f] = \frac{1}{2} E[R_{\hat{y},y} + E[y_f \hat{y}_m | y_m]] = R_{\hat{y},y}(1+r)/2$$

Using a similar reasoning, we can show that $var(\hat{y}_p) = (1 + rR^2_{\hat{y},y})/2$. Therefore, Equation (3.5) can be rewritten as

$$(3.5') \quad \bar{y}_p = R_{\hat{y},y} \left( \frac{1+r}{1 + rR^2_{\hat{y},y}} \right) \hat{y}_p + \varepsilon_p$$

Besides, we can also write

$$(3.6) \quad \hat{y} = \hat{y}_p + \varepsilon_m,$$

where $\varepsilon_m$ (Mendelian segregation) is independent of $\hat{y}_p$. Combining Equations (3.5') and (3.6) leads to

$$(3.7) \quad cov(\bar{y}_p, \hat{y}) = R_{\hat{y},y} \left( \frac{1+r}{1 + rR^2_{\hat{y},y}} \right) cov(\hat{y}_p, \hat{y}) + cov(\hat{y}, \varepsilon_p) = R_{\hat{y},y} \left( \frac{1+r}{1 + rR^2_{\hat{y},y}} \right) var(\hat{y}_p) + cov(\varepsilon_m, \varepsilon_p)$$

which, assuming $cov(\varepsilon_m, \varepsilon_p) = 0$, implies that $cov(\bar{y}_p, \hat{y}) = R_{\hat{y},y} \left( \frac{1+r}{1 + rR^2_{\hat{y},y}} \right) var(\hat{y}_p) = R_{\hat{y},y}(1+r)/2$.

Therefore,

$$(3.8) \quad R_{\hat{y},\bar{y}_p} = corr(\bar{y}_p, \hat{y}) = \frac{R_{\hat{y},y}(1 + r)}{\sqrt{2(1+r)}}.$$

It follows that

$$R^2_{\hat{y},\bar{y}_p} = \frac{R^2_{\hat{y},y}(1 + r)^2}{2(1+r)} = R^2_{\hat{y},y}(1+r)/2 \Leftrightarrow \frac{1}{1 - R^2_{\hat{y},\bar{y}_p}} = \frac{1}{1 - R^2_{\hat{y},y}(1+r)/2}$$

We now express below $\alpha_{PGS}$ and $\alpha_{PA}$ as a function of $h^2$, $R_{\hat{y},y}$, and r.

We first recall that $\alpha_{PGS} = \frac{R_{\hat{y},y} - R_{\hat{y},\bar{y}_p} R_{\bar{y}_p,y}}{1 - R^2_{\hat{y},\bar{y}_p}}$

$$R_{\hat{y},\bar{y}_p}R_{\bar{y}_p,y} = \frac{R_{\hat{y},y}(1 + r)}{\sqrt{2(1+r)}} \times h^2\sqrt{\frac{1+r}{2}} = R_{\hat{y},y}h^2(1 + r)/2.$$

$$R_{\hat{y},y} - R_{\hat{y},\bar{y}_p}R_{\bar{y}_p,y} = R_{\hat{y},y}[1 - h^2(1 + r)/2].$$

To calculate $\alpha_{PA} = \sqrt{\frac{2}{1+r}}\left(\frac{R_{\bar{y}_p,y} - R_{\hat{y},\bar{y}_p}R_{\hat{y},y}}{1 - R^2_{\hat{y},\bar{y}_p}}\right)$

$$\sqrt{\frac{2}{1+r}}\left(R_{\bar{y}_p,y} - R_{\hat{y},\bar{y}_p}R_{\hat{y},y}\right) = \sqrt{\frac{2}{1+r}}\left[h^2\sqrt{\frac{1+r}{2}} - \frac{R^2_{\hat{y},y}(1 + r)}{\sqrt{2(1+r)}}\right] = h^2 - \frac{R^2_{\hat{y},y}(1 + r)}{1+r} = h^2 - R^2_{\hat{y},y}$$

Finally,

$$(3.2''')\quad \alpha_{PGS} = \frac{R_{\hat{y},y} - R_{\hat{y},\bar{y}_p}R_{\bar{y}_p,y}}{1 - R^2_{\hat{y},\bar{y}_p}} = \frac{R_{\hat{y},y}[1 - h^2(1 + r)/2]}{1 - R^2_{\hat{y},y}(1+r)/2}$$

and

$$(3.3''')\quad \alpha_{PA} = \sqrt{\frac{2}{1+r}}\left(\frac{R_{\bar{y}_p,y} - R_{\hat{y},\bar{y}_p}R_{\hat{y},y}}{1 - R^2_{\hat{y},\bar{y}_p}}\right) = \frac{h^2 - R^2_{\hat{y},y}}{1 - R^2_{\hat{y},y}(1+r)/2}$$

Special case: $r = 0$

$$\alpha_{PGS} = \frac{R_{\hat{y},y}[1 - h^2/2]}{1 - R^2_{\hat{y},y}/2} \quad \text{and } \alpha_{PA} = \frac{h^2 - R^2_{\hat{y},y}}{1 - R^2_{\hat{y},y}/2}$$

The relative contribution of $\hat{y}$ and $\bar{y}_p$, defined above as $\omega_{PGS} = \alpha_{PGS}/(\alpha_{PGS} + \alpha_{PA})$ and $\omega_{PA} = \alpha_{PA}/(\alpha_{PGS} + \alpha_{PA})$ can be expressed as

$$\omega_{PGS} = \frac{R_{\hat{y},y}[1 - h^2(1 + r)/2]}{R_{\hat{y},y}[1 - h^2(1 + r)/2] + h^2 - R^2_{\hat{y},y}} \quad \text{and } \omega_{PA} = \frac{h^2 - R^2_{\hat{y},y}}{R_{\hat{y},y}[1 - h^2(1 + r)/2] + h^2 - R^2_{\hat{y},y}}$$

These two relative contributions are equal when $\omega_{PA} = \omega_{PGS} = 1/2$, i.e. when

$$2h^2 - 2R^2_{\hat{y},y} = R_{\hat{y},y}[1 - h^2(1 + r)/2] + h^2 - R^2_{\hat{y},y} \Leftrightarrow R^2_{\hat{y},y} + R_{\hat{y},y}[1 - h^2(1 + r)/2] - h^2$$

or equivalently, when

$$R_{\hat{y},y} = \frac{-[1 - h^2(1 + r)/2] + \sqrt{[1 - h^2(1 + r)/2]^2 + 4h^2}}{2}$$

This therefore proves Equation (S3.4).

**Prediction accuracy from a linear regression model fitting both PGS and parental average**

The expected prediction accuracy ($R^2_{\hat{y}+\bar{y}_p}$) from combining PGS and parental information can be expressed as

$$R^2_{\hat{y}+\bar{y}_p} = \alpha_{PGS}\text{cov}(\hat{y}, y) + \alpha_{PA}\text{cov}(\bar{y}_p, y) = \frac{R^2_{\hat{y},y}[1 - h^2(1+r)/2] + h^2(h^2 - R^2_{\hat{y},y})(1+r)/2}{1 - R^2_{\hat{y},y}(1+r)/2}$$

which can be simplified as

(3.9) $$R^2_{\hat{y}+\bar{y}_p} = \frac{R^2_{\hat{y},y}+\left(\frac{h^2}{2}\right)(1+r)\left[h^2-2R^2_{\hat{y},y}\right]}{1-R^2_{\hat{y},y}(1+r)/2}$$

**Prediction accuracy and proportion of causal variants captured**
We assume that the trait of interest in underlain by M independent causal SNPs and that m ($m \leq M$) of them are included in a PGS. Moreover, we assume that the population has been undergoing assortative mating for multiple generations, until an equilibrium is reached. We derive below how large m needs to be for the prediction accuracy of the derived PGS, in the equilibrium population, to equal $R^2_{\hat{y},y}$.

We denote $\rho = rh^2$, $f_0 = m/M$, $\gamma = \rho/(1-\rho)$, $\alpha = \gamma/(2M)$ the expected correlation between trait-increasing alleles induced by assortative mating, and $\sigma^2_{g,0}$ and $\sigma^2_{g,eq}$ the genetic variances in a randomly and assortatively mating populations, respectively.

In the equilibrium population, the variance of the PGS can be expressed as

(Int. 3.1) $\mathrm{var}(\hat{y}) \approx \sigma^2_{g,0} f_0(1+\gamma f_0)$, and the covariance between y and $\hat{y}$ as

(Int. 3.2) $\mathrm{cov}(\hat{y}, y) \approx \sigma^2_{g,0} f_0(1+\gamma)$.

Equations (Int. 3.1) and (Int. 3.2) are proven below.

Therefore,

(3.10) $$R^2_{\hat{y},y} = \frac{\mathrm{cov}(\hat{y},y)^2}{\mathrm{var}(\hat{y})\mathrm{var}(y)} \approx \sigma^2_{g,0} f_0 \frac{(1+\gamma)^2}{1+\gamma f_0}$$

Diving the previous equation by $\sigma^2_{g,eq}$ leads to and $\phi_{eq} = R^2_{\hat{y},y}/\sigma^2_{g,eq}$.
Therefore, Equation (3.10) implies that

(3.11) $$(1+\gamma f_0)\phi_{eq} \approx \left(\frac{\sigma^2_{g,0}}{\sigma^2_{g,eq}}\right) f_0(1+\gamma)^2 = f_0(1-\rho)(1+\gamma)^2 \Leftrightarrow f_0 \approx \frac{\phi_{eq}}{(1-\rho)(1+\gamma)^2 - \gamma\phi_{eq}}$$

where $\phi_{eq} = R^2_{\hat{y},y}/\sigma^2_{g,eq}$. Note that $\sigma^2_{g,eq}/\sigma^2_{g,0}$ is the inflation in genetic variance due to assortative mating, which is predicted in theory to equal $1/(1-\rho)$.

Using a similar reasoning, Yengo et al.[31] (Eq. 1.20 in their Supplementary Note) derived the relationship between $f_0$ and the proportion $f_{eq} = h^2_{SNP}/h^2$ of equilibrium heritability explained by the m SNPs included in the PGS as:

(3.12) $$f_0 = \frac{1-\rho}{2\rho}\left[\sqrt{\left(1+\frac{4f_{eq}\rho}{(1-\rho)^2}\right)} - 1\right] \underset{|\rho|\ll 1}{\approx} f_{eq}/(1-\rho).$$

**Proof of Equation (Int. 3.1) and (Int. 3.2)**
We assume an infinitesimal model, where each causal SNP explains the same amount of trait variance. For simplicity, we assume the squared effect size of each causal SNP to equal $b^2 = \sigma^2_{g,0}/M$; and that SNP effects are estimated with negligible errors so that they could be assumed to be equal to their true value. Finally, we assume that the m first SNPs are included in the PGS.

Under these assumptions, the PGS (i.e. ŷ) can be written as

$$\hat{y} = \left(\frac{\sigma_{g,0}^2}{M}\right)\sum_{j=1}^{m} z_j$$

where $(z_j - 2p_j)/\sqrt{2p_j(1-p_j)}$ is the centred and scaled count of trait-increasing allele at SNP j and $p_j$ the trait-increasing allele frequency at that same SNP. By definition, we have that $\text{var}(z_j) = 1 + \alpha$, and that $\text{cov}(z_j, z_k) = 2\alpha$. It follows that

$$\text{var}(\hat{y}) = \frac{\sigma_{g,0}^2}{M}[m(1+\alpha) + m(m-1)2\alpha] = \sigma_{g,0}^2\left(\frac{m}{M}\right)[1 + (2m-1)\alpha] = \sigma_{g,0}^2 f_0\left[1 + \left(f_0 - \frac{1}{2M}\right)(2M\alpha)\right]$$

For large values of $M$, this simplifies as $\text{var}(\hat{y}) \approx \sigma_{g,0}^2 f_0(1 + \gamma f_0)$.

Similarly, we can write $\text{cov}(\hat{y}, y)$ as $\text{cov}(\hat{y}, y) = \frac{\sigma_{g,0}^2}{M}[m(1+\alpha) + m(M-1)2\alpha] \approx \sigma_{g,0}^2 f_0(1+\gamma)$.

### Supplementary Note 4: Saturation of GWAS signals within pathways and gene sets

#### Overview and main results

We assessed the enrichment of broad categories of biological pathways for different GWAS sample sizes, using two different gene sets enrichment methods, DEPICT[32] and MAGMA.[33] Besides, we evaluated the prioritization of 14,462 gene sets, hierarchically clustered into 20 groups of related gene sets based on gene set membership (see Methods below, Suppl. Figs. 20 - 21, Suppl. Table 13). We observed an enrichment of OMIM genes in clusters 1, 2, 5, 6, 11, 16, and 17 (Bonferroni P < 0.05 vs. random genes (Suppl. Fig. 22, Suppl. Table 14). At all sample sizes tested (range N=130,010 to N=5,314,291), similar sets of the clusters consistently showed significant enrichments in DEPICT (clusters 2, 5, 11, 16, and 17) and MAGMA-prioritized gene sets (clusters 5, 11, 16, and 17; Suppl. Fig. 21). Thus, the broad patterns of gene set enrichment are apparent even at moderate sample sizes and remain quite stable as sample sizes increase.

In contrast with clusters of gene sets, individual genes may require larger sample sizes or multiple ancestries to be implicated by GWAS. To address these questions, we assessed the fraction of OMIM genes that contain an approximately independent genome-wide significant signal (identified with COJO) across the range of GWAS sample sizes. As sample size increases and the number of independent signals increases, the percent of the 462 OMIM genes overlapping a signal also increases (Suppl. Fig. 23b); however, after subtracting the null background from randomly sampled sets of 462 genes, the percentage above background of OMIM genes that overlap GWAS signal plateaus at a sample size of ~2.5 M (Suppl. Fig. 23c). In comparing the trans-ancestry meta-analysis with the largest European-ancestry-only GWAS with, we did not observe a noticeable increase in overlapping OMIM genes above background.

We also sought to examine more directly whether the height GWAS results implicate highly similar biology across different continental ancestries. We used MAGMA and DEPICT to prioritize genes based on GWAS results for EUR, EAS, and AA ancestries. We then compared the enrichment of heritability with stratified LD score regression (LDSC)[12,34] for each set of prioritized genes, evaluated either in the same ancestry or in the other two ancestries. Genes prioritized in one ancestry by both MAGMA and DEPICT showed comparable enrichment of heritability when evaluated either in that ancestry or in the other two ancestries (Suppl. Fig. 24, Suppl. Table 15), strongly confirming the shared biology implicated by GWAS results from different ancestries.

#### Methods

<u>Evaluation of gene set enrichment analysis (GSEA) methods across sample sizes.</u>

For GWAS summary statistics from multiple sample sizes (Tables 1 - 2) two GSEA approaches were applied (DEPICT and MAGMA). DEPICT release 173 was used; the top 1000 SNPs pruned by p-value from each set of summary statistics were used as input for each sample. MAGMA version v1.07b was used; SNPs were annotated with genes within 100kb, and genes were removed if the missingness of their pathway membership was over 0.2.

To evaluate the ability of GSEA methods to identify groups of gene sets at different sample sizes, 14,462 gene sets, each consisting of Z-scores for 19,987 genes (selected in ref.[32]), were hierarchically clustered into 20 clusters as follows. Pairwise distances between gene sets were defined as the Euclidean distance between the gene sets' Z-scores and the elbow method was used to choose the number of clusters, evaluating average distances between cluster centroids as the number of clusters is varied (Suppl. Fig. 20). For DEPICT and MAGMA, enrichment of prioritization in each cluster was defined as the number of prioritized gene sets in each cluster divided by the size of each cluster; a gene set was considered prioritized if it was in the top 10% of gene sets as prioritized by the GSEA method. Enrichment of OMIM genes in each gene set was defined as the number of OMIM genes in each gene set divided by the size of each gene set divided by the proportion of all genes in OMIM, and then enrichment of OMIM genes in each cluster was defined as the average of the enrichment of OMIM genes in each gene set in that cluster.

Genes were defined to be "in a gene set" if the gene's gene-set Z-score is > 1.96, as described previously.[35] Null distributions for each cluster were generated by randomly selected prioritized gene sets (for DEPICT and MAGMA) or prioritized genes to evaluate enrichment significance.

To evaluate saturation of height-associated gene identification, the percentage of OMIM genes overlapping independent COJO signals was calculated. "Overlapping" was defined as having at least one COJO SNP within the gene body, as defined with the plink version 1.9 hg19 gene list (URL: https://www.cog-genomics.org/static/bin/plink/glist-hg19). A null distribution was calculated by drawing an equivalent number of random genes (binned by size into 20 bins, same number of genes per bin) to match OMIM genes, and calculating the percent of the random genes near a COJO SNP.

Benchmarking of gene prioritization across different ancestries

We applied DEPICT and MAGMA to prioritize genes on height GWAS of European, African-American and East Asian ancestry, resulting in three sets of prioritized genes for each method. To allow for a fair comparison, we used subsets of the available cohorts to create three equally sized GWAS (N~100,000). For MAGMA, we converted gene set prioritizations to gene prioritizations as described previously.[35] For both DEPICT and MAGMA, we then used Benchmarker[35] to evaluate the performance of these three sets of genes in each of the three different ancestries, resulting in three within-ancestry and six cross-ancestry scenarios.

The Benchmarker method is based on a leave-one-chromosome-out approach where one chromosome is withheld, and GWAS results for the remaining 21 chromosomes are used to prioritize genes on the withheld chromosome, iterating across each withheld chromosome. For each of the discovery GWAS ancestries, we selected the top 10% of the prioritized genes on each left out chromosome, resulting in 1,893 prioritized genes. We subsequently annotated SNPs within 50kb of the prioritized genes to generate a LD score annotation for these SNPs using LDSC.[25] Lastly, we applied stratified LDSC[34] (S-LDSC) to compare the three annotation sets to the full GWAS results for each of the three ancestries to determine whether the performance of genes prioritized and then evaluated across the same ancestry would be more enriched for heritability compared with those prioritized and evaluated in different ancestries. Reference panels were based on the 1000 Genomes Phase 3 reference panels[36] for LD score estimation, matching the reference panel ancestry with the GWAS results for that same ancestry. In addition, a category of SNPs that locate within 50kb of any gene in the prioritization method and a set of 53 annotations of known genomic importance were included in the S-LDSC as conditional covariates. The analysis was based on 1,217,311 HapMap3 SNPs. The results of the S-LDSC is summarized using proportional $h^2_{\mathrm{SNP}}$ (proportion of heritability explained by the annotation), the regression coefficient (average per-SNP contribution of the annotation to heritability), and enrichment in heritability ($h^2$ divided by the proportion of SNPs in the annotation). To access the performance difference between two annotations, we calculated p-values based on standard errors from the different estimates.
